# Autophagy regulator ATG5 preserves cerebellar function by safeguarding its glycolytic activity

**DOI:** 10.1101/2024.01.27.577372

**Authors:** J. Tutas, M. Tolve, E. Özer-Yildiz, L. Ickert, I Klein, M Hosseini, F. Liebsch, F. Dethloff, P. Gavalisco, H. Endepols, M. Yang, T. Georgomanolis, B. Neumaier, A. Drzezga, S. Daun, G. Schwarz, B. Thorens, G. Gatto, C. Frezza, NL Kononenko

## Abstract

Dysfunctions in autophagy, a highly conserved cellular mechanism responsible for the degradation of intracellular components within lysosomes, often result in neurodegeneration. The neuroprotective effect of autophagy varies across neuronal subtypes, and the mechanisms of selective vulnerability of neurons to autophagy dysfunction are currently unknown. Utilizing a mouse model of ATG5 deficiency in inhibitory neurons and a comprehensive approach, including PET imaging, metabolomics, stable-isotope labeling studies, and live cell imaging, we establish that autophagy contributes to the survival of cerebellar Purkinje cells (PCs) by safeguarding their glycolytic activity. We show that the core autophagy protein ATG5 downregulates the levels of the glucose transporter 2 (GLUT2) during brain maturation. Autophagy-deficient PCs exhibit increased glucose uptake, along with elevated levels of glycolytic intermediates and methylglyoxal-modified proteins. We propose lysophosphatidic acid and serine as glycolytic intermediates inducing PC death and demonstrate that deletion of GLUT2 in ATG5-deficient mice mitigates PC neurodegeneration and restores their ataxic gait. Taken together, this study reveals a novel neuroprotective role of autophagy in preventing excessive glycolytic metabolism in the brain.

## Introduction

Autophagy is a highly conserved lysosomal pathway that removes and recycles damaged or unnecessary cellular components^1^. The most common form of autophagy is macroautophagy (hereafter autophagy). During this process, parts of the cytoplasm together with the cargo destined for degradation are enclosed in double-membrane vesicles, the so-called autophagosomes, which are passed on to the lysosomes for degradation. There are two ubiquitin-like conjugating systems necessary for the elongation and closure of the autophagosome membrane, namely ATG12-ATG5-ATG16L1 and LC3-I/LC3-II ^2, 3^. This step is essential for autophagy progression, and the absence of components of the ubiquitin-like conjugating system such as ATG5 leads to a functional inhibition of the autophagy machinery ^4^. In recent years, there has been a particular emphasis on exploring the role of autophagy in the brain, driven by its association with neurodegeneration ^5-9^. Deficiency in *Atg5* or *Atg7* in neuronal progenitors leads to cell loss associated with accumulation of aggregated and ubiquitinated proteins ^8, 9^. However, the question of whether protein deposits or impaired autophagy alone triggers neurodegeneration, and the specific mechanisms by which this occurs, remains unanswered.

Autophagy is constitutively active at low levels and acts as a quality control mechanism. Its recognized pivotal role in cell survival stems from the selective clearance of toxic proteins and dysfunctional organelles ^10^. However, autophagy can extend its impact on cell survival beyond its conventional housekeeping function by actively regulating cellular metabolism ^11, 12^. In a nutrient-deprived environment, autophagy enables metabolic adaptation across species by ensuring the availability of survival-critical biomolecules^13-16^. Newborn mice require autophagy to sustain serum amino acid levels during neonatal starvation ^17, 18^. In adult mice, autophagy serves to maintain serum glucose levels during fasting ^19^, and its disruption has been associated with insulin resistance ^20, 21^. Starvation-induced autophagy also contributes free fatty acids for energy supply by degrading lipid droplets ^22^, although the role of autophagy in lipid droplet biogenesis under nutrient deprivation has also been reported^23^. Autophagy can also regulate metabolism independently of starvation, and this function has received particular attention in tumor cells ^24, 25^. Autophagy-deficient cancer cells exhibit increased glycolytic metabolism even under nutrient-rich conditions ^26^, and can undergo metabolic rewiring by metabolizing more glucose to lactate through aerobic glycolysis in the presence of oxygen ^27^. This metabolic switch is required for cancer cells to grow by biosynthesis, and may be achieved in part through the role of macroautophagy and chaperone-mediated autophagy in the recycling and trafficking of key proteins in the glycolytic pathway ^28-31^. One of the products of accelerated glycolysis is methylglyoxal (MG), a reactive dicarbonyl with the potential to glycate proteins and lipids ^32^. Increased MG production leads to carbonyl stress, a recognized hallmark of aging ^33, 34^. Elevated levels of carbonyl proteins have been detected in cerebrospinal fluid and plasma of patients with age-associated neurodegenerative diseases, including Parkinson’s (PD) and Alzheimer’s diseases (AD) ^35, 36^. Whether these carbonyl proteins originate from accelerated aerobic glycolysis in neurons, a pathway traditionally considered neuroprotective^37, 38^, is currently unclear. Intriguingly, a metabolic switch to aerobic glycolysis has been recently shown to underlie the neurodegeneration in sporadic AD patient-derived neurons ^39^. Autophagy is also compromised during aging ^40^, but whether autophagy can serve a neuroprotective role by safeguarding glycolytic metabolism in the brain is yet to be determined.

The most well-known function of neuronal autophagy in preserving energy metabolism is its involvement in the degradation of damaged and/or old mitochondria, a process known as mitophagy ^41, 42^. However, our understanding of the housekeeping-independent role of autophagy in brain metabolism is still limited when compared to non-neuronal cells. Autophagy in the hypothalamus is involved in mobilizing lipids for fatty acid supply and regulating lipolysis and glucose metabolism ^43, 44^. Autophagy-deficient microglia accumulate lipid droplets and exhibit a reduction in fatty acid β-oxidation, a major lipid consuming pathway ^45^. Recently, we found that the crucial autophagy modifier ATG5 operates in cortical synapses to maintain functional cAMP/PKA signaling, but is not essential for the survival of excitatory cortical ^46^ and/or inhibitory cortical and striatal neurons ^47^, aligning with findings from several other studies ^48, 49^. This stands in stark contrast to the progressive loss of Purkinje cells (PC) in the cerebellum observed in mice upon the deletion of ATG5 ^50^. The mechanism behind the selective vulnerability of PCs under conditions of defective autophagy remains currently elusive.

Here, we find that the core autophagy protein ATG5 regulates PCs survival independently of its conventional role in clearing protein aggregates and dysfunctional mitochondria. Instead, ATG5 acts neuroprotective by controlling the excessive glycolytic activity of PCs. Utilizing a mouse model with targeted ATG5 deficiency in inhibitory neurons and employing a comprehensive approach involving positron emission tomography (PET) imaging, quantitative proteomics and metabolomics, stable isotope-labeling studies, live-imaging, and in-depth kinematic analysis, we demonstrate that autophagy functions in PCs to downregulate the levels of glucose transporter 2 (GLUT2) during brain maturation. Autophagy-deficient PCs accumulate GLUT2 on their plasma membrane, a phenotype accompanied by an increase in glucose uptake, upregulation of glycolytic intermediates, and facilitated flux of glucose into the glycolysis pathway. The excessive glycolysis in ATG5 KO PCs results in increased non-mitochondrial ATP, elevated lactate production and increased levels of methylglyoxal-modified proteins. We reveal that several products of uncontrolled glycolysis, including serine and lysophosphatidic acid (LPA), are toxic to PCs and that levels of these intermediates are elevated in autophagy-deficient cerebellum. Remarkably, deletion of GLUT2 in ATG5 KO mice mitigates PC neurodegeneration and improves their ataxic gait. Our results demonstrate for the first time that the neuroprotective functions of autophagy in the brain include its role in preventing excessive glycolytic metabolism.

## Results

### Differential vulnerability of inhibitory cortical and cerebellar neurons to ATG5 loss

To understand how precisely autophagy contributes to the degeneration of PCs, we capitalized on the previously published mouse line lacking the crucial autophagy component ATG5 in inhibitory neurons (*Atg5*flox:flox/*Slc32a1-*Cre^tg^ KO mice, further defined as ATG5 cKO mice)^47^. In these mice, the inhibition of autophagy is reflected in significantly increased protein levels of autophagy receptor p62 and downregulated LC3 levels at autophagosomal membranes in the cortex (Fig. S1a-c), as well as in the cerebellum (Fig. S1d-f), which occurs as early as one month of age. We previously observed no loss of forebrain GABAergic neurons in 3-month-old ATG5 cKO mice, a phenotype that was consistent until 10 months of age ^47^. To perform an unbiased analysis of brain-wide atrophy in ATG5 cKO mice in-vivo, we performed longitudinal PET imaging, using the [^18^F]-FDG-(Fluorodeoxyglucose) radiotracer, which is used as a downstream marker of neuronal activity and an indirect reporter of neurodegeneration. In agreement with our previous work, in which no neurodegeneration of autophagy-deficient cortical and striatal GABAergic neurons was detected ^47^, we found only minor changes in the cortical and/or striatal FDG-PET signal in cKO mice compared to control mice (Fig. 1a). In contrast, a strong decrease in FDG uptake was detected in the cKO cerebellum, suggesting a decrease in cerebellar neuronal activity in mice with autophagy deficiency. These changes already occurred at the age of 3 months and persisted until the age of 12 months. To determine whether this decrease in FDG-PET signal is related to neurodegeneration, we measured changes in synapse density using the PET radiotracers [^18^F]-UCB-H and/or [^18^F]-MNI1126, which both bind to synaptic vesicle glycoprotein 2A (SV2A) (Fig. 1b). The SV2A-PET signal was significantly reduced in the cerebellum and cortex of 3- and 12-month-old ATG5 cKO animals compared to their control littermates (i.e., *Atg5*wt/wt: *Slc32a1-*Cre^tg^ mice, further defined as WT mice). However, the cerebellum was the only brain region that showed a positive correlation of the SV2A-PET signal with [^18^F]-FDG-PET in ATG cKO mice (Fig. 1c), suggesting that the reduced glucose metabolism is at least partly due to the loss of neurons and/or their synapses in this region. In agreement with this finding, we detected no alterations in GFAP levels (a marker of astrogliosis) in the ATG5 cKO cortex (Fig. S1g), whereas GFAP was significantly increased in the cerebellum of 3-month-old autophagy-deficient mice (Fig. S1h).

**Figure 1.**
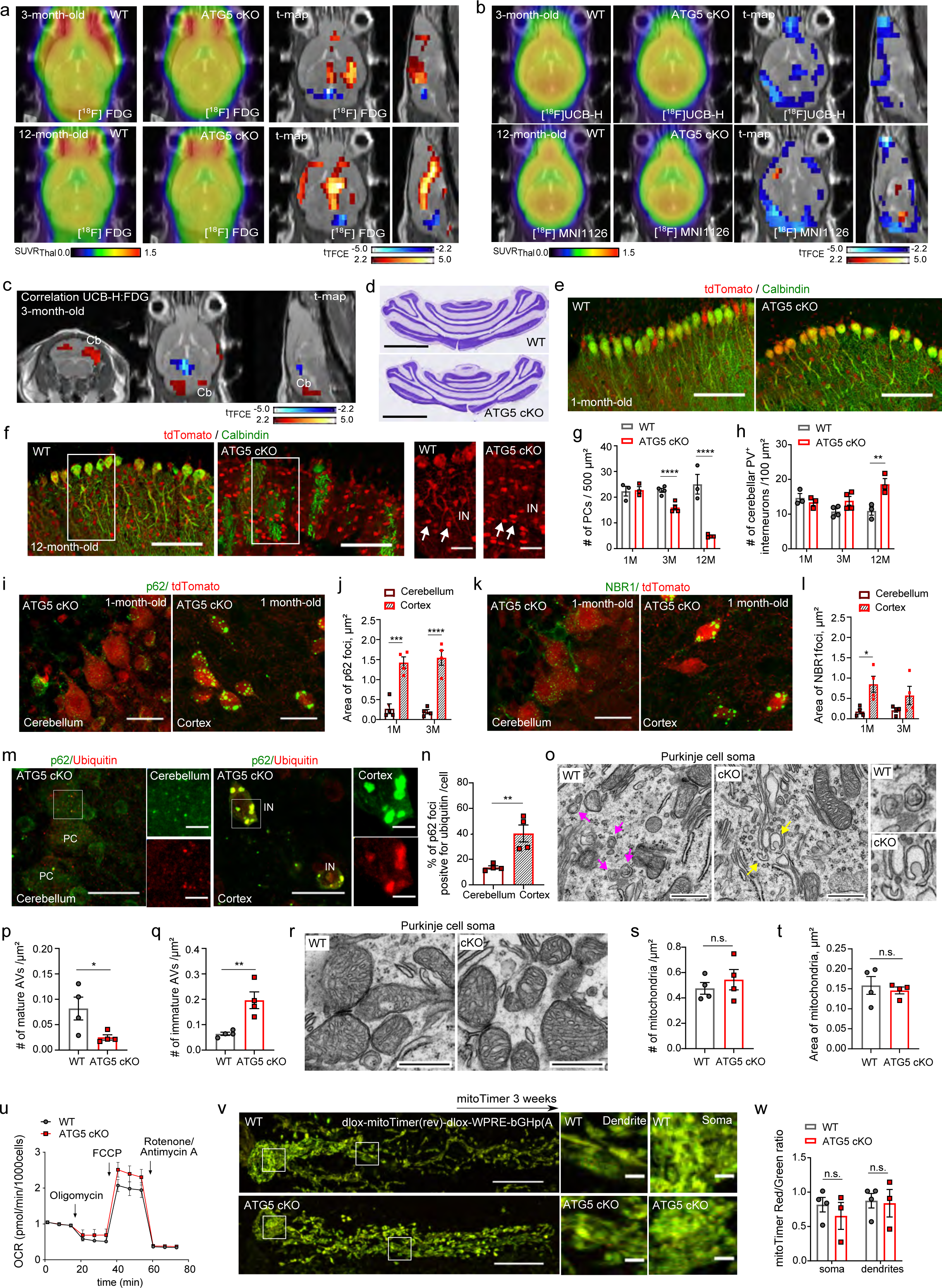
Loss of autophagy attenuates the vulnerability of cerebellar PCs independent of its role in protein and mitochondria quality control. **a,** Longitudinal [^18^F]FDG-PET imaging in 3 and 12 months old WT and ATG5 cKO mice (WT 3-month-old N=7, WT 12-month-old N=6, ATG5 cKO N=6 per age group). T-map with voxel-wise comparison between ATG5 cKO and WT (t-test, corrected for multiple testing) is shown to the right. Red and blue voxels indicate significantly (p<0.05) higher and lower tracer uptake in ATG5 cKO mice. **b**, Longitudinal [^18^F]UCB-H and [^18^F]MIN1126-PET imaging in in 3 and 12 month old WT and ATG5 cKO mice, respectively (WT 3-month-old N=7, WT 12-month-old N=6, ATG5 cKO N=6 per age group). T-map with voxel-wise comparison between ATG5 cKO and WT (t-test, corrected for multiple testing) is shown to the right. Red and blue voxels indicate significantly (p<0.05) higher and lower tracer uptake in ATG5 cKO mice. **c,** Correlation between [^18^F]UCB-H- and [^18^F]FDG-PET signals for 3-month old pooled ATG5 cKO and WT animals (Pearson correlation, t-test for significantly correlated voxels, corrected for multiple testing). Red and blue voxels indicate significantly (p<0.05) positive and negative correlation in ATG5 cKO mice. **d**, Representative Nissl-stained cerebellar sections from 3-month-old WT and ATG5 cKO mice. Scale bar, 2mm. **e,f,** Representative confocal images of the cerebellum from WT and ATG5 cKO mice at the age of 1 (e) and 12 (f) months expressing tdTomato as a reporter of Cre recombination, immunostained for calbindin. Arrows indicate cerebellar interneurons (IN). Scale bar: 100 µm (inserts 25µm). **g**, Analysis of calbindin-positive PC density in WT and ATG5 cKO cerebellum at the age of 1, 3 and 12 months. Each dot represents one animal (N=3 per genotype for 1 and 12 months old animals and N=4 per genotype for 3-month-old animals). Two-way ANOVA followed by Holm-Šidák multiple comparisons test (WT 1 month: 22.33 ± 1.88, ATG5 cKO 1 month: 22.78 ± 1.34, WT 3 months: 22.67 ± 0.79, ATG5 cKO 3 months: 15.99 ± 0.98; p = 0.015, WT 12 months: 25.04 ± 3.82, ATG5 cKO 12 months: 4.88 ± 0.29; p<0.0001). **h**, Analysis of parvalbumin (PV)-positive molecular cell layer interneuron density in WT and ATG5 cKO cerebellum at the age of 1, 3 and 12 months (see also Fig. S1j). Each dot represents one animal (N=3 per genotype for 1 and 12 months old animals, N=4 per genotype for 3 months-old animals). Two-way ANOVA followed by Holm-Šidák multiple comparisons test (WT 1 month: 14.68 ± 1.52, ATG5 cKO 1 month: 13.42 ± 0.89, WT 3 months: 10.78 ± 0.75, ATG5 cKO 3 months: 13.85 ± 1.25, WT 12 months: 10.92 ± 1.24, ATG5 cKO 12 months: 18.61 ± 1.63; p=0.0019). **i**, Representative confocal images of the cerebellum and cortex from ATG5 cKO mice at the age of 1 month expressing tdTomato as a reporter of Cre recombination, immunostained for p62 (see also Fig. S1k for WT control). Scale bar: 25 µm. **j**, Immunofluorescence analysis of the area of p62 foci in the somata of PCs and cortical interneurons of 1- and 3-month old ATG5 cKO animals (see also Fig. S1l). Each dot represents one animal (N=4 per genotype). Two-way ANOVA followed by Holm-Šidák multiple comparisons test (cerebellum 1 month: 0.27 ± 0.12, cortex 1 month: 1.42 ± 0.14; p<0.0001, cerebellum 3 months: 0.197 ± 0.05, cortex 3 months: 1.54 ± 0.18; p<0.0001). **k**, Representative confocal images of the cerebellum and cortex from ATG5 cKO mice at the age of 1 month expressing tdTomato as a reporter of Cre recombination, immunostained for NBR1 (see also Fig. S1m for WT control). Scale bar: 25 µm. **l**, Immunofluorescence analysis of the area of NBR1 foci in the somata of PCs and cortical interneurons of 1 and 3 month old WT and ATG5 cKO animals (see also Fig. S1n). Each dot represents one animal (N=4 per genotype). Two-way ANOVA followed by Holm-Šidák multiple comparisons test (cerebellum 1 month: 0.17 ± 0.05, cortex 1 month: 0.84 ± 0.19; p=0.019, cerebellum 3 months: 0.21 ± 0.05, cortex 3 months: 0.57 ± 0.22). **m**, Representative confocal images of the cerebellum and cortex from ATG5 cKO mice at the age of 3 months, immunostained for p62 and Ubiquitin. Scale bar: 25 µm (inserts 5µm). PC, Purkinje cell, IN-interneuron. **n**, Analysis of total p62 puncta per cell positive for ubiquitin in 3-month old cortex and cerebellum of ATG5 cKO mice. Each dot represents one animal (N=4 per genotype). Unpaired two-tailed t-Test between two groups (cerebellum: 13.62 ± 1.27, cortex: 40.3 ± 6.65; p=0.007). **o**, Representative EM images of WT and ATG5 cKO PC somata at the age of 3 months. Arrows in magenta indicate mature autophagosomes (AVs). Arrow in yellow indicate immature AVs. Scale bar 1 µm. **p**, Analysis of mature AV density in the soma of WT and ATG5 cKO mice. Each dot represents one animal (N=4 mice per genotype). Unpaired two-tailed t-Test between two groups (WT: 0.08 ± 0.02, ATG5 cKO: 0.02 ±0.005; p=0.048). **q,** Analysis of immature AV density in the PC soma of WT and ATG5 cKO mice. Each dot represents one animal (N=4 mice per genotype). Unpaired two-tailed t-Test between two groups (WT: 0.06 ± 0.006, ATG5 cKO: 0.19 ± 0.03; p=0.007). **r**, Representative EM images of mitochondria in the soma of WT and ATG5 cKO PCs. A. Scale bar 0.5 µm. **s,** Analysis of mitochondria density in the PC soma of WT and ATG5 cKO mice. N=4 mice per genotype. Each dot represents one animal (N=4 mice per genotype). Unpaired two-tailed t-Test between two groups (WT: 0.475 ± 0.045, ATG5 cKO: 0.544 ± 0.079). **t,** Analysis of mitochondria area in the soma of WT and ATG5 cKO mice. N=4 mice per genotype. Each dot represents one animal (N=4 mice per genotype). Unpaired two-tailed t-Test between two groups (WT: 0.158 ± 0.022, ATG5 cKO: 0.145 ± 0.0008). **u**, Oxygen consumption rate (OCR) during the Seahorse XF Cell Mito Stress test in WT and ATG5 cKO primary cerebellar cells (DIV17) (N=5 cultures per genotype). See Fig. S2b for analysis. **v**. Representative images of mitoTimer-expressing PCs in WT and ATG5 cKO cerebellum at the age of 3 months. Scale bar: 20µm (1 µm insert). **w**, mitoTimer-based fluorescence image analysis of mitochondria turnover (red fluorescence divided by green fluorescence) in PC somata and dendrites in WT and ATG5 cKO cerebellum at the age of 3 months. Each dot represents one animal with N=4 (WT)/3(KO) per genotype. Two-way ANOVA followed by Holm-Šidák multiple comparisons test (WT soma: 0.81 ± 0.1, ATG5 cKO soma: 0.65 ± 0.19, WT dendrites: 0.87 ± 0.1, ATG5 cKO dendrites: 0.84 ± 0.2). Data information: 1M, 3M and 12M indicate 1, 3 and 12 months, respectively. Squares in Fig. 1f, 1m, 1v indicate regions magnified. All graphs show mean ± SEM. n.s.—non-significant; * indicates P ≤ 0.05; ** indicates P ≤ 0.01; *** indicates P ≤ 0.001; **** indicates P ≤ 0.0001.

Cerebellar atrophy and loss of cerebellar PCs were reported in several mouse models with autophagy deficiency^50-53^. Thus, we analyzed the number of PCs by Nissl-staining and immunohistochemistry in WT and ATG5 cKO mice, additionally carrying the tdTomato allele (*Ai9*) as a reporter (i.e., *Atg5*wt/wt: *Slc32a1-*Cre^tg^: *Ai9* and *Atg5*flox/flox: *Slc32a1-*Cre^tg^: *Ai9* mice). We found that the cerebellum was smaller (Fig. 1d) and contained significantly less calbindin-positive PCs in 3-month-old ATG5 cKO mice compared to controls (Fig. 1e-g, Fig. S1i, see also Fig. 7j). The loss of PCs was not attributed to developmental defects, as their number was comparable in the cerebellum of 1-month-old WT and ATG5 cKO mice (Fig. 1e,g), and was progressive, with only a few PCs detected in the cerebellum lacking ATG5 at 12 months of age (Fig. 1f, g). Remarkably, this cell loss was selective for PCs, as inhibitory GABAergic interneurons remained present in the cerebellum of ATG5 cKO mice (Fig. 1f, Fig. S1j), and their number was even significantly increased in 12-month-old animals (Fig. 1h). These findings strongly suggest that the same type of neurons across various brain regions exhibit distinct levels of vulnerability to autophagy loss.

### Reverse correlation of ubiquitin-positive protein aggregates and neurodegeneration in autophagy-deficient neurons

To gain insight into the observed selective vulnerability of cerebellar PCs upon autophagy dysfunction, we first analyzed neuronal autophagy defects via p62 aggregate imaging using immunohistochemistry. We found that although p62 levels were significantly increased at 1 and 3 months of age in both ATG5 cKO cortex and cerebellum (see Fig. S1c,f), p62 was not aggregated in characteristic foci in autophagy-deficient cerebellar PCs (Fig. 1i,j, Fig. S1k,l). A similar phenotype was observed by analyzing the distribution of another autophagy receptor NBR1 (Fig. 1k,l, Fig. S1m,n). In contrast, p62 and NBR1 puncta formed aggregates in ATG5 cKO cortical neurons (Fig. 1i-l). At 3 months of age, approximately 40% of these p62-positive foci in cortical ATG5 cKO neurons were also positive for ubiquitin, whereas ubiquitin-positive p62 puncta were present in less than 15% of autophagy-deficient PCs (Fig. 1m,n).

Several studies have reported the existence of ATG5/ATG7-independent autophagy ^54, 55^. To evaluate if the absence of p62/NBR1 foci in ATG5 cKO PCs is due to the fact that they are still capable of forming autophagsomes, we analyzed autophagosome numbers in both WT and ATG5 cKO PCs by electron microscopy (EM). We found that while mature autophagosomes were abundant in WT PCs, their number was significantly reduced in PCs lacking ATG5 (Fig. 1o,p). Conversely, the number of unprocessed immature autophagosome intermediates was significantly increased in ATG5 cKO PC compared to control cells (Fig. 1q), indicating that the deletion of ATG5 significantly impairs autophagosome formation in cerebellar PCs.

Next, we asked if the impaired clearance of damaged mitochondria in autophagy-deficient PCs could account for their vulnerability. The mitochondria of ATG5 cKO PCs were similar in morphology to WT mitochondria analyzed by EM (Fig. 1r-t, Fig. S2a). Moreover, Seahorse-based mitochondrial respiration was unaltered in cultured cerebellar ATG5 cKO neurons (Fig. 1u, Fig. S2b), and no change in mitochondria turnover was detected in ATG5 cKO PCs compared to controls, as measured by mitoTimer ratio analysis in adeno-associated virus (AAV)-transduced PCs *in vivo* (Fig. 1v,w). The absence of mitophagy defects was also evident in the unaltered ratio of the mitochondria-localized-mKeima-Red (mito-mKeima) reporter in PCs *ex vivo* (Fig. S2c,d), along with unchanged levels of the mitophagy receptor BNIP3 (Fig. S2e,f). These data suggest that mitophagy in cerebellar PCs may proceed independently of ATG5. Moreover, these results indicate that the increased susceptibility of PCs to neurodegeneration under autophagy deficiency cannot be attributed to the accumulation of protein aggregates and/or damaged mitochondria.

### Multimodal omics analysis reveals accelerated glycolysis in ATG5 cKO cerebellum

To reveal the precise cellular mechanisms behind neurodegeneration in the ATG5 cKO cerebellum, we conducted a comprehensive analysis of the cerebellar proteome in 1-month-old (before neurodegeneration) and 3-month -old (after the onset of neurodegeneration) WT and ATG5 cKO animals. A total of 609 and 853 significantly dysregulated proteins (with a log2 fold change ><0.25) were identified in the cerebellum of ATG5 cKO mice at the age of 1 and 3 months, respectively (Fig. 2a,b, Table S1). KEGG pathway analysis of the upregulated proteins between both age groups indicated a predominant association with metabolic pathways, carbon metabolism, and pathways in cancer. In contrast, the downregulated proteins were clustered in pathways associated with neurodegenerative diseases (Fig. 2c,d). Gene Ontology (GO) biological process and GO cellular compartment analyses revealed alterations in protein transport and localization to membrane compartments, synapses and mitochondria (Fig. S3a-d). Only 32 proteins were commonly upregulated in the ATG5 cKO cerebellum at 1 and 3 months, and were clustered in pathways of metabolism, autophagy and ferroptosis (Fig. 2e, Fig. S3e). Commonly downregulated proteins were not associated with any KEGG pathway and were mostly components of cellular protein localization and synapses (Fig. S3f-h).

**Figure 2.**
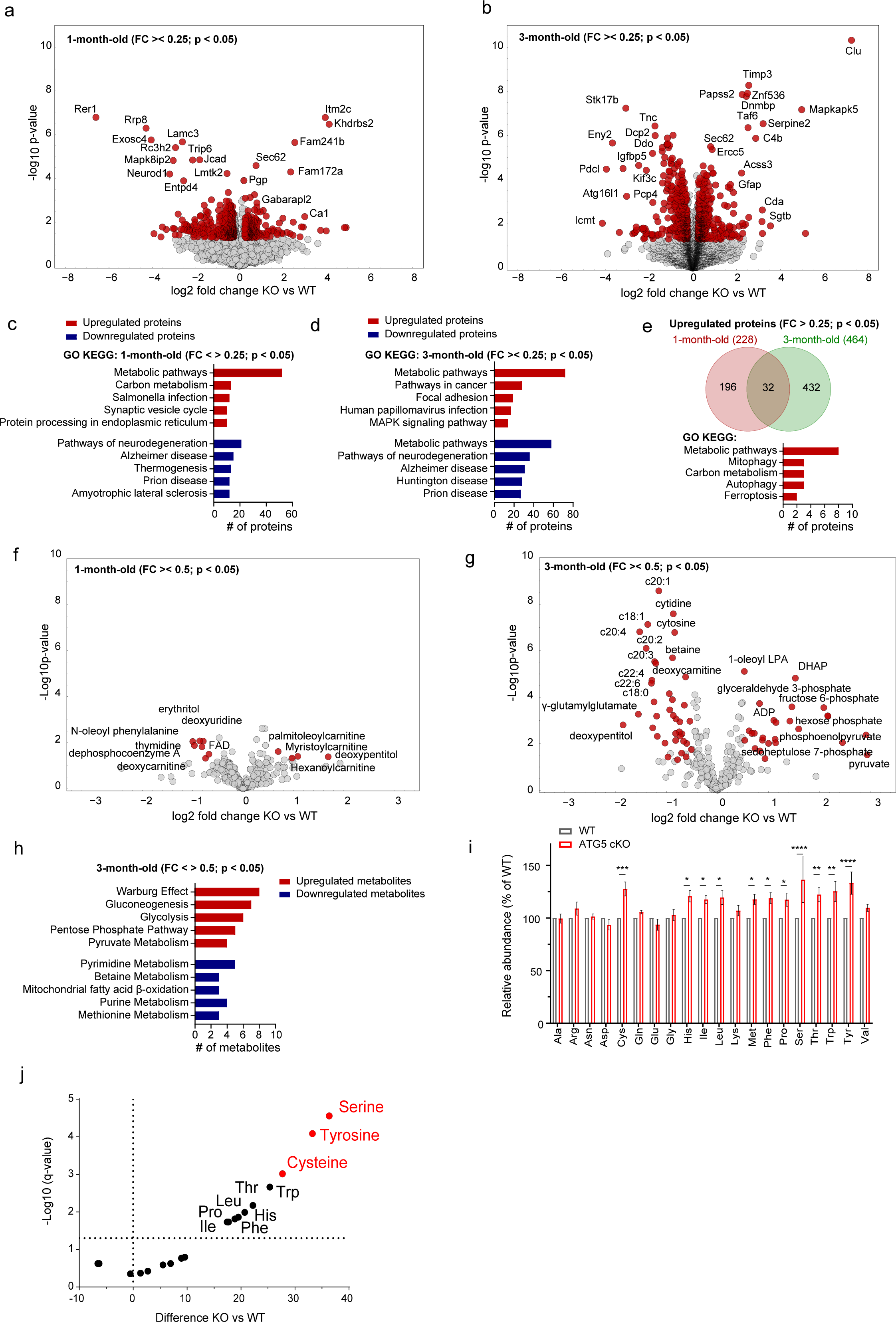
Proteomic and metabolic analyses identify metabolic rewriting in the ATG5 cKO cerebellum. **a,b,** Volcano plot of differentially expressed proteins in WT and ATG5 cKO cerebellum at the age of 1 month (a) and 3 months (b), analyzed using label-free proteomic approach. Red colored circles mark significantly deregulated proteins at p < 0.05 and log2 fold change of < > 0.25. See also Table S1. N=5 per genotype. **c**,**d**, ShinyGO v0.741-based GO analysis of KEGG-enriched terms in the cerebellar proteome (p < 0.05 and log2 fold change of < > 0.25) at 1 month (c) and 3 months of age (d). See also Fig. S3a-d for GO BP and GO CC analyses. **e**, Venn diagram of commonly upregulated proteins (p < 0.05 and log2 fold change of < > 0.25) in the 1- and 3 months-old ATG5 cKO cerebellum. See also Fig. S3f-h for commonly downregulated proteins. **f**,**g**, Volcano plot of differentially abundant metabolites in WT and ATG5 cKO cerebellum at the age of 1 month (f) and 3 months (g), analyzed using untargeted metabolomics approach. Red colored circles mark significantly deregulated metabolites at p < 0.05 and log2 fold change of < > 0.5. See also Table S2. N=5 per genotype. **h,** MetaboAnalyst-based pathway analysis of significantly upregulated and downregulated metabolites in 3-month-old ATG5 cKO cerebellum (p < 0.05 and log2 fold change of < > 0.5). **i**, Relative amino acid profile in 1-month old WT and ATG5 cKO cerebellum identified using targeted metabolomics approach. Multiple unpaired T-tests with linear Benjamini, Krieger and Yekutieli correction (WT levels set to 100%, ATG5 cKO serine: 136.35 %*±* 21.66; q=0.00003, ATG5 cKO tyrosine: 133.22 % ± 10.68; q=00001, ATG5 cKO cysteine: 127.67 % ± 6.65; q=0.00096, ATG5 cKO tryptophan: 125.31 % ± 9.53; p=0.0021, ATG5 cKO threonine: 122.19 % ± 6.69; q=0.0067, ATG5 cKO histidine: 120.67 % ± 5.34; q=0.0102, ATG5 cKO leucine: 119.5 % ± 6.83; q=0.013, ATG5 cKO phenylalanine: 118.83 % ± 5.21; q=0.015, ATG5 cKO methionine: 117.68 % ± 4.76; q=0.018, ATG5 cKO isoleucine: 117.6 % ± 3.72; q=0.018, ATG5 cKO proline:117.4 % ± 21.66; q=0.018, see also supplement Fig. S2). **j**, Volcano plot of the differentially abundant amino acids in 1-month-old ATG5 cKO cerebellum shown in (i). Red colored circles mark strongly upregulated amino acids. Data information: All graphs show mean ± SEM. n.s.—non-significant; * indicates P ≤ 0.05; ** indicates P ≤ 0.01; *** indicates P ≤ 0.001; **** indicates P ≤ 0.0001.

The proteome dataset described above suggests that the conditional loss of ATG5 in the cerebellum results in the dysregulation of metabolic pathways beginning as early as one month of age. To gain a deeper understanding of the metabolome changes that might occur in the autophagy-deficient cerebellum, we performed a semi-targeted metabolomic analysis. At one month of age, only a few metabolite changes were detected in the ATG5 cKO cerebellum compared to the WT (Fig. 2f), whereas the metabolome of the ATG5 cKO cerebellum was prominently altered at 3 months of age (Fig. 2g, Table S2). Upregulated metabolites clustered in metabolic pathways associated with glucose metabolism, the pentose phosphate pathway, and pyruvate metabolism (Fig. 2h). Downregulated metabolites were associated with pyrimidine metabolism and mitochondrial fatty acid β-oxidation.

The supply of amino acids (AA) is an important function of autophagy to promote cell survival ^56^. Surprisingly, we did not detect a decrease in AA content in the cerebellum of mice conditionally lacking ATG5 (Fig. 2i), suggesting that cerebellar AA production via autophagy likely functions primarily under nutrient-deprived conditions. Conversely, we observed a significant increase in AAs in ATG5 cKO cerebellar lysates, and, interestingly, some of these highly upregulated AAs can be synthesized by the intermediates of the glycolysis pathway, for instance, serine and cysteine^57^ (Fig. 2i,j). This implies that the loss of cerebellar autophagy leads to a rewiring of glycolytic metabolism.

### Conditional ATG5 loss in the cerebellum increases glucose flux and promotes glycolytic capacity

To further characterize the glycolytic changes in the cerebellum of ATG5 cKO mice, we next performed a targeted LC-MS/MS metabolomic analysis of glycolytic intermediates. Already at 1 month of age, there was a significant increase in glycolytic intermediates measured in the ATG5 cKO cerebellum, such as fructose-6-phosphate, glucose-6-phosphate, mannose-6-phosphate and sedoheptulose-7-phosphate (Fig. 3a, Table S3). The increase in fructose-6-phosphate and glucose-6-phosphate became even more pronounced in the cerebellum of the 3-month-old animals (Fig. 3b). Additionally, the levels of 6-phosphogluconate and fructose-1,6-biphosphate were also significantly upregulated. In agreement with unchanged mitochondrial respiration measured by the Seahorse Metabolic Analyzer in cultured cerebellar ATG5 cKO neurons (see Fig. 1u), we detected no alterations in the tricarboxylic acid (TCA) cycle in the 1-month-old ATG5 cKO cerebellum (Fig. S4a). Interestingly, we observed a small, but significant decrease in TCA intermediates in the 3-month-old ATG5 cKO cerebellum (Fig. S4b), suggesting that excessive glycolysis (already occurring as early as 1 month of age) may lead to a later inhibition of mitochondria respiration, reminiscent of a phenomenon previously described in proliferating non-neuronal cells ^58^ and brain astrocytes ^59^.

**Figure 3.**
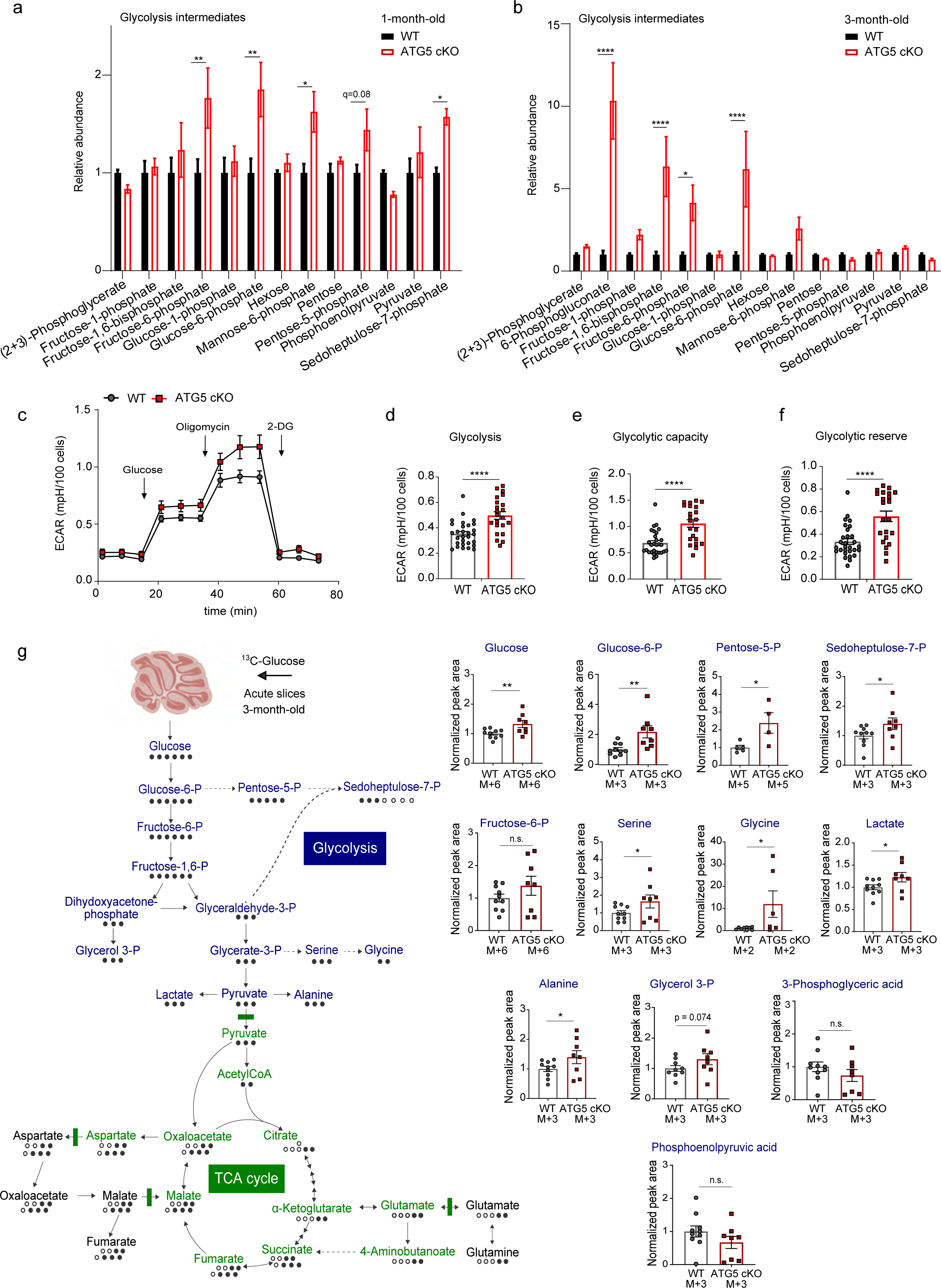
The conditional loss of ATG5 results in upregulated glycolysis and increases the glycose flux in the cerebellum. **a,b,** Relative profile of glycolytic intermediates in 1-month old (a) and 3-month-old (b) WT and ATG5 cKO cerebellum identified using targeted metabolomics approach (N=5 per genotype). Multiple unpaired T-tests with linear Benjamini, Krieger and Yekutieli correction (WT vs ATG5 cKO at 1 month: Glucose-6-phosphate q=0.0013, Fructose-6-phosphate q=0.0028, Mannose-6-phosphate q=0.0147, Sedoheptulose-7-phosphate q=0.0217; WT vs ATG5 cKO at 3 month: Glucose-6-phosphate q=0.00001, Fructose-6-phosphate q=0.0106, Fructose-1,6-biphosphate q=0.00001, 6-Phosphogluconate q<0.000001 (further statistics description is in source data for Fig.3). **c,** Extracellular acidification rates (ECAR), produced by the flux of H^+^ upon conversion of glucose to lactate in primary cerebellar cells (DIV17) from WT (N=5) and ATG5 cKO (N=6) mice during Seahorse XF glycolysis stress test with injections of 10 mM glucose, 1.5 µM oligomycin and 50 nM 2-DG. **d**, Analysis of glycolysis rates in the Seahorse XF glycolysis stress test shown in (c). Each dot represents one well with ± 20 000 cells (from N=5 WT and N=6 ATG5 cKO mice). Two tailed unpaired T-test (WT: 0.35 ± 0.018, ATG5 cKO: 0.49 ± 0.02; p<0.0001). **e**, Analysis of glycolytic capacity in the Seahorse XF glycolysis stress test shown in (c). Each dot represents one well with ± 20 000 cells (from N=5 WT and N=6 ATG5 cKO mice). Two tailed unpaired T-test (WT: 0.68 ± 0.04, ATG5 cKO: 1.05 ± 0.07; p<0.0001). **f**, Analysis of glycolytic reserve in the Seahorse XF glycolysis stress test shown in (c). Each dot represents one well with ± 20 000 cells (from N=5 WT and N=6 ATG5 cKO mice). Two tailed unpaired T-test (WT: 0.33 ± 0.02, ATG5 cKO: 0.55 ± 0.04; p<0.0001). **g**, Schematic and analysis of metabolic flux of glucose in acute cerebellar 3 month-old WT and ATG5 cKO slices using ^13^C-glucose as a tracer (from N=10 WT and N=8 ATG5 cKO mice). One tailed unpaired T-test (WT levels set to 1, ATG5 cKO glucose: 1.32 ± 0.12; p=0.007, ATG5 cKO glucose-6-P: 2.16 ± 0.4; p=0.0038, ATG5 cKO pentose-5-P: 2.38 ± 0.57; p=0.017, ATG5 cKO seduheptulose-7-p: 1.4 ± 0.19; p=0.03, ATG5 cKO fructose-6-p: 1.37 ± 0.29, ATG5 cKO serine: 1.64 ± 0.36; p=0.044; ATG5 cKO glycine: 11.96 ± 5.95; p=0.019, ATG5 cKO lactate: 1.22 ± 0.1; p=0.03, ATG5 cKO alanine: 1.39 ± 0.22; p=0.045, ATG5 cKO glycerol-3-P: 1.3 ± 0.18; p=0.074; ATG5 cKO 3-pohosphoglyceric acid: 0.73 ± 0.18, ATG5 cKO phoshoenolpyruvate: 0.66 ± 0.18). Data information: All graphs show mean ± SEM. n.s.—non-significant; * indicates P ≤ 0.05; ** indicates P ≤ 0.01; *** indicates P ≤ 0.001; **** indicates P ≤ 0.0001.

Next, we directly assessed the glycolytic capacity of WT and ATG5-deficient cerebellar neurons cultured in 10 mM glucose. Using the Seahorse Metabolic Analyzer to measure extracellular acidification rate (ECAR), we observed a marked increase in glycolytic function in cKO neurons compared to WT (Fig. 3c-f). Glycolytic capacity was further upregulated in ATG5 KO neurons cultured in 30 mM glucose (Fig. S4c,d), suggesting that higher glucose availability facilitates glycolytic metabolism in autophagy-deficient neurons. Finally, we also evaluated the impact of ATG5 deficiency on glucose flux in acute cerebellar WT and ATG5 cKO slices by the ^13^C-labeling technique using ^13^C-glucose as a tracer. We found a significant increase in total ^13^C incorporation in the majority of glycolytic intermediates (including glucose, lactate, pentose-5-phosphate, and sedoheptulose-7-phosphate) as well as in serine, glycine and alanine in cKO slices (Fig. 3g, Fig. S4e, Table S4). This indicates an overall increase in glycolytic flux in the autophagy-deficient cerebellum. Notably, glucose flux into the TCA cycle remained unaltered between cerebellar slices from WT and ATG5 cKO animals (Fig. S4f).

The data described above show that ATG5 loss increases glucose metabolism in the cerebellum. To further test whether autophagy-deficient PCs take up more glucose compared to controls, we utilized the fluorescent glucose substrate 2-(N-(7-nitrobenz-2-oxa-1,3-diazol-4-yl)amino)-2-deoxyglucose (2-NBDG) and measured its uptake in PCs ex-vivo using organotypic WT slice culture (Fig. S5a). The results demonstrated a significant facilitation of glucose uptake in ATG5-deficient PCs (Fig. 4a,b). Concomitant with increased glucose uptake, autophagy-deficient PCs also produced more lactate, measured by AAV-mediated expression of the lactate FRET sensor “laconic” ^60^ in organotypic cerebellar slices treated with 1 µM of the monocarboxylate transporter (MCT)-1 and MCT-2 inhibitor AR-C155858 to prevent extracellular lactate export (Fig. 4c-e). Notably, this phenotype was selective to PCs, as ATG5-deficient molecular layer interneurons did not differ from controls in their lactate production (Fig. 4f, Fig. S5b). To ascertain whether the elevated lactate levels in ATG5 cKO PCs are also associated with excessive ATP production through aerobic glycolysis, we measured cytoplasmic ATP levels in WT and ATG5 cKO PCs using the genetically encoded ATeam1.03^YEMK^ FRET sensor under control conditions and conditions where mitochondrial ATP production was inhibited. The levels of cytoplasmic ATP were significantly higher in ATG5 cKO PCs compared to controls (Fig. 4g,h) and remained elevated even after inhibition of mitochondrial respiration with oligomycin A (Fig. 4i-k). Of note, the remaining ATP levels in WT PCs after blocking mitochondrial respiration (Fig. 4j,k) suggest a significant contribution of glycolysis to the energy demand of PCs, at least under *ex vivo* conditions. Consistent with data indicating increased ATP production in autophagy-deficient PCs, we did not detect a decrease in phosphorylated AMP-activated protein kinase (AMPK) levels in cerebellar ATG5 cKO lysates (Fig. S5c,d).

**Figure 4.**
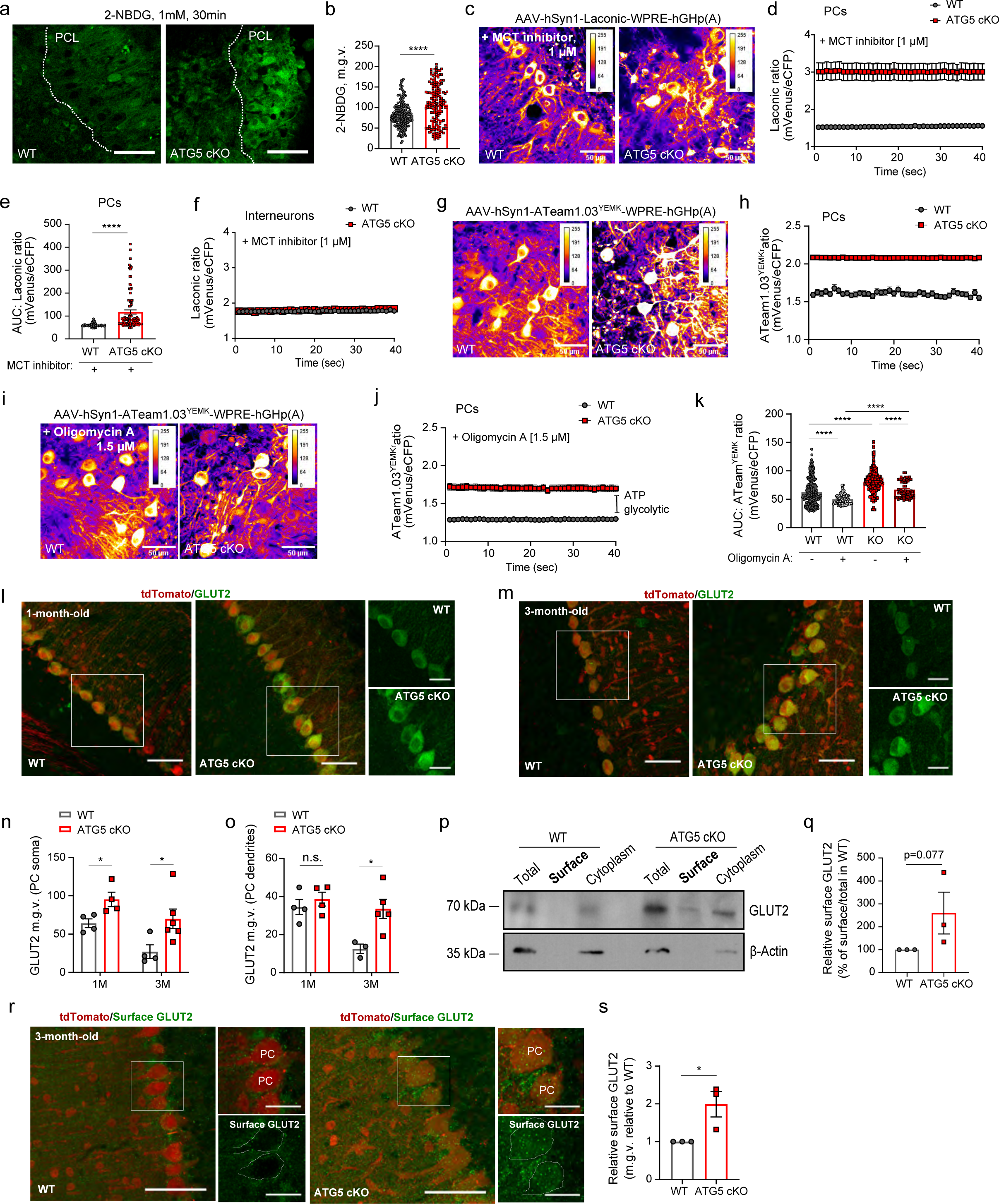
Excessive glycolysis in ATG5 cKO PCs correlates with their elevated GLUT2 levels. **a,b** Representative confocal images of 2-NBDG uptake (a) and analysis of 2-NBDG levels (b) in WT and ATG5 cKO cerebellar organotypic slice culture at DIV 21. Scale bar: 50 µm. PCL-PC layer. Each dot represents a single PC (for WT 190 cells from N=7 and for ATG5 cKO 168 cells from N=5 mice). Two-tailed unpaired T-test (WT: 82.37 ± 2.19, ATG5 cKO: 107.4 ± 3.79; p<0.0001). **c**, FRET-based imaging of WT and ATG5 cKO cerebellar organotypic slice cultures transduced with AAV-hSyn1-Laconic. Scale bar: 50 µm. Slices were treated with 1 µM of MCT-1/2 inhibitor AR-C155858 to prevent extracellular lactate export. **d**, Ratiometric analysis of Laconic signal (mVenus/eCFP ratio) in WT (83 cells from N=6) and ATG5 cKO (93 cells from N=5) PCs. **e**, Area under the curve (AUC) analysis of Laconic signal (mVenus/eCFP ratio) in WT and ATG5 cKO PCs. Each dot represents a single PC (83 cells from N=6 for WT and 93 cells from N=5 for ATG5 cKO). Two-tailed unpaired T-test (WT: 60.13 ± 0.84, ATG5 cKO: 117.3 ± 8.92; p<0.0001). **f**, Ratiometric analysis of Laconic signal (mVenus/eCFP ratio) in WT (158 cell from N=6) and ATG5 cKO (134 cells from N=5) cerebellar interneurons. **g**, FRET-based imaging of WT and ATG5 cKO cerebellar organotypic slice cultures transduced with AAV-hSyn1-ATeam1.03YEMK. Scale bar: 50 µm. **h**, Ratiometric analysis of ATeam1.03YEMK signal (mVenus/eCFP ratio) in WT (488 cells from N=6) and ATG5 cKO (317 cells from N=6) PCs. **i**, FRET-based imaging of WT and ATG5 cKO cerebellar organotypic slice cultures transduced with AAV-hSyn1-ATeam1.03YEMK and treated with 1,5µM of Oligomycin A to inhibit the OXPHOS function. Scale bar: 50 µm. **j**, Ratiometric analysis of ATeam1.03YEMK signal (mVenus/eCFP ratio) in WT (131 cells from N=3) and ATG5 cKO (66 cells from N=4) PCs treated with 1,5µM of Oligomycin A. **k**, Area under the curve (AUC) analysis of ATeam1.03YEMK signal in WT and ATG5 cKO PCs. Each dot represents a single PC (N=6 per genotype with 488 cells for WT and 317 cells for ATG5 cKO PCs without oligomycin A and 131 cells from N=3 WT and 66 cells from N=4 ATG5 cKO oligomycin A-treated PCs). Two-way ANOVA followed by Holm-Šidák multiple comparisons test (WT without oligomycin: 62.57 ± 1.02, WT with oligomycin: 50.2 ± 0.75; p<0.0001, ATG5 cKO without oligomycin: 81.16 ± 1.1; p<0.0001, ATG5 cKO with oligomycin: 66.88 ± 1.75; p<0.0001, WT with oligomycin vs ATG5cKO with oligomycin p<0.0001). **l,m**, Representative confocal images of WT and ATG5 cKO cerebellum at the age of 1 (l) and 3 (m) months expressing tdTomato as a reporter of Cre recombination, immunostained for GLUT2. Scale bar: 50 µm, inserts 10 µm. **n,** Immunofluorescence analysis of GLUT2 protein levels in PC soma of 1 and 3 month old WT and ATG5 cKO mice. Each dot represents one animal (N=4 per genotype for 1 month old animals and 3-month-old WT animals, N=6 for 3 months old for ATG5 cKO animals). Two-way ANOVA mix model followed by Holm-Šidák multiple comparisons test (WT 1 month: 64.24 ± 5.7, ATG5 cKO 1 month: 95.26 ± 9.43; p=0.031, WT 3 months: 27.05 ± 8.92, ATG5 cKO 3 months: 69.97 ± 12.75; p=0.022). **o,** Immunofluorescence analysis of GLUT2 protein levels in PC dendrites of 1 and 3 month old WT and ATG5 cKO mice. Each dot represents one animal (N=4 per genotype for 1-month old animals, N=3 for 3-month-old WT and N=5 for 3-month-old ATG5 cKO animals, N=3 for 3-month old WT animals). Two-way ANOVA followed by Holm-Šidák multiple comparisons test (WT 1 month: 34.47 ± 3.96, ATG5 cKO 1 month: 38.6 ± 3.61, WT 3 months: 12.54 ± 2.47, ATG5 cKO 3 months: 33.57 ± 5.02; p=0.02). **p**,**q,** Immunoblot-based biotinylation analysis of surface GLUT2 levels in WT and ATG5 cKO cerebellar acute slices at the age of 3 months. Each dot represents one animal (N=3 per genotype). Two-tailed unpaired T-test (WT level were set to 100 %, ATG5 cKO: 259.7 ± 91.3, p=0.077). **r**, Representative confocal images of WT and ATG5 cKO cerebellum at the age of 3 months expressing tdTomato as a reporter of Cre recombination, immunostained for surface GLUT2. Scale bar: 50 µm, inserts 10µm. PC somata are outlines in images magnified to the right. **s**, Immunofluorescence analysis of GLUT2 surface levels in PC soma in WT and ATG5 cKO cerebellum at the age of 3 months. Each dot represents one animal (N=3 per genotype). Two-tailed unpaired T-test (WT level were set to 1, ATG5 cKO: 1.98 ± 0.33, p=0.021). Data information: 1M and 3M indicate 1 and 3 months, respectively. PC, Purkinje cell. m.g.v.-mean gray value. Squares in Fig. 4l, 4m, 4r indicate regions magnified. All graphs show mean ± SEM. n.s.—non-significant; * indicates P ≤ 0.05; ** indicates P ≤ 0.01; *** indicates P ≤ 0.001; **** indicates P ≤ 0.0001.

### Loss of ATG5 increases glucose transporter 2 (GLUT2) levels in PCs

We next explored the connection between the loss of autophagy and increased glucose cerebellar metabolism. Enhanced glucose uptake and glycolysis have been reported in cells overexpressing different members of the *Slc2* family of facilitative glucose transporters (GLUT) ^61-64^. Out of the 14 GLUT family members present in mammals, the transporters GLUT1, GLUT2, GLUT3, and GLUT4 are expressed in the brain^65^. Thus, we next analyzed the protein levels of GLUT1-4 in PCs in 1-month and 3-month-old WT and ATG5 cKO mice by immunohistochemistry. Levels of GLUT1, 3 and 4 were unaltered in ATG5 cKO PCs (Fig. S5e-j). In contrast, we found that GLUT2 levels were significantly increased in PCs lacking ATG5, and that this increase was evident as early as 1 month of age (Fig. 4l-n). The accumulation of GLUT2 was initially observed at the cell soma, and was subsequently evident in the dendrites of 3-month-old ATG5 cKO PCs (Fig. 4o). Consistent with increased glycolysis in the 1-month-old cerebellum and elevated GLUT2 levels in PCs at this age, we also found that the levels of hexokinase 2 (HK2), which is a major enzyme in glucose phosphorylation to produce glucose-6-phopshate, were also significantly upregulated in 1-month-old PCs in ATG5 cKO mice (Fig. S5k,l). The increase in GLUT2 proteins levels was not due to a change in mRNA levels measured by qPCR (Fig. S5m). Since glucose uptake typically involves the translocation of GLUT2 from an intracellular pool to the plasma membrane ^66^, we hypothesized that a portion of the upregulated GLUT2 is localized at the plasma membrane in ATG5 cKO cerebellum. Indeed, we detected a substantial, yet insignificant (p=0.077), increase in total surface levels in the cerebellum conditionally lacking ATG5 (Fig. 4p,q). Given that the whole cerebellar lysate contains various cell types, including glial and granule cells, which could attenuate the effect of ATG5 cKO on GLUT2 surface levels, we conducted a specific analysis of GLUT2 surface levels in WT and ATG5 cKO PCs using an N-terminus-specific GLUT2 antibody (binding to the GLUT2 exoplasmic loop between helices 1 and 2) (Fig. 4r). This analysis revealed a significant upregulation of surface GLUT2 levels in PCs lacking ATG5 (Fig. 4s). In summary, these findings suggest that the increased GLUT2 protein levels may contribute to the excessive glycolysis observed in PCs lacking ATG5.

### GLUT2 undergoes degradation by the autophagy-endolysosomal system in PCs *in vivo*

Several components of the glycolytic pathway have been previously reported to be degraded by autophagy in non-neuronal cells^28-31^. We therefore hypothesized that autophagy, via its crucial modifier ATG5, limits glycolysis by degrading cerebellar GLUT2, a function that could be critical in the transition from the adolescent to the mature brain, when the need for glycolysis decreases significantly ^67^. To test this hypothesis, we first analyzed the levels of GLUT2 in the WT cerebellum during the course of maturation. We found that GLUT2 levels in PCs drop significantly between 1 month and 3 months of age (Fig. 5a-c, Fig. S6a). This decrease in GLUT2 was associated with a concomitant increase in the autophagosome number in 3-month-old PCs compared to 1 month, visualized by the *bone fide* marker of autophagosomes LC3 (Fig. 5d,e). The decrease in GLUT2 during maturation was likely due to its degradation via the autophagy-endolysosomal system, since GLUT2 levels were significantly upregulated in the acute cerebellar slices treated with the lysosomotropic agent chloroquine (Fig. 5f,g) and significantly decreased when cerebellar slices were subjected to starvation using artificial cerebrospinal fluid (ACSF), a buffered salt solution with 10 mM glucose (Fig. 5h,i). This effect was cerebellum-specific, as GLUT2 levels remained unaltered in chloroquine-treated acute cortical slices albeit significant inhibition of autophagy under both conditions (Fig. 5g, Fig. S6b,c). Of note, the ratio of LC3II/LC3I, used as a readout of autophagy flux upon chloroquine treatment, was slightly higher in the cerebellar lysates, which may suggest a higher level of constitutive autophagy in the cerebellum compared to the cortex (Fig. S6c). In line with this, the number of LC3 puncta in the WT cortex was low and remained unaltered between the ages of 1 month and 3 months (Fig. S6d,e).

**Figure 5.**
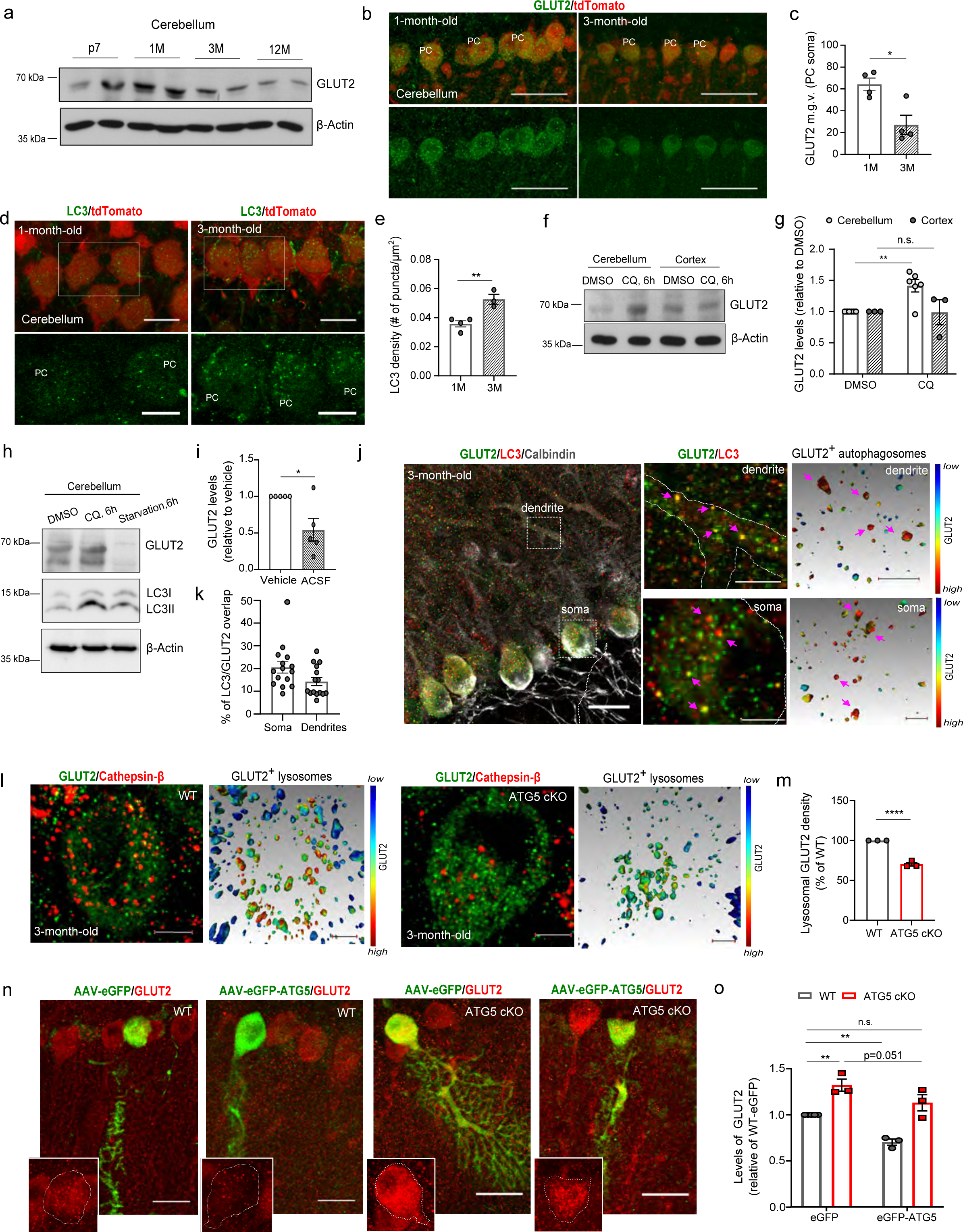
Cerebellar GLUT2 is degraded by ATG5-dependent autophagy. **a,** Representative immunoblot of GLUT2 levels in the WT cerebellum in the course of its maturation. **b**,**c,** Representative confocal images and fluorescence-based analysis of GLUT2 protein levels in PCs of 1 and 3 months old WT cerebellum immunostained for GLUT2. Each dot represents one animal (N=4 for each condition). Two-tailed unpaired T-test (1 month: 64.24 ± 5.7, 3 months: 27.05 ± 7.18; p=0.012). Scale bar 50µm. **d,e**, Representative confocal images and fluorescence-based analysis of LC3 puncta density in PCs of 1 and 3 month old WT cerebellum immunostained for LC3. WT and ATG5 cKO mice were expressing tdTomato as a reporter of Cre recombination. Scale bar, 20 µm, insert 10µm. Each dot represents one animal (N=4 for 1-month old and N=3 for 3-month old animals). Two-tailed unpaired T-test (1 month: 0.035 ± 0.002, 3 months: 0.05 ± 0.003; p=0.0072). **f,g**, Immunoblot analysis of GLUT2 protein levels in control and chloroquine (CQ)-treated (6h) cerebellar and cortical acute slices of 3-month old WT animals. Each dot represents one animal (N=6 for cerebellum and N=3 for cortex). Two-way ANOVA followed by Holm-Šidák multiple comparisons test (Levels in DMSO-treated slices were set to 1, cerebellum + CQ: 1.45 ± 0.098; p=0.0048, cortex + CQ: 0.99 ± 0.2). **h,i,** Immunoblot analysis of GLUT2 protein levels in control, chloroquine (CQ)-treated and in subjected to starvation in ACSF (6h) cerebellar acute slices of 3-month old WT animals. Each dot represents one animal (N=5 per condition). One-tailed unpaired T-test (Levels in Vehicle-treated slices were set to 1, ACSF: 0.540 ± 0.160; p=0.0105). **j,** Representative confocal images of WT cerebellum at the age of 3 months immunostained for GLUT2, LC3 and calbindin. Calbindin was used as PC marker. AMIRA-based 3D reconstructions of magnified regions are shown to the right. Colocalization of LC3 and GLUT2 is color-coded, with the warm colors indicating strong colocalization between two channels (see color-coded horizontal bar for the definition). Arrows indicate GLUT2^+^ autophagosomes. Scale bar: 20 µm, magnified regions, 5µm, 3D reconstruction 1µm. **k**, Colocalization analysis of LC3 overlap with GLUT2 in PC soma in WT cerebellum at the age of 3 months. Each dot represents one section for N=3 mice. (PC soma: 20.51% ± 2.53 %, PC dendrites: 14.26 % ± 1.73 %). **l,** Representative confocal images and AMIRA-based 3D reconstruction of GLUT2^+^ lysosomes in WT and ATG5 cKO cerebellum at the age of 3 months. Cathepsin-β was used as lysosomal marker. AMIRA-based 3D reconstructions of GLUT2^+^ lysosomes is color-coded, with the warm colors indicating strong colocalization between two channels (see color-coded horizontal bar for the definition). Scale bar 5 µm in confocal images, 1 µm in 3D-reconstrctions. **m**, AMIRA-based colocalization analysis of GLUT2 with β-Cathepsin D in WT and ATG5 cKO PCs at the age of 3 months. Levels in cKO were normalized to WT and set as 100%. Each dot represents one animal with N=3. One-tailed unpaired T-test (ATG5 cKO: 70.36 ± 2.23 p<0.0001). **n**, Representative confocal images of PCs in 3 month-old WT and ATG5 cKO cerebellum transduced either with EGFP-ATG5 or EGFP as a control. Scale bar: 20 µm. **o**, Immunofluorescence analysis of GLUT2 levels in EGFP- or EGFP-ATG5-overexpressing WT and ATG5 cKO PCs. Each dot represents one animal (N=3 per genotype for ATG5 cKO animals and WT +eGFP-ATG5, N=5 for WT animals + eGFP). GLUT2 levels were normalized to WT-eGFP and set as 1. Two-way ANOVA followed by Holm-Šidák multiple comparisons test (ATG5 cKO eGFP: 1.32 ± 0.06; p=0.0024 compared to WT-eGFP, p=0.051 compared to ATG5 cKO eGFP-ATG5, WT eGFP-ATG5: 0.7 ± 0.034, p=0.0032 compared to WT eGFP, ATG5 cKO eGFP-ATG5: 1.13 ± 0.08). Data information: p7, 1M, 3M and 12M indicate postnatal day 7, 1, 3 and 12 months, respectively. m.g.v.-mean gray value. PC, Purkinje cell. Squares in Fig. 5d and 5j indicate regions magnified. All graphs show mean ± SEM. n.s.—non-significant; * indicates P ≤ 0.05; ** indicates P ≤ 0.01; *** indicates P ≤ 0.001; **** indicates P ≤ 0.0001.

To better understand whether GLUT2 is trafficked within autophagosomes in PCs, we analyzed its colocalization with LC3 in the PC soma and dendrites using AMIRA-based 3D reconstruction in 3-month-old WT mice (Fig. 5j). We found that approximately 20% of LC3-positive autophagosomes contained GLUT2 (Fig. 5k). Moreover, in agreement with the previously reported lysosomal degradation of GLUT2 in non-neuronal cells ^68^, GLUT2 was abundant in WT lysosomes identified by cathepsin-β expression, whereas its lysosomal density was markedly reduced in ATG5 cKO PCs (Fig. 5l,m). Overexpression of ATG5 was previously reported to activate autophagy *in vivo* ^69^. Thus, we hypothesized that if GLUT2 is degraded by autophagy, overexpression of ATG5 should restore its levels in autophagy-deficient PCs. Consistent with this hypothesis, we found that GLUT2 protein levels were significantly downregulated in WT PCs overexpressing eGFP-ATG5, and that GLUT2 could be restored to near-physiological WT levels by re-expressing ATG5 in cKO PCs (Fig. 5n,o). Taken together, our data demonstrate that GLUT2 undergoes degradation by the autophagy-endolysosomal system in PCs *in vivo*.

### Putative glycolytic by-products LPA and serine contribute to the vulnerability of cerebellar PCs to neurodegeneration

Our findings on aberrant glycolytic activity in the ATG5 cKO PCs and their selective vulnerability under this condition prompted us to investigate whether accelerated glycolysis is causally implicated in the PC degeneration. Uncontrolled glycolysis can result in the production of methylglyoxal (MG) via the intermediate dihydroxyacetone phosphate (DHAP) (Fig. 6a). MG can readily react with proteins, lipids and nucleic acids to form advanced glycation end products (AGEs) that are implicated in various pathophysiological mechanisms, including neurodegeneration ^70^ (Fig. 6a, see also Fig. 2g). To test whether cKO of ATG5 leads to upregulation of MG-modified proteins in the cerebellar PCs, we analyzed their levels using immunohistochemistry. We found that the levels of MG-modified proteins were significantly increased in the PCs, but not in the molecular layer interneurons of 1-month and 3-month-old mice (Fig. 6b, Fig. S7a-c). The levels of MG-modified proteins were also upregulated in the whole ATG5 cKO cerebellar lysates compared to controls, but these changes did not reach significance (Fig. 6d,e).

**Figure 6.**
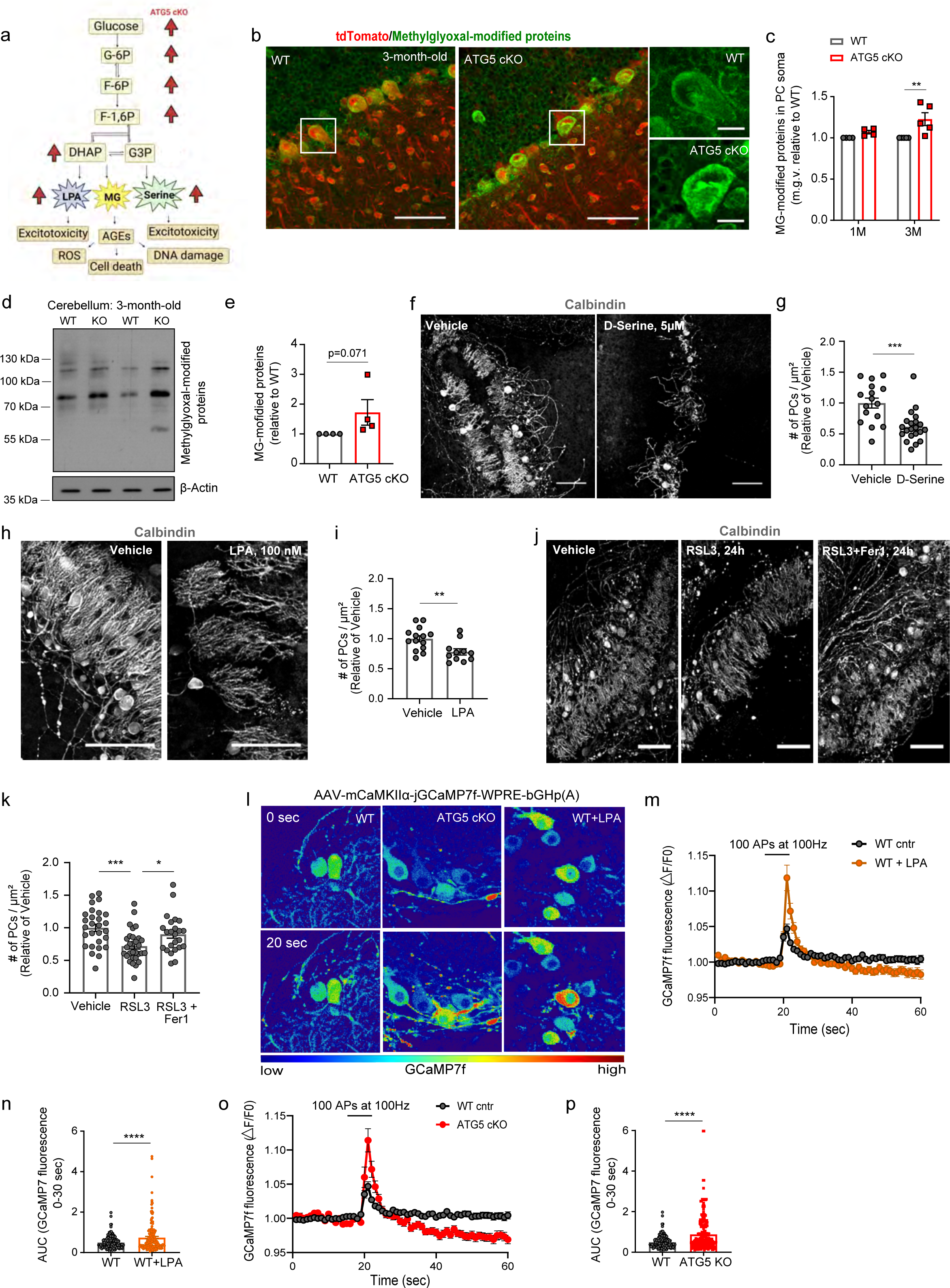
By-products of aerobic glycolysis cause neurodegeneration of cerebellar PCs. **a,** Schematic representation of glycolysis and its major by-products known to induce excitotoxicity in neurons. Red arrows indicate glycolytic by-products that were upregulated in the current study. **b**,**c**, Representative confocal images at the age of 3 months (b) and immunofluorescence analysis (c) of methylglyoxal(MG)-modified proteins in WT and ATG5 cKO PCs. WT and ATG5 cKO mice were expressing tdTomato as a reporter of Cre recombination. Scale bar, 50µm, inserts 10µm. Each dot represents one animal (N=5 for 3 months old animals and N=4 for 1 month old animals). Two-way ANOVA followed by Holm-Šidák multiple comparisons test (WT levels were set to 1, ATG5 cKO 1 month: 1.07 ± 0.021, ATG5 cKO 3 months: 1.227 ± 0.077; p=0.004). **d,e**, Representative immunoblot (d) and immunoblot analysis (e) of methylglyoxal-modified proteins in cerebellar lysates of WT and ATG5 cKO mice at the age of 3 months. Each dot represents one animal (N=4 per genotype). Protein levels were normalized to WT levels and set to 1. One-tailed unpaired T-test (ATG5 cKO: 1.723 ± 0.428; p=0.0712). **f,g,** Representative confocal images of PCs (f) and analysis of their density (g) in WT cerebellar organotypic slices at DIV21 cultured for 3 weeks either in control media (vehicle) or in the media containing 5µM of D-serine. Calbindin antibody was used as a marker of PCs. Scale bar: 100 µm. Each dot in (g) represents a single image (N=3 per genotype with 16 images for control and 21 images for D-serine treated slices). Two-tailed unpaired T-test (vehicle: 1 ± 0.08, D-serine treated: 0.62 ± 0.05; p=0.0003). **h,i,** Representative confocal images of PCs (h) and analysis of their density (i) in WT cerebellar organotypic slices at DIV21 cultured for 7 days either in control media (vehicle) or in the media containing 100 nM of LPA. Calbindin antibody was used as a marker of PCs. Scale bar: 100 µm. Each dot in (i) represents a single image (N=3 per genotype with 14 images for control and 11 images for LPA treated WT slices). Two-tailed unpaired T-test (vehicle: 1.00 ± 0.051, LPA-treated: 0.78 ± 0.051; p=0.007). **j,k,** Representative confocal images of PCs (j) and analysis of their density (k) in WT cerebellar organotypic slices at DIV21 treated for 24h either with DMSO (Vehicle), 1 µM RSL3 or 10 µM Fer-1 for 24h and immunostained for calbindin as PC marker. Scale bar: 100 µm. Each dot in (k) represents a single image (N=4 per genotype with 28 images for control, 30 images for RSL3 and 25 images for RSL3+Fer1 treated slices). One-way ANOVA followed by Holm-Šidák multiple comparisons test (vehicle: 1 ± 0.05, RSL3: 0.71 ± 0.04; p=0.0007 for comparison to vehicle, RSL3+Fer-1: 0.9 ± 0.059; p=0.037 for comparison to RSL3). **l**, Representative heat map of GCaMP7f fluorescence at baseline (0 sec) and after stimulation (20 sec) with 100 action potentials at 100 Hz in WT-untreated and WT-LPA-treated, and ATG5 cKO cerebellar organotypic slices at DIV21. WT slices were treated with 100 nM LPA for one week. **m,n,** GCAMP7f normalized fluorescence (m) and its area under the curve analysis (AUC, n) upon responses to 100 action potentials (APs) applied at 100Hz in untreated and LPA-treated WT PCs. Each dot in (n) represents a single PC (N=3 per genotype with 164 cells for WT and 166 cells for WT+LPA). Two-tailed unpaired T-test (untreated: 0.5 ± 0.02, LPA treated: 0.9 ± 0.18; p=0.0002). **o,p,** GCAMP7f normalized fluorescence (o) and its area under the curve analysis (AUC, p) upon responses to 100 action potentials (APs) applied at 100Hz in WT and ATG5 cKO PCs. Each dot in (p) represents a single PC (N=3 per genotype with 164 cells for WT and 142 cells for ATG5 cKO). Two-tailed unpaired T-test (WT: 0.5 ± 0.025, ATG5 cKO: 0.86 ± 0.069; p<0.0001). Data information: 1M and 3M indicate 1 and 3 months, respectively. Squares in Fig. 6b indicate regions magnified. All graphs show mean ± SEM. n.s.—non-significant; * indicates P ≤ 0.05; ** indicates P ≤ 0.01; *** indicates P ≤ 0.001; **** indicates P ≤ 0.0001.

In addition to MG, several other biosyhaventhesis pathways can branch off from glycolysis, including DHAP-mediated synthesis of lysophosphatidic acid (LPA)^71^ and the production of L-serine by 3-phosphoglycerate (3PG)^72^. L-serine can then be racemized into D-serine in the brain^73^. Since both LPA and serine (being a precursor of D-serine and glycine) can exert neurotoxic effects via augmenting neurotransmission^74-76^ and both metabolites were significantly upregulated in ATG5 cKO cerebellum (Fig. 6a, see also Fig. 2), we investigated their role in PCs survival. Treatment of WT cerebellar organotypic slice cultures with 5µM D-Serine and/or 100 nM LPA resulted in a substantial decrease in PC density (Fig. 6f-i). This cell loss was likely due to activation of ferroptosis since the loss of PCs could be triggered by treatment with the ferroptosis inducer RSL3 and prevented by their additional treatment with ferrostatin 1 (Fer1) (Fig. 6j,k), which is in line with the proteome data indicating activation of the ferroptosis pathway in the ATG5 cKO cerebellum (see Fig. 2e). Of note, the cell death of PCs upon autophagy deficiency was apoptosis-independent (Fig. S7d,e). The neurotoxic effect of LPA on PC death was accompanied by an increase in PC network excitation, as assessed through calcium imaging in cerebellar slice cultures transduced with an AAV encoding a GCaMP7f ^CamKII^α construct (CamKIIα is predominantly expressed in PCs but not granule cells or interneurons in the cerebellum) (Fig. 6l). The neuronal activity triggered by high-frequency bursts of action potentials (100Hz) was significantly elevated in WT PCs treated with LPA compared to controls (Fig. 6l-n). This phenotype was also observed in ATG5 cKO PCs (Fig. 6o,p). Of note, the baseline GCaMP7f fluorescence remained unaltered in both LPA-treated WT and ATG5 cKO PCs (Fig. S7f,g). Taken together, our findings suggest that glycolytic by-products LPA and serine contribute to PC degeneration and that accelerated glycolysis may be a key factor in the vulnerability of PCs observed under autophagy deficiency.

### GLUT2 deletion in ATG5 cKO mice mitigates PC neurodegeneration and improves their ataxic gait

Our data described thus far suggest that the loss of autophagy stabilizes GLUT2 levels in cerebellar PCs, a phenotype leading to uncontrolled glycolysis and subsequent PCs death. Therefore, we asked whether reducing GLUT2 (encoded by the *Slc2a2* gene) levels could prevent PC loss in ATG5 cKO mice. Hence, we generated double KO mice with *Slc2a2* gene inactivation by crossing previously described *Slc2a2* floxed mice ^77, 78^ with the ATG5 cKO mice (Fig. 7a, Fig. S8a). The resulting ATG5:GLUT2 cKO mice were born without abnormalities and exhibited weight gain similar to their WT littermates (Fig. S8b). We have previously reported that ATG5 cKO mice were leaner compared to control animals^47^. Intriguingly, the deletion of GLUT2 resulted in a modest improvement in weight gain in ATG5 cKO mice, but these changes did not reach significance (Fig. S8c). Conversely, the sole deletion of GLUT2 in GABAergic neurons was found to be inconsequential for weight regulation in mice, aligning with previous findings ^77^. We found that reducing GLUT2 protein levels (Fig. S8d-g) prevented excessive glucose uptake in ATG5:GLUT2 cKO PCs (Fig. 7b,c) and markedly downregulated the production of lactate compared with ATG5 cKO PCs (Fig. 7d-f). Glucose uptake in PCs lacking solely GLUT2 was not impaired, suggesting a compensation by other glucose transporters in the absence of GLUT2 (Fig. S8h). Additionally, levels of MG-modified proteins were also significantly reduced in ATG5:GLUT2 cKO mice (Fig. 7g,h), consistent with the hypothesis that stabilized GLUT2 in autophagy-deficient PCs contributes to increased formation of glycolytic by-products such as MG.

**Figure 7.**
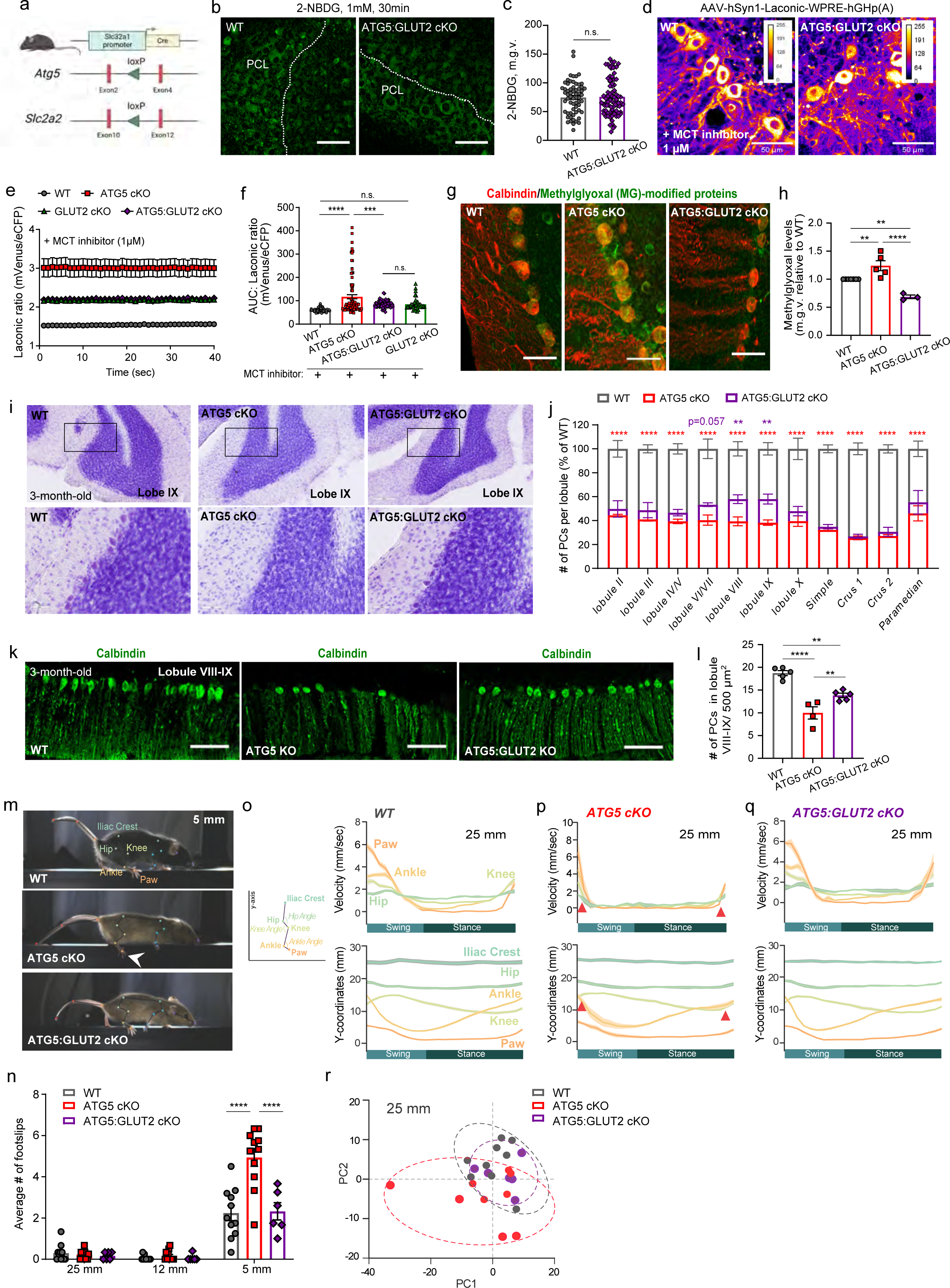
GLUT2 deletion in ATG5 cKO mice mitigates PC neurodegeneration and improves their ataxic gait. **a,** Schematic representation of the generation of ATG5cKO:GLUT2cKO mice with simultaneous deletion of ATG5 and GLUT2 in GABAergic neurons using the *Slc32a1* promoter. **b, c**, Representative confocal images of 2-NBDG uptake (b) and analysis of 2-NBDG levels (c) in WT and ATG5:GLUT2cKO PCs in cerebellar organotypic slice culture at DIV 21. Scale bar: 50 µm. PCL-PC layer. Each dot represents a single PC (for WT 62 cells and for DKO 83 cells from N=5). Two-tailed unpaired T-test (WT: 74 ± 3.6, DKO: 75.8 ± 3.5). **d**, FRET-based imaging of WT and ATG5:GLUT2 cKO cerebellar organotypic slice cultures transduced with AAV-hSyn1-Laconic. Scale bar: 50 µm. Slices were treated with 1 µM of MCT-1/2 inhibitor AR-C155858 to prevent extracellular lactate export. **e,** Ratiometric analysis of Laconic signal (mVenus/eCFP ratio) in WT (83 cells from N=6), ATG5 cKO (93 cells from N=5) ATG5:GLUT2cKO (69 cells from N=4) and GLUT2 cKO (35 cells from N=3) PCs. **f**, Area under the curve (AUC) analysis of Laconic signal (mVenus/eCFP ratio) in WT, ATG5 cKO, ATG5:GLUT2 cKO and GLUT2 cKO PCs. Each dot represents a single PC (83 cells from N=6 for WT, 93 cells from N=5 for ATG5 cKO, 69 cells from N=4 for ATG5:GLUT2cKO and 35 cells from N=3 for GLUT2 cKO). One-way ANOVA followed by Holm-Šidák multiple comparisons test (WT: 60.13 ± 0.84; ATG5 cKO: 117.3 ± 8.92; p<0.0001 in comparison to WT, DKO: 86.08 ± 1.77; p=0.0008 in comparison to ATG5 cKO, GLUT2 cKO: 85.18 ± 4.59). **g,h,** Representative confocal images (g) and immunofluorescence analysis (h) of methylglyoxal(MG)-modified proteins in WT, ATG5 cKO and ATG5:GLUT2 cKO PCs at the age of 3 months. Scale bar, 50µm. Each dot in (h) represents one animal (N=7 for WT, N=5 for KO and N=3 for DKO). One-way ANOVA followed by Holm-Šidák multiple comparisons test (levels were normalized to WT and set to 1, ATG5 cKO: 1.24 ± 0.08; p=0.0041 in comparison to WT and p<0.0001 in comparison to DKO, ATG5:GLUT2 cKO: 0.68 ± 0.03; p=0.0041 in comparison to ATG5 cKO). **i,** Representative Nissl-stained images of WT, ATG5 cKO and ATG5:GLUT2 cKO cerebellar 40µm-thick sections at the age of 3 months. Scale bar: 200µm, inserts 50 µm. **j**, Nissl staining-based analysis of averaged PC density per cerebellar lobule per 40µm section in 3-month-old WT, ATG5 cKO and ATG5 cKO:GLUT2 cKO cerebellum. (N=4 for WT and ATG5 cKO and N=5 for ATG5:GLUT2 cKO). Numbers in ATG5 cKO and ATG5 cKO:GLUT2 cKO were normalized to WT set to 100%. Two-Way ANOVA followed by Holm-Šidák multiple comparisons test (lobule VIII: p=0.0065, lobule IX: p=0.0040). Further statistics can be found in source data for Fig. 7. **k,l,** Representative confocal images of PCs immunostained for calbindin (k) and immunofluorescence-based analysis of their density (l) in WT, ATG5 cKO and ATG5:GLUT2 cKO cerebellum at the age of 3 months. Scale bar: 200 µm. Each dot in (l) represents one animal (N=5 for WT, N=4 for ATG5 cKO and N=5 for ATG5:GLUT2 cKO). One-Way ANOVA with Holm-Šidák multiple comparison test (WT: 18.74 ± 0.55, ATG5 cKO: 9.98 ± 1.35; p<0.0001 in comparison to WT and p=0.006 in comparison to DKO, DKO: 13.84 ± 0.49; p=0.0019 in comparison to WT). **m**, Individual frames from SIMI-based kinematic analysis videos showing control, ATG5 cKO and ATG5:GLUT2 cKO mice walking on the narrow beam (5mm). White arrowhead points to the slip in ATG5 cKO. **n**, Bar graph showing the average number of slips as mice crossing beams of different widths. Each dot in (n) represents one animal (N=11 for WT, N=11 for ATG5 cKO and N=6 for ATG5:GLUT2 cKO). Two-way ANOVA followed by Holm-Šidák multiple comparison (for 5 mm: WT: 2.24 ± 0.36, ATG5 cKO: 4.93 ± 0.43; p<0.0001 in comparison to WT and p<0.0001 in comparison to DKO, DKO: 2.32 ± 0.41). Further descriptive statistics can be found in source data for Fig.7. **o-q**, Line graphs showing the variation in hip, knee, ankle and hindpaw velocities (upper line plots) and coordination of the iliac crest, hip, knee, ankle and hindpaw on the y-axis (lower line plots) during a normalized step cycle in WT, ATG5cKO and ATG5:GLUT2 cKO mice walking on a 25mm-wide beam. Red arrowheads in (p) indicate the points in the step cycle at which the relative positions of the join differ in ATG5 cKOs compared to controls and ATG5:GLUT2 cKOs. N=9 mice for controls and ATG5 cKOs, N=6 for ATG5:GLUT2 cKOs. **r**, PCA analysis based on the y-coordinates, angles, and velocities per each normalized point (=25) of the step cycle as mice walked on a 25mm-wide beam. Individual mice are represented as colored circles, and dashed lines group mice of similar genotype. N=9 mice for controls and ATG5 cKOs, N=6 for ATG5:GLUT2 cKOs. Data information: Squares in Fig. 7i indicate regions magnified. All graphs show mean ± SEM. Data presented in Fig. 7o-q show mean as a filled dark line, and SEM as the shaded area around it. n.s.—non-significant; * indicates P ≤ 0.05; ** indicates P ≤ 0.01; *** indicates P ≤ 0.001; **** indicates P ≤ 0.0001.

To determine whether the rescue of glucose uptake, lactate production, and MG levels in ATG5:GLUT2 cKO mice has a protective impact on PC survival, we analyzed the number of PCs across various cerebellar regions in WT, ATG5 cKO and ATG5:GLUT2 cKO mice at 3 months of age. We found that the number of PCs was significantly reduced in all cerebellar lobules (in line with the analysis of calbindin-positive PCs in Fig. 1g) (Fig. 7i,j). Remarkably, GLUT2 reduction improved the survival of PCs in cerebellar lobules VI-IX, corresponding to the vermis region, by approximately 20% (Fig. 7j). This rescue phenotype was evident in both Nissl-stained cerebellum and cerebellar sections analyzed by immunohistochemistry using the calbindin antibody as a selective PCs marker (Fig. 7k,l).

Lesions in the cerebellar vermis cause truncal and gait ataxia in humans^79-81^, while cerebellar PC dysfunction or degeneration is the most frequent finding in animal models with ataxic symptoms^82^. To investigate whether the conditional deletion of ATG5 in mice leads to gait ataxia through PC loss, we evaluated their motor performance by examining the ability to traverse beams of varying widths, including wide (25 mm), regular (12 mm), and narrow (5 mm) beams. As the beam width decreases, mice typically adjust their intra- and interlimb kinematics to maintain body posture. Dysfunctions in the cerebellar circuitry can manifest as an inability to adapt limb kinematics, resulting in an increased number of slips. At 3 months of age, ATG5 cKO mice, although not impaired in their ability to stay on the rotarod (Fig. S8i), revealed a significant increase in footslips when crossing the narrow beam (5 mm width) compared to littermate controls (Fig. 7m,n, Video EV1). This phenotype was progressive, as 12-month-old ATG5 cKO mice could no longer able to cross the 5mm beam and showed a significant number of slips even on the wider 12mm beam (Fig. S8j). Strikingly, ATG5:GLUT2 cKO mice resembled the controls, displaying fewer footslips on the narrow beam (Fig. 7n, Video EV1). We further analyzed hindlimb movements by tracking and assessing kinematics with DeepLabCut and AutoGaitA (Fig. 7o, see also Fig. 7m). Examination of kinematic features revealed defects in intralimb coordination in 3-month-old ATG5 cKO mice, even on the wide (25 mm) beam (Fig. S9a, Video EV2,3). The relative positions of the knee and ankle, as well as the coordination of angle aperture and velocity differed in ATG5 cKOs (Fig 7p, Fig. S9b, as indicated by red arrowheads) compared to controls and were rescued by GLUT2 deletion (Fig.7q, Fig. S9b). Rescue of hindlimb kinematics was confirmed by comparing knee and hip angles, with significant differences observed in ATG5 cKO mice compared to ATG5:GLUT2 cKOs (Fig. S9c) and was further evident in Principal Component Analysis (PCA), where controls and ATG5;GLUT2 cKO mice were clustered together, while ATG5 cKO mice occupied a different position in the PCA space (Fig. 7r). Despite the footslip rescue, ATG5:GLUT2 cKO mice, similar to ATG5cKO animals, walked with a longer stance phase compared to controls, while swing duration remained unaltered (Fig. S9d,e). These kinematic alterations between genotypes persisted when the mice were required to cross narrower beams, albeit with smaller differences, likely due to the control and ATG5:GLUT2 cKO mice beginning to adapt their gait to the balance perturbations (Fig. S9f-y).

Taken together, our data show that conditional loss of ATG5 in mice induces a progressive gait ataxia, manifested by an increase in footslips and disturbed gait kinematics. This phenotype is effectively rescued by simultaneous deletion of GLUT2, with the double mutants showing a) fewer footslips on the narrow beam compared to ATG5 cKOs (Fig. 7n) and b) more physiological intralimb coordination when walking on a wide beam compared to ATG5 cKOs (Fig. 7o-r, S9a-c).

## Discussion

Loss of ATG5 function causes ataxia in both humans and flies ^83, 84^, yet the specific cellular pathway responsible for this phenotype remains elusive. Our study reveals a novel role of glycolytic metabolism under autophagy-deficient conditions as a possible pathological mechanism for gait ataxia. Through a comprehensive proteome and metabolome analysis, we identified a sustained dysregulation of the glycolysis pathway in the ATG5 cKO cerebellum. This metabolic shift is likely due to the marked upregulation of GLUT2 in autophagy-deficient PCs, resulting in increased glucose uptake and elevated production of glycolytic by-products, including MG. Excess MG has been associated with ataxia in mice, as evidenced by their uncoordinated beam walk ^85^, closely resembling the phenotype observed in ATG5 cKO mice in the current study. Moreover, MG is a major precursor of AGEs that are detrimental to neurons via the generation of reactive oxygen species (ROS) and inducing oxidative and inflammatory stress ^86^. Conversely, the reduction of AGEs has been proven to extend the lifespan in model organisms^87^. In line with this, we find that autophagy-deficient PCs undergo neurodegeneration, but their survival in the cerebellar vermis can be prolonged by downregulating GLUT2 levels, a manipulation that prevents the accumulation of MG-modified proteins. Our findings on selective rescue of PCs in the vermis suggest that the vermal PCs are particularly vulnerable to excessive glycolysis and/or that the loss of PCs in other cerebellar lobules is independent of glycolytic by-products. We observed that simultaneous deletion of GLUT2 in ATG5 cKO mice increased the number of vermal PCs by approximately 20% (with an overall rescue of PC number by up to 60%), which correlated with the recovery of gait dysfunction in these animals. These results align with clinical observations, as lesions or atrophy of the cerebellar vermis are associated with truncal ataxia in humans ^79^ and with a recent study in a mouse model of spinocerebellar ataxia 1 showing that PCs residing in the posterior vermis are particularly vulnerable to neurodegeneration ^88^. While our study provides compelling evidence for the rescue of ataxic gait in ATG5:GLUT2 cKO mice, we acknowledge that the specific contributions of other cell types, in addition to PCs, and the potential impact on the physiology of remaining PCs in ATG5 cKO mice cannot be definitively excluded at present. Gait abnormalities can also arise from disruptions in spinal cord circuits, as shown by spinal cord interneuron ablation studies ^89-91^. However, the phenotypes reported in these studies are more severe compared to our mutants, affecting the gait by causing limb hyperflexion and limiting locomotion even on wider walking surfaces. This suggests that the ataxic gait in ATG5 cKO mice may be predominantly caused by cerebellar dysfunctions. Recent single-cell RNA sequencing studies in the spinal cord confirm the absence of GLUT2 expression, reinforcing that the rescue is primary due to the restoration of the vermal glycolytic pathway ^92^. Although autophagy has previously been shown to be dispensable for the survival of motoneurons in the spinal cord^93^, the effects on spinal cord interneurons following ATG5 loss-of-function remain unknown. Nonetheless, our data emphasize the role of GLUT2-dependent glucose metabolism in gait disorders, urging further investigation into the specific neuronal circuits involved in future studies.

Autophagy can regulate cellular metabolism via three primary pathways: it can recycle amino acids and/or lipids ^18, 22^, maintain energy balance by regulating the quality and dynamics of the mitochondria^94^, and/or it can modulate the levels of key proteins in metabolic pathways ^28-31^. Here we unveil that cerebellar autophagy is not absolutely required for mitochondria quality control and/or recycling of amino acids but rather functions in PCs to regulate the degradation of the glucose transporter GLUT2. Our findings are in line with the previously described role of the autophagy-endolysosomal system in the regulation of GLUT1, GLUT2 and GLUT4 trafficking in non-neuronal cells^31, 68, 95 96^. To our knowledge, this is the first report on the regulatory role of autophagy in GLUT2 degradation in neurons, expanding the significance of autophagy beyond its canonical role in the turnover of damaged proteins and organelles and implicating it in the modulation of key metabolic pathways in the brain. Loss of autophagy via ATG5 deletion leads to a specific accumulation of GLUT2 in the PCs, driving excessive glycolytic metabolism. This finding introduces a new perspective on the role of autophagy in regulating GLUT2 levels in the cerebellum, distinguishing it from previous knowledge about GLUT1 and GLUT3 being autophagic substrates in the forebrain ^97^. How is GLUT2 targeted to autophagosomes? The absence of p62, ubiquitin and NBR1 foci in autophagy-deficient PCs questions the role of selective autophagy in cerebellar PCs and suggests that GLUT2 degradation may occur through bulk targeting to autophagosomes during starvation. This hypothesis aligns with a previous study showing that autophagosome formation in PCs is continuously stimulated by nutrient withdrawal ^98^ and our data revealing that GLUT2 can be downregulated in the cerebellum by starvation. Future work is required to elucidate the precise pathway targeting GLUT2 to autophagosome membranes.

Although the requirement of aerobic glycolysis in neurons has been debatable for decades^99-101^, several recent studies indicate that mature neurons require glucose uptake and glycolysis *in vivo* ^102-106^. Moderate aerobic glycolysis contributes to the antioxidant status of neurons and is essential to maintain synaptic function ^106^. Interestingly, the human cerebellum reveals significantly low aerobic glycolysis rates compared to other brain regions^102, 107^. How these regional differences in aerobic glycolysis are achieved is currently unknown. Here, we propose that the glycolytic activity in the cerebellum is attenuated by the functional involvement of autophagy. Several lines of evidence indicate that autophagy is a key negative regulator of cerebellar glycolysis. First, using two unbiased approaches (i.e., untargeted and targeted LC-MS/MS metabolome analysis), we show that several glycolytic metabolites are highly and significantly upregulated in the ATG5 cKO cerebellum (Figs. 2-3). Second, using ^13^C-glucose as a tracer, we reveal that loss of ATG5 significantly facilitates the glucose flux into glycolysis, but not the TCA pathway (Fig. 3g, Fig. S4). Third, ATG5-deficient PCs, but not cerebellar interneurons, produce significantly more lactate and non-mitochondria-derived ATP (Fig. 4). Finally, ATG5-deficient PCs accumulate MG, a reactive carbonyl species generated endogenously during glycolysis (Fig. 6). The heightened glycolytic activity within PCs lacking autophagic machinery is one of the central revelations of our study. Previously, the loss of *Atg7* in mouse embryonic fibroblasts has been reported to cause an increase in glycolytic activity, as a compensation due to accumulation of damaged mitochondria upon loss of autophagy ^108^. The increased glucose uptake and subsequent acceleration of glycolysis in ATG5 cKO PCs reported here can be indicative of a concerted effort to meet the energy demands of PCs in the absence of autophagy-mediated quality control mechanisms. However, due to the absence of features of quality control dysfunction, i.e. mitophagy and protein aggregations, we favor the hypothesis that the glycolytic cascade in autophagy-deficient PCs extends beyond ATP production. Our study suggests that during cerebellar maturation, autophagy keeps aerobic glycolysis low by degrading GLUT2 and attenuates the effects of glycolytic by-products, including MG, serine and LPA on cerebellar PC physiology. The observed metabolic transformation in the absence of autophagy underscores the pivotal role of glucose metabolism in neuronal vulnerability. Notably, we find that the cerebellum exhibits more pronounced alterations in glucose metabolism compared to the cortex, emphasizing cell-type-specific responses to autophagy loss. The hypothesis of disturbed glucose metabolism as a vulnerability factor in cerebellar neurodegeneration is also in line with cerebellar gray matter reduction in patients with type 2 diabetes mellitus ^109^ and with evidence of gait alterations in older adults with diabetes ^110, 111^, a period of life when autophagy is also reduced^112^. Our study also challenges conventional explanations, such as defective mitochondrial clearance or impaired oxidative phosphorylation in neurodegenerative movement disorders such as Parkinson’s disease ^113^, thereby urging further exploration into the role of excessive glycolysis in neuronal cell death. The metabolic rewiring observed in autophagy-deficient PCs echoes similar metabolic adaptations reported in other neurodegenerative disorders^39, 114-117^, suggesting a common thread in the nexus between autophagy and cellular metabolism.

FDG-PET signals are routinely employed in human imaging studies to assess neuronal activity and/or aerobic glycolysis efficiency in the brain ^37, 38^. Our study reports a decrease in the FDG-PET signal in the ATG5cKO cerebellum, which at first glance is inconsistent with the concomitant observation of increased glucose metabolism at the individual ATG5 cKO PC level. We attribute this apparent discrepancy to the inherent limitations of PET imaging. While PET provides valuable temporal *in vivo* resolution for metabolic changes, it is constrained spatially. Notably, accelerated glycolysis in our model manifests as early as 1 month of age. However, the FDG-PET study was conducted in 3-month-old mice, a stage when PC density is significantly reduced due to autophagy deficiency. Considering that each individual PC receives up to 200,000 synapses^118^, the observed loss of PCs would result in a substantial reduction in their presynaptic inputs. This aligns with the results of our SV2A PET study and supports the hypothesis that the combined effects of PC loss and/or reduced synaptic inputs (compartments consuming a substantial amount of glucose in neurons^119^), together with the limited spatial resolution of FDG-PET, explain the lack of a hypermetabolism phenotype in the ATG5 cKO cerebellum in PET studies.

Previous research utilizing mouse models with brain-confined loss of *Atg5* and *Atg7* genes through *Nes* promoter-driven Cre recombinase expression underscored the importance of autophagy in maintaining neuronal health^9, 17^. However, the broad expression of *Nes*-Cre in all neuronal progenitors poses challenges in attributing neurodegeneration solely to autophagy defects in neurons. In contrast, our study utilized the *Atg5*flox:*Slc32a1*-Cre mouse line^47^, enabling a more refined exploration by directly comparing autophagy-deficient GABAergic neurons across the different brain regions in the same animal. This approach unveiled a distinct region-specific vulnerability of the same neuronal subclass, elucidating the progressive degeneration of PCs and the emergence of ataxic phenotypes. Our data on the resistance of cerebellar interneurons to ATG5 deletion are consistent with a recent study^120^ and our previous work ^46, 47^ showing that forebrain interneurons in the same mouse model can tolerate autophagy dysfunction. Remarkably, the vulnerability of PCs to autophagy impairment stands out for its lack of typical apoptotic cell death and intracellular aggregates seen in other autophagy-related neurodegenerative models. This unique characteristic propelled us to explore alternative cell death mechanisms, with our data hinting at the potential involvement of ferroptosis - a cell death pathway recently implicated in several neurodegenerative disorders, including ataxia^121-123^. Whether and how exactly the excessive glycolysis causes the death of PCs by ferroptosis should be further investigated in future studies.

In conclusion, our study provides the first evidence of the intricate relationship between autophagy, glycolysis, and the susceptibility of cerebellar PCs to neurodegeneration. The implications extend beyond the immediate purview of cerebellar pathophysiology, offering a broader conceptual framework for understanding neurodegenerative processes. Targeting the identified metabolic vulnerabilities, including glycolytic dysregulation and GLUT2 perturbations, may offer novel strategies to ameliorate the effects of ataxia associated with ATG5 loss-of-function. The delineation of these molecular intricacies opens avenues for targeted therapeutic interventions aimed at modulating cellular metabolism to ameliorate the impact of neurodegenerative conditions at their metabolic nexus.

## Methods

### Mouse models

All animal experiments were approved and performed according to the regulations of the LANUV (Landesamt für Natur, Umwelt und Verbraucherschutz Nordrhein-Westfalen) guidelines. Mice were maintained in a pathogen-free environment in ventilated polycarbonate cages. Animals were housed in groups of five animals per cage with constant temperature and humidity at 12h/12h light/dark cycles. Food and water were provided *ad libitum*. *Atg5*flox:*Slc32a1*-Cre:tdTomato (ATG5 cKO) mice were described previously ^47^. To create ATG5:GLUT2 cKO mice, mice with floxed exon 3 of the *Atg5* gene were crossed with mice, where the exon 11 of the *Slc2a2* gene was flanked by loxP (kindly provided by Dr. Bernhard Thorens, University of Lausanne, Switzerland)^77, 78^. Table S1 indicates genotyping primers used to genotype the animals in the current study.

### PET imaging

Animals were anesthetized with isoflurane in O2/air 3:7 (induction 5%, maintenance 1.5–2.0 %), and a catheter for tracer injection ([^18^F]UCB-H in 3-month-old mice, [^18^F]MNI1126 in 12-month-old mice) was inserted into the lateral tail vein. After placement in the tail vein, the needle tip was secured with adhesive tape, and the syringe with the tracer solution was attached to the other end of the tube. Mice were placed on an animal holder (medres® GmbH, Cologne, Germany) and fixed with a tooth bar in a respiratory mask. Body temperature was maintained at 37 °C using a feedback-controlled warming system. Eyes were protected from drying out by the application of eye and nose ointment (Bepanthen, Bayer). A PET scan in list mode was conducted using a Focus 220 micro PET scanner (CTI-Siemens, Erlangen, Germany) with a resolution at the center of field of view of 1.4 mm. Data acquisition started with intravenous tracer injection (activity: 6–12 MBq in 125 µl) and lasted for 40 min. This was followed by a 10 min transmission scan using a 57Co point source for attenuation correction. After the scan was finished, the catheter was removed and the mice were returned to their home cage. For [^18^F]FDG-PET, animals were briefly anesthetized (see above) and 9-12 MBq [^18^F]FDG in 125 µl were injected intraperitoneally. The mice were then placed in a solitary cage where they spent the following 35 min awake. Subsequently, they were anesthetized again and scanned with a resolution at the center of field of view of 1.4 mm for 30 min. This protocol takes advantage of metabolic trapping of [^18^F]FDG^124^, which allows awake tracer uptake and subsequent scanning under anesthesia.

After full 3D rebinning, summed images were reconstructed using an iterative OSEM3D/MAP procedure^125^, resulting in voxel sizes of 0.47 × 0.47 × 0.80 mm. For all further processing of the images including statistics, the software VINCI 5.21 for MacOS X (Max Planck Institute for Metabolism Research, Cologne, Germany) was used. Images were co-registered and intensity-normalized to the thalamus. To this end, an elliptical volume of interest (VOI) of 7.2 mm^3^ (40 voxels) was placed inside the thalamus. Each image was divided by the mean value of the thalamus VOI, resulting in the “standardized uptake value ratio” (SUVR_Thal_). No further postprocessing (e.g. Gauss filtering or spatial morphing) was done. For comparison of ATG5 cKO versus controls, a voxel-wise t-test was performed for each tracer using VINCI 5.21 for MacOS X. The equation for the unpaired t-test (i.e. difference between the two group means divided by the pooled standard error of both groups) was executed manually in VINCI. The resulting t-maps were corrected for multiple comparisons using a threshold-free cluster enhancement (TFCE) procedure described in detail in ^126^. The TFCE procedure was implemented as a Python script in VINCI. For final thresholding at p<0.05, a permutation test with 10,000 permutations was performed in RStudio 1.0 for MaxOS X using the SUVR_Thal_ values of the voxel with the highest TFCE value. The 95% quantile was calculated, and the corresponding TFCE level was used as the lower threshold of the t_TFCE_-map. The resulting t_TFCE_-maps were displayed in voxel view in shades of red (ATG5 cKO>controls) or blue (ATG5 cKO<controls) and projected onto a C57BL/6 T2-weighted MRI template.

### Primary cerebellar culture

Mice were sacrificed at the postnatal day (p) 7-9 and brains were collected in ice-cold PBS containing 13 mM glucose, 300 mg BSA (fatty acid-free) and 1.5 mM MgSO_4_ (solution A). Cerebellum was isolated and cut into 700 µm thick-pieces using a McIlwain tissue chopper (Cavey Laboratory Engineering Co. LTD). Pieces were incubated in solution A containing 0.25 mg trypsin for 15 min at 37°C. To stop trypsinization, the solution A containing 0.5 mg Sobyean Trypsin Inhibitor (SBTI) and 600U DNase was added twice the volume as solution A with trypsin and centrifuged at 1000 g for 1 min at 4°C. Afterwards supernatant was removed, and the cell suspension was dissolved in 1 ml DMEM-based (Gibco) growth medium containing 0.4 % B-27 (50x Gibco), 19 mM KCl (Roth), 14 mM HEPES (Roth), 10 mM glucose (Sigma), 1 mM sodium pyruvate (100x Gibco), 0.25% L-GlutMax (100x Gibco) 1 % P/S (10000 Units/ml Penicillin, 10,000 µg/ml Streptomycin, Thermo Fisher Sci) and mechanistically dissociated using fire-polished glass pipettes. Dissociated cells were carefully layered on top of EBSS (Gibco), containing 4% (w/v) bovine serum albumin (BSA); 3 mM MgSO_4_ and centrifuged at 1500 g for 5 min at 4°C. Supernatant was discarded, cell pellets were dissolved in growth medium containing 10% FBS and cell density was determined using a Neubauer counting chamber. Cells were then plated at a concentration of 750 000 per well and fed with growth medium after 1 h. After 24 h and 7 days half of the medium was replaced by fresh growth medium containing 4 µM AraC. Cerebellar neurons were kept for 17 days in vitro (DIV 17) under constant conditions at 37°C/5% CO_2._ If necessary, neurons were transduced with AAVs at DIV3 (0.2 µl per ml medium).

### Seahorse assay

Metabolic measurements were carried out in standard 96-well Seahorse microplates on a Seahorse XF96 analyzer, using the Mito Stress Test and Glycolysis Stress Test (Agilent). Cerebellar cells were plated in 96-well Seahorse XF Cell Culture Microplates plates and analyzed at DIV17. For the Mito Stress Test, cells were washed 1 h before the experiment and incubated with Seahorse XF Base Medium supplemented with 1 mM pyruvate, 2 mM glutamine and 10 mM glucose in a 37°C non-CO_2_ incubator. Oxygen consumption rate (OCR) was first measured upon basal respiration i.e within the first three rounds without the injection of inhibitors. Afterwards, 1.5 µM oligomycin (complex V inhibitor) was injected to measure ATP production, followed by 2 µM FCCP (uncoupling of oxygen consumption from ATP production) injection to measure maximal respiration capacity and 0.5 µM rotenone/antimycin A (inhibitors of complex I and III) combined with Hoechst was added at the end to measure non-mitochondrial respiration. Afterwards, cell density was calculated based on the Hoechst signal via a microplate reader and used for normalization of OCR values per cell density. For Glycolysis Stress Test, cells were washed and incubated with Seahorse XF Base Medium supplemented with 2 mM glutamine in a 37°C non-CO_2_ incubator 1 h prior to the experiment. The extracellular acidifECAR was first measured without any injection to analyze the non-glycolytic acidification caused by other processes than glycolysis. The first injection of 10 mM glucose indicates the rate of glycolysis under basal conditions. Afterwards, 1.5 µM oligomycin (complex V inhibitor) was injected to measure the maximum glycolytic capacity of the cells, followed by 50 mM 2-deoxy-glucose (2-DG, an inhibitor of glycolysis by inhibiting hexokinase) combined with a Hoechst injection to measure the glycolytic reserve of the cells. Afterwards, cell density was calculated based on the Hoechst signal via a microplate reader and used for normalization of ECAR values per cell density.

### Live imaging in cerebellar organotypic slice culture

Mice were sacrificed at p6-9 and brains were collected in ice-cold Hanks’ Balanced Salt Solution (HBSS, Thermo Fisher Scientific). Cerebellum was isolated and cut into 300 µm-thick sagittal sections (McIlwain tissue chopper, Cavey Laboratory Engineering Co. LTD). Sections were separated and washed three times in pre-warmed (37°C) HBSS by carefully transferring slices with glass pipettes. Slices were then transferred onto membrane inserts with prewarmed organotypic slice culture (OTC) cultured MEM-based (Sigma) medium containing 0.00125% ascorbic acid (Roth), 10 mM D-glucose (Sigma), 1 mM GlutaMAXX^TM^ (100x Gibco), 20% (v/v) horse serum (Gibco), 0.01 mg/ml insulin (Thermo Fisher Sci), 14.4 mM NaCl (Roth), 1% P/S (Thermo Fisher Sci). Medium was replaced every second day and slices were kept for 21 days (DIV21) under constant conditions at 37°C/5% CO_2_. OTC were either used for live imaging or for immunohistochemistry (IHC) and therefore fixed in 4% PFA for 30 min at room temperature. For IHC slices at DIV21 were fixed in 4% PFA for 1 h at RT washed 3 times with PBS and then processed as fixed brain sections (described below in detail).

Viral transduction of OTCs was done on DIV1 by adding 1 µl of adeno-associated virus (AAV) (see Table S2 for the list of used AVVs in the current study) on top of each single slice. Live imaging was performed at days in vitro (DIV) 21 in carbogenated OTC imaging medium (dH_2_O; 2 mM CaCl_2_; 10 mM D-glucose; 3 mM KCl; 1 mM MgCl_2_; 136 mM NaCl; 24 mM NaHCO_3_; 1.25 mM NaH_2_HPO_4_) upon constant perfusion. For live imaging, the TCS SP8 MP-OPO (Leica) multiphoton confocal microscope was used, equipped with a 20x/0.75 multi-immersion objective. Scans were performed at a 512x512 pixel resolution, bidirectional recording of 1 frame per second. Videos of 40 s length were recorded if not stated otherwise. For ratiometric FRET measurements, the donor eCFP was excited at 448 nm, and emission of both eCFP and mVenus were captured by individual detectors between 450 nm and 500 nm for eCFP and between 520 and 570 nm for mVenus. Recordings were analyzed using ImageJ (Fiji) by measuring fluorescence intensity of eCFP and mVenus individually in the region of interest. Backgrounds were subtracted for each fluorophore and the ratio of mVenus/ eCFP was calculated and used as a readout of either ATP or lactate levels. For lactate imaging, the OTC medium also contained 1 µM MCT-inhibitor AR-C155858 (Tocris). For GCamP7f imaging, videos with a length of 60 s were recorded (1 frame/s). After 20 s of baseline recordings, slices were stimulated once with 100 APs at 100 Hz using an RC-47FSLP stimulation chamber (Warner Instruments). Recordings were then analyzed using ImageJ (Fiji). First background was subtracted and then baseline fluorescence was measured by averaging fluorescence of the first 19 frames before stimulation. Fluorescent values of each frame were then normalized to baseline and plotted over time. In mitoTimer live imaging, a fluorescent timer, or DsRed1-E5, shifts over time from green (excitation and emission maxima = 483 nm and 500 nm) to red (excitation and emission maxima = 558 nm and 583 nm) as the protein matures. The mitochondrial targeting sequence in this construct is derived from cytochrome *c* oxidase subunit VIII^127^. For mt-Keima live imaging, single plane images were taken at a 1024x1024 pixel resolution with bidirectional recording. Keima is a pH-dependent fluorescent protein that has a single emission peak around 620 nm with dual excitation wavelengths that is tagged with a mitochondrial target sequence of COX VIII. In an alkaline environment mt Keima is excited at 458 nm and in acidic environment at 561. Images were analyzed using ImageJ (Fiji) by measuring fluorescence intensity individually in the region of interest. Backgrounds were subtracted for each fluorophore and the ratio of 561/ 456 was calculated and used as a readout of mitophagy.

### NBDG uptake in cerebellar organotypic slice culture

For 2-NBDG uptake, cerebellar OTCs were incubated in OTC medium without glucose supplemented with 1 mM 2-NBDG (Biomol) for 30 min at 37°C in the dark. OTCs were washed twice with OTC imaging medium before life imaging. Videos were recorded for 40 s (1 frame/s) multiphoton confocal microscope TCS SP8 MP-OPO (Leica) equipped with 20x/0.75 multi immersion objective in OTC imaging medium. Recordings were then analyzed using ImageJ (Fiji). Fluorescence of each frame was analyzed after background subtraction. Average fluorescence values over time were used for further analysis.

### Preparation of acute slices, starvation and chloroquine treatment

Mice were sacrificed at the age of three months via cervical dislocation, and brains were isolated. Cortex and cerebellum were separated and cut with a vibratome (Leica) into 150 μm horizontal sections in ice-cold, carbogen-saturated (95% O_2_ and 5% CO_2_) low-Ca^2+^ osmolarity-adjusted artificial cerebrospinal fluid (ACSF: 125 mM NaCl, 2.5 mM KCl, 1.25 mM NaH_2_PO_4_ (0,4 M), 25 mM NaHCO_3_, 10 mM Glucose, 0.5 mM CaCl_2_, 3.5 mM MgCl_2,_ pH=7.4). Cerebellar and cortical acute slices were incubated in control medium (OTC medium), control medium with 40 µM chloroquine (Sigma) or ACSF for 6h at 37°C and 5% CO_2_. Afterwards samples were shock frozen in liquid nitrogen and stored at -80°C until further processing via immunoblotting.

### Untargeted metabolomics

Untargeted metabolomics was performed on 10-20 mg of fresh-frozen cerebellum from one and three months-old animals. For metabolite extraction 25 µl of metabolite extraction solution (LC-MS grade methanol 50%, LC-MS grade acetonitrile 30%, Ultrapure water 20%, Valine-d8 final concentration 5 µM) was added per mg tissue and tissue was lysed in a homogenizer (2x 30s cycle at 6 000 rpm at 4°C. Tissue extracts were transferred into fresh tubes, shaken in a thermomix (Eppendorf) at full speed for 15 min at 4°C and centrifugated at full speed for 20 min at 4°C. Equal amounts of each sample were used for further analysis. Metabolites were separated chromatographically utilizing a Millipore SeQuant ZIC-pHILIC analytical column (5 µm, 2.1 × 150 mm), complemented with a 2.1 × 20 mm guard column (both with 5 mm particle size), and a binary solvent system was employed. Solvent A consisted of 20 mM ammonium carbonate and 0.05% ammonium hydroxide, while Solvent B was pure acetonitrile. The column oven was maintained at 40°C and the autosampler tray at 4 °C. The chromatographic gradient flowed at a rate of 0.200 mL/min and was programmed as follows: 0–2 min at 80% B, a linear gradient from 80% B to 20% B between 2-17 min, a swift change from 20% B to 80% B between 17-17.1 min, and a hold at 80% B from 17.1-23 min. Samples were injected in a randomized order with a volume of 5 µl each. A pooled quality control (QC) sample, created by mixing equal parts of all individual samples, was analyzed at regular intervals in between the test samples. Metabolites were quantified using a Vanquish Horizon UHPLC connected to an Orbitrap Exploris 240 mass spectrometer (both provided by Thermo Fisher Scientific), with measurements facilitated by a heated electrospray ionization source. Spray voltages were configured at +3.5kV/-2.8kV, the RF lens value at 70, the heated capillary temperature at 320 °C, and the auxiliary gas heater at 280 °C. Sheath, aux, and sweep gas flow rates were set at 40, 15, and 0, respectively. For MS1 scans, the mass range was m/z=70-900, with a standard AGC target and auto-set maximum injection time. Experimental samples’ data acquisition employed full scan mode with polarity switching, at an Orbitrap resolution of 120,000. The AcquireX Deep Scan workflow was used for untargeted metabolite identification, applying an iterative data-dependent acquisition strategy with multiple pooled sample injections. The setup included a full scan resolution of 60,000, fragmentation resolution of 30,000, and a fragmentation intensity threshold of 5.0e3. Dynamic exclusion was activated after one occurrence, with a 10s exclusion duration and a 5ppm mass tolerance. The isolation window was 1.2 m/z, with normalized HCD collision energies in stepped mode at 30, 50, 150. Mild trapping was enabled. Metabolite identification was conducted using the Compound Discoverer software (v 3.2, Thermo Fisher Scientific), with criteria including a precursor ion m/z within 5 ppm of the predicted theoretical mass according to the chemical formula, fragment ions within 5 ppm to an internal spectral library of authentic compound standards (analyzed with the same ddMS2 method) with a minimum match score of 70, and retention time within 5% of a purified standard under the same chromatographic conditions. Peak area integration and chromatogram review were performed using Tracefinder software (v 5.0, Thermo Fisher Scientific). Each detected metabolite’s peak area was normalized against the total ion count (TIC) of the respective sample to account for variations from sample handling to analysis, serving as variables for in-depth statistical analysis. Pathway analysis of untargeted metabolomics was performed using MetaboAnalyst 5.0 (https://www.metaboanalyst.ca/home.xhtml).

### 13C6-glucose tracing of amino acids and glycolysis- and TCA cycle-metabolites using liquid chromatography-high-resolution mass spectrometry (LC-HRS-MS)

Cerebellar acute slices prepared from 3-month-old mice were incubated in ACSF containing 10 mM ^12^C_6_ glucose (unlabeled) or 1 mM ^13^C_6_-labeled glucose for 1h at 37°C and 5% CO_2_. The Sliceswere transferred to 2 mL Eppendorf tubes, washed once in 75 mM ammonium carbonate pH=7.4 and the fresh weight of samples was determined. To each of the samples a 5 mm stainless steel metal ball was added before snap-freezing them in liquid nitrogen. The frozen samples were pulverized for 1 min using a TissueLyser (Qiagen) set to 25 Hz for 1 min.

Immediately after homogenization 1 ml of −20 °C mixture of UPLC-grade acetonitrile:methanol:water (2:2:1 (v:v:v)), containing 375 nmol U-^13^C^15^N amino acid mix (Cambridge isotopes MSK_A2-1.2), each 150 ng/mL of ^13^C_10_ ATP, ^15^N_5_ ADP and ^13^C_10_^15^N_5_ AMP (Sigma) and 200 ng/mL of citric acid ^2^H_4_ (Sigma), was added. After addition of the extraction buffer, the samples were immediately vortexed for 10 seconds before incubating them for additional 30 min at 4 °C on an orbital shaker. To remove the insoluble material the metal balls were removed from each sample using a magnet and the tubes were centrifuged for 10 min at 21.000 x *g* at 4°C. The cleared supernatant was transferred to a fresh 1.5 mL Eppendorf tube and immediately concentrated to complete dryness in a speed vacuum concentrator set to 20°C.

For the LC-MS analysis the dried samples were resuspended in 150 µL of ice-cold UPLC-grade water, vortexed thoroughly and centrifuged for 5 min at 21.000 x *g* at 4°C. The cleared supernatant was transferred to a fresh 1.5 mL Eppendorf tube. For the LC-MS analysis of amino acids 50 µL of the available 150 µL extract were mixed with 25 µl of 100 mM sodium carbonate (Sigma), followed by the addition of 25 µl 2% [v/v] benzoylchloride (Sigma) in acetonitrile (UPC/MS-grade, Biosove, Valkenswaard, Netherlands). The samples were thoroughly mixed and kept at 20°C until analysis. For the LC-HRMS analysis, 1 µl of the derivatized sample was injected onto a 100 x 2.1 mm HSS T3 UPLC column (Waters). The flow rate was set to 400 µl/min using a binary buffer system consisting of buffer A (10 mM ammonium formate (Sigma), 0.15% [v/v] formic acid (Sigma) in UPC-MS-grade water (Biosove, Valkenswaard, Netherlands). Buffer B consisted of acetonitrile (IPC-MS grade, Biosove, Valkenswaard, Netherlands) 0.1% [v/v] formic acid (Sigma). The column temperature was set to 40°C, while the LC gradient was: 0% B at 0 min, 0-15% B 0-4.1min; 15-17% B 4.1 – 4.5 min; 17-55% B 4.5-11 min; 55-70% B 11 – 11.5 min, 70-100% B 11.5 - 13 min; B 100% 13 - 14 min; 100-0% B 14 -14.1 min; 0% B 14.1-19 min; 0% B. The mass spectrometer (Q-Exactive Plus) was operating in positive ionization mode recording the mass range m/z 100-1000. The heated ESI source settings of the mass spectrometer were: Spray voltage 3.5 kV, capillary temperature 300°C, sheath gas flow 60 AU, aux gas flow 20 AU at 330°C and the sweep gas was set to 2 AU. The RF-lens was set to a value of 60. For the analysis of anionic metabolites of the glycolysis and the TCA cycle, the remaining 100 µl were transferred to polypropylene autosampler vials (Chromatography Accessories Trott, Germany) before performing Anion exchange chromatography (Integrion, Thermo Fisher Scientific), coupled to high resolution mass spectrometry (Q-Exactive HF, Thermo Fisher Scientific) analysis. In brief, 5 µL of the resuspended polar metabolite extract were injected in push-partial mode, using an overfill factor of 1, onto a Dionex IonPac AS11-HC column (2 mm × 250 mm, 4 μm particle size, Thermo Fisher Scientific) equipped with a Dionex IonPac AG11-HC guard column (2 mm × 50 mm, 4 μm, Thermo Fisher Scientific). The column temperature was held at 30°C, while the auto sampler temperature was set to 6°C. A potassium hydroxide gradient was generated using a potassium hydroxide cartridge (Eluent Generator, Thermo Scientific), which was supplied with deionized water (Milli-Q IQ 7000, Millipore). The metabolite separation was carried at a flow rate of 380 µL/min, applying the following gradient conditions: 0-3 min, 10 mM KOH; 3-12 min, 10−50 mM KOH; 12-19 min, 50- 100 mM KOH; 19-22 min, 100 mM KOH, 22-23 min, 100-10 mM KOH. The column was re-equilibrated at 10 mM for 3 min. For the analysis of metabolic pool sizes the eluting compounds were detected in negative ion mode using full scan measurements in the mass range m/z 77 – 770. The heated electrospray ionization (ESI) source settings of the mass spectrometer were: Spray voltage 3.2 kV, capillary temperature was set to 300°C, sheath gas flow 50 AU, aux gas flow 20 AU at a temperature of 330°C and a sweep gas glow of 2 AU. The S-lens was set to a value of 50. The LC-MS data analysis of the amine- and the glycolysis- and TCA-compounds was performed using the TraceFinder software (Version 5.1, Thermo Fisher Scientific). The identity of each compound was validated by authentic reference compounds, which were measured at the beginning and the end of the sequence. For data analysis the area of all detectable isotopologues mass peaks of every required compound were extracted and integrated using a mass accuracy <3 ppm and a retention time (RT) tolerance of <0.05 min as compared to the independently measured reference compounds. If no independent ^12^C experiments were carried out, where the pool size was determined from the obtained peak area of the ^12^C monoisotopologue, the pool size determination was carried out by summing up the peak areas of all detectable isotopologues per compound. These areas were then normalized, as performed for un-traced ^12^C experiments, to the internal standards, which were added to the extraction buffer, followed by a normalization to the fresh weight of the analyzed samples. The relative isotope distribution of each isotopologue was calculated from the proportion of the peak area of each isotopologue towards the sum of all detectable isotopologues.

### Targeted metabolomics

Levels of amino acids, glycolysis intermediates and TCA cycle intermediates were analyzed in fresh-frozen cerebellum from 1 and 3 month-old mice. Intermediates (organic acids, sugar phosphates) of the glycolysis and the TCA cycle in mouse cerebellar tissue were determined by Anion-Exchange Chromatography coupled to Electrospray Ionization High-Resolution Mass Spectrometry (IC-ESI-HRMS) using a procedure previously described ^128^ (Aravamudhan et al. 2021). Approximately 50 mg of snap-frozen mouse tissue were homogenized in ice-cold acetonitrile/methanol/water 2:2:1 (v/v/v) (1 mg / 10 µl) using the Precellys 24 Homogenisator (Peqlab) at 6,400 rpm for two times 10 seconds with a 5-second pause. 100 µl of homogenate were mixed with further 225 µl of acetonitrile/methanol/water 2:2:1 (v/v/v) and 25 µl of a mixture of isotope-labeled internal standards in Milli-Q water (5 µM 13C6-D-glucose-6-phosphate (13C6-G6P) and 5 µM D4-succinic acid (D4-SUC), both Eurisotop). After thorough mixing and centrifugation (16,100 RCF, 5 min, 4 °C), 300 µl of supernatant were dried under reduced pressure. The residue was resolved in 100 µl of Milli-Q water, transferred to autoinjector vials and immediately measured. IC-HRMS analysis was performed using a Dionex Integrion RFIC system (Thermo Scientific) coupled to a Q Exactive HF quadrupole-orbitrap mass spectrometer (Thermo Scientific) as previously described ^128, 129^. The exact m/z traces of the internal standards and endogenous metabolites were extracted and integrated using Skyline 21.2.0.369 (open-source). Endogenous metabolites were quantified by normalizing their peak areas to those of the internal standards: D4-SUC was used for organic acids, and 13C6-G6P for sugar phosphates. Amino acids and GABA of mouse cortex and cerebellum tissue were derivatized with benzoyl chloride and quantified by Liquid Chromatography coupled to Electrospray Ionization Tandem Mass Spectrometry (LC-ESI-MS/MS) using a procedure previously described (Wrong et al., 2016)^129^. The homogenates prepared for the analysis of sugar phosphates and organic acids (see above) were centrifuged (16,100 RCF, 5 min, 4 °C). 20 µl of the supernatant were mixed with 10 μl of the MassChrom Internal Standard Mixture Amino acids and Acylcarnitines from Dried Blood (Chromsystems), reconstituted in 5 ml water/methanol 2:1 (v/v), and 10 μl of a 10 µM solution of d6-GABA (Sigma-Aldrich) in Milli-Q water. Endogenous and isotope-labeled amino acids and GABA were derivatized by adding 10 µl of freshly prepared 2 % benzoyl chloride in acetonitrile and 10 µl of 100 mM sodium carbonate in water and thorough mixing (Wrong et al., 2016). After addition of 40 µl of Milli-Q water and centrifugation (16,100 RCF, 5 min, 4 °C), 80 µl of the supernatant were transferred to autoinjector vials and immediately measured. LC-MS/MS analysis was performed by CECAD Lipidomics Facility using a Nexera X2 UHPLC System (Shimadzu) coupled to a QTRAP 6500 triple quadrupole/linear ion trap mass spectrometer (SCIEX) as previously described ^128-130^. The LC chromatogram peaks of benzoylated endogenous amino acids and GABA, and internal standards were integrated using the MultiQuant 3.0.2 software (SCIEX). The peak areas of the benzoylated metabolites were normalized to those of the internal standards.

### Proteomics

Total proteome analysis was performed on fresh frozen cerebellum from 1 and 3 months-old animals. Samples were prepared by in-solution digestion and all solution were provided by the proteomics facility. Samples were lysed in Urea lysis buffer (50 mM TEAB; 8M Urea; 50x Protease inhibitor), centrifuged for 15 min at 20 000 g and protein concentration was assessed using Bradford assay (Sigma). 50 µg of protein for each sample was transferred in fresh tubes, incubated with 5 mM Dithiothreitol (DTT) for 1h at 25°C and then incubated with 40 mM Chloroacetamide (CAA) for 30 min in the dark. Afterwards samples were incubated with Lysyl Endopeptidase (LysC) at an enzyme:substrate ratio of 1:75 for 4h at 25°C. Samples were then diluted with 50 mM Triethylammoniumbicarbonate (TEAB) to achieve a final concentrate of 2 M Urea before overnight incubation in Trypsin (1:75 ratio) at 25°C. The next day samples were acidified with formic acid (final concentration 1%). Peptides were extracted and purified using Stagetips. First StageTips were equilibrated with washes once in methanol, buffer B (80% acetonitril; 0.1% (v/v) formic acid) and twice in buffer A (dH2O; 0.1% (v/v) formic acid), each time followed by centrifugations at 2 600 rpm for 1-2 min. For peptide purification, samples were centrifugated at 13 000 rpm for 5 min and the loaded onto StageTips. Samples were centrifugated at 2 600 rpm for 5 min, StageTips were washed with buffer A and centrifugated at 2 600 for 3 min. Then StageTips were washed twice with buffer B and each time centrifugated at 2 600 for 3 min. Eluted peptides were dried in vacuo, stored at 4°C and provided to proteomics facility for further processing. Samples were analyzed by the CECAD Proteomics Facility on an Orbitrap Exploris 480 (Thermo Scientific, granted by the German Research Foundation under INST 1856/71-1 FUGG) mass spectrometer equipped with a FAIMSpro differential ion mobility device that was coupled to an UltiMate 3000 (Thermo Scientific). Samples were loaded onto a precolumn (Acclaim 5µm PepMap 300 µ Cartridge) for 2 min at 15 ul flow before reverse-flushed onto an in-house packed analytical column (30 cm length, 75 µm inner diameter, filled with 2.7 µm Poroshell EC120 C18, Agilent). Peptides were chromatographically separated at a constant flow rate of 300 nL/min and the following gradient: initial 6% B (0.1% formic acid in 80 % acetonitrile), up to 32% B in 72 min, up to 55% B within 7.0 min and up to 95% solvent B within 2.0 min, followed by column wash with 95% solvent B and re-equilibration to initial condition. The FAIMS pro was operated at -50V compensation voltage and electrode temperatures of 99.5 °C for the inner and 85 °C for the outer electrode. For the Gas-phase fractionated library, a pool generated from all samples was analyzed in six individual runs covering the range from 400 m/z to 1000 m/z in 100 m/z increments. For each run, MS1 was acquired at 60k resolution with a maximum injection time of 98 ms and an AGC target of 100%. MS2 spectra were acquired at 30k resolution with a maximum injection time of 60 ms. Spectra were acquired in staggered 4 m/z windows, resulting in nominal 2 m/z windows after deconvolution using ProteoWizard (Chambers, 2012). For the samples, MS1 scans were acquired from 399 m/z to 1001 m/z at 15k resolution. Maximum injection time was set to 22 ms and the AGC target to 100%. MS2 scans ranged from 400 m/z to 1000 m/z and were acquired at 15 k resolution with a maximum injection time of 22 ms and an AGC target of 100%. DIA scans covering the precursor range from 400 - 1000 m/z and were acquired in 60 x 10 m/z windows with an overlap of 1 m/z. All scans were stored as centroid. The gas-phase fractionated library was built in DIA-NN 1.8.1 (Demichev 2020) using A Swissprot mouse canonical database (UP589, downloaded 04/01/22) with settings matching acquisition parameters. Samples were analyzed in DIA-NN 1.8.1 as well using the previously generated library and identical database. DIA-NN was run with the additional command line prompts “—report-lib-info” and “—relaxed-prot-inf”. Further output settings were: filtered at 0.01 FDR, N-terminal methionine excision enabled, maximum number of missed cleavages set to 1, min peptide length set to 7, max peptide length set to 30, min precursor m/z set to 400, max precursor m/z set to 1000, cysteine carbamidomethylation enabled as a fixed modification. Afterwards, DIA-NN output was further filtered on library q-value and global q-value <= 0.01 and at least two unique peptides per protein using R (4.1.3). Finally, LFQ values calculated using the DIA-NN R-package. Afterwards, analysis of results was performed in Perseus 1.6.15^131^. Pathway analysis of proteomic approaches was performed using ShinyGO (South Dakota State University; Ge, Jung and Yao, 2020). Venn diagram analysis was done using Venny2.1 (Oliveros, J.C. (2007-2015) Venny. An interactive tool for comparing lists with Venn’s diagrams. https://bioinfogp.cnb.csic.es/tools/venny/index.html).

### Immunohistochemical analysis of brain sections

Mice were euthanized at 1, 3 and 12 months of age by an overdose of 1.2% ketamine, and 0.16% xylazine in PBS (i.p., 100µL per 10 g body weight) and transcardial perfusion was performed as previously described ^132^. Brains were carefully dissected and postfixed in 4% PFA (pH=7.4) overnight before being processed for immunohistochemistry as previously described^132^. Corresponding horizontal free-floating 40 µm sections from WT and KO littermates were washed three times in PBS (2 × 5 min each). Sections were blocked with 10% normal goat serum (NGS) or 10% normal donkey serum (NDS) in 0.5 % Triton-X in PBS (PB-T) for 1 h at room temperature (RT). Primary antibodies (see Appendix Table S3) were incubated in sections in 3% NGS and 0.3% PBS-T for 48h at 4°C. Sections were washed three times 10 min in 0.3% PBS-T before incubation in fluorescence-labeled secondary antibodies in 3% NGS/NDS and 0.3% PBS-T for 2 h at RT protected from light (see Appendix Table S3). The sections were imaged at a Leica TCS SP8 (Leica Microsystems) equipped with PL Apo 10x/0.40 CS2, PL Apo 40x/0.85 CORR CS and PL APO 63x/1.40 oil CS2 and Leica Stellaris 5 confocal microscopes (Leica Microsystems) equipped with a HCPL FLUOTAR 10x/0.30, HC PL APO 20x/0.75 CS2, APO 63×/1.32 FLYC CORR CS2 objective and a continuous excitation white light laser. Images were visualized using Fiji (ImageJ). Samples were scanned at a resolution of 1,024 × 1,024 pixels with bidirectional recording and stacks of 20-30 optical sections were taken. Mean gray values were used as fluorescence readout of protein levels after background subtraction. If otherwise stated, fluorescence levels were normalized to WT levels and used for statistical analysis. For colocalization analysis, samples were scanned at a resolution of 1,024 × 1,024 pixels using Leica Stellaris 5 confocal microscope equipped with Plan-Apochromat 63×/1.30 GLYC objective. Stacks of 20-30 optical sections were taken with a fixed section size of 330 nm. Reconstructions and colocalization analysis were performed with Amira Software 2020.2 (Thermo Fisher Scientific). For each channel isosurfaces were extracted and positive colocalizations were defined when distance between voxels were smaller than 300 nm. The surface of distant voxels were mapped using the surface distance tool and plotted as histogram and as a color-coded image. For cresyl-violet staining 40 μm horizontal sections were mounted on gelatin-coated “super-frost plus” glass slides and dried overnight. Sections were hydrophylized in water for 1 min before incubation in cresyl-violet solution for 7 min. Sections were then washed three times in water for 2 min before being dehydrated in ascending ethanol solutions (70%, 80%, 90%, 96%, 100%) for 2 min in each solution. Afterwards, sections were incubated in xylene for at least 2 min and then covered with mounting solution Entellan ® (Merck). Images were acquired at a S360 Hamamatsu slide scanner using a 40x objective. Aperio ImageScope viewing software (Leica, version 12.4.3.5008) was used to analyze the number of Purkinje cell somata for individual cerebellar lobes.

### Electron microscopy

For electron microscopy (EM) animals at the age of 3 months were perfused using 2% Formaldehyde (Science Services, München, Germany) and 2.5% Glutaraldehyde (Merck, Darmstadt, Germany) in 0.1M Cacodylate buffer. Brains were removed and fixed overnight in the same fixative. 40 µm horizontal vibratome sections were prepared and the region of interest was extracted using a biopsy punch. Postfixation was applied using 1% Osmiumtetroxid (Science Services, München, Germany) and 1% Potassium hexacyanoferrat (Merck, Darmstadt, Germany) for 30 min at 4°C. After 3x5min wash with ddH2O, samples were dehydrated using ascending ethanol series (50%, 70%, 90%, 100%) for 10 min each. Infiltration was carried out with a mixture of 50% Epon/ethanol for 1h, 70% Epon/ethanol for 2h and overnight with pure Epon (Merck, Darmstadt, Germany). After fresh Epon for 4h, vibratome sections were mounted onto empty epon blocks and covered with Aclar foil. After 48h hardening at 60°C, Aclar foil was removed and samples were trimmed to the region of interest. Ultrathin sections (70nm) were cut using a diamond knife (Science Service, München, Germany) on an UC6 ultramicrotome (Leica, Wetzlar, Germany) and collected onto pioloform coated slot grids. Poststaining was performed with 1.5 % uranyl acetate (Agar Scientific, Stansted, United Kingdom) for 15 min and Reynolds lead citrate (Roth, Karlsruhe, Germany) solution for 3 min. Images were acquired using a JEM-2100 Plus Transmission Electron Microscope (JEOL, Tokio, Japan) operating at 80kV equipped with a OneView 4K camera (Gatan, Pleasanton, USA). EM quantifications were carried out manually on acquired images (10 000K) of Purkinje cell somata. Number of mitochondria were counted and normalized to the total area of the cell. The total area of single mitochondria and the total length (perimeter) of the corresponding cristae was analyzed. Cristae length was then normalized to the total area of mitochondria.

### Immunoblotting analysis

Mice were sacrificed at the age of one, 3 and/or 12 months via cervical dislocation. Brains were isolated, individual brain regions were separated, shock frozen in liquid nitrogen and stored at - 80°C until further use. Samples were homogenized in RIPA buffer (dH_2_O; 1% Igepal; 150 nM NaCl; 0.1% SDS; 0.5% SOD; 50 mM Tris) containing protease inhibitor (Roche) and phosphatase inhibitor (ThermoScientific) using a Wheaton otter-Elvehjem Tissue Grinder. Afterwards samples were sonicated (10 pulses), incubated on ice for 45 min and centrifuged at 13 000 rpm for 15 min at 4°C. Supernatants were transferred into fresh tubes and proteins concentrations were assessed using Bradford assay (Sigma). Sample were then mixed with 4x SDS buffer and boiled for 95°C for 5 min. Depending on the assay, 10-20 μg protein per sample were loaded onto SDS-page gels for protein separation and then transferred onto nitrocellulose or methanol-activated PVDF membranes via full-wet transfer assay (BioRad) or semi-wet transfer. Protein transfer was confirmed by Ponceau S staining. Membranes were blocked in 5% milk or bovine serum albumin (BSA) in TBS containing 1% Tween (TBS-T) for 1h at RT followed by primary antibody incubation in TBS over night at 4°C. Afterwards, membranes were washed three times with TBS-T for 10 min and then incubated with HRP-tagged secondary antibodies for 1h at RT followed by three washes in TBS-T for 10 min at RT. Protein levels were visualized using ECL-based autoradiography film system (Super RX-N, Fujifilm) or ChemiDoc^TM^ Imaging system (BioRad) and analyzed using Gel Analyzer plugin from ImageJ (Fiji). Protein levels were always first normalized to loading control and to the WT control.

### Surface biotinylation assay on acute cerebellar slices

Cerebellar acute slices were washed in 37°C pre-warmed ACSF (see above) for 30 min before incubation in ACSF containing 1 mg/ml EZ-Link™ Sulfo-NHS-SS-Biotin (Thermo Fisher Sci) for 1h at 4°C. Afterwards sections were incubated in ice-cold ACSF containing 100 mM glycine to stop the biotinylation reaction for 30 min at 4°C and washed three times with ice-cold ACSF. Sections were transferred into fresh tubes, lysed in lysis buffer, sonicated, incubated on ice for 45 min and centrifuged at 13 000 rpm for 15 min at 4°C. Supernatants were transferred into fresh tubes and proteins concentrations were assessed using Bradford assay (Sigma). Supernatant was used as input fractions. 300 µg of protein of each sample were incubated with appropriate amount of prewashed Streptavidin beads (Thermo Fisher Sci) over night at 4°C. The next day samples were centrifuged at 13 000 rpm for 30s at 4°C and the supernatant was transferred into fresh tubes. Samples account for non-biotinylated fraction and were stored at -80°C until further use for immunoblotting. Beads were washed three times with ice cold lysis buffer (dH_2_O; 1% Igepal; 150 nM NaCl; 0.1% SDS; 0.5% SOD; 50 mM Tris) and once with 50 mM HCl. Samples were centrifugated at 13 000 rpm for 30s at 4°C and supernatant was discarded. For protein elution, 40µl of 2x SDS+ DTT were added and samples were incubated for 5 min at 95°C. After samples were equilibrated to RT, supernatant was collected and stored at -80°C until further use. Collected samples contained biotinylated proteins. Input and non-biotinylated fraction were equally mixed with 2x SDS+DTT according to their initial volume and incubated for 5min at 95°C. For immunoblotting the same amount of all samples were loaded (input, biotinylated, and non-biotinylated fraction).

### AAV production

pAAV-mDlx-eGFP-ATG5 was generated from pEGFP-C1-hApg5 (a gift from Noboru Mizushima, Addgene plasmid # 22952) and pAAV-mDlx-GFP-Fishell-1 (a gift from Gordon Fishell, Addgene plasmid # 83900; http://n2t.net/addgene:83900; RRID:Addgene_83900) by restriction subcloning. EGFP-hATG5 was amplified using forward (TCGCCACCATGGTGAG) and reverse (GCAGGGGTGGCGCGCCTTCAATCTGTTGGCTGTGGG) primers. PCR product and pAAV-mDlx-GFP-Fishell-1 were digested, using SgsI-FD and NcoI-FD (both ThermoFisher Scientific), and ligated with T4 DNA Ligase (NEB). The plasmids pAdDeltaF6 (Addgene plasmid #112867; http://n2t.net/addgene:112867; RRID:Addgene_112867) and pAAV2/rh10 (Addgene plasmid # 112866; http://n2t.net/addgene:112866; RRID:Addgene_112866) were a gift from James M. Wilson. All DNA constructs were confirmed by Sanger sequencing (Eurofins).

Recombinant AAV2/rh10 particles were prepared in HEK293T cells (DSMZ no. ACC 635) by transfecting either pAAV-mDlx-eGFP or pAAV-mDlx-eGFP-ATG5 together with pAdDeltaF6 and pAAV2/rh10. Viral particles were precipitated with PEG/NaCl and cleared with chloroform extraction^133^. AAVs were purified by adapting scalable anion-exchange chromatography strategies^134, 135^. Cleared AAVs were concentrated roughly 20-fold with pre-washed (PBS + 0.001% (v/v) Poloxamer 188, Sigma Aldrich) 100 kDa Amicon filters (Merch/Millipore) and diluted 10-fold in buffer A (10 mM bis-tris-propane pH 9.0, 1 mM MgCl2). AAVs were applied at a flow-rate of 3 mL/min to a self-packed 1 mL column (POROSTM HQ 50 µm strong anion exchange resin, Thermo Fisher Scientific), which was equilibrated in buffer A. After injection, the column was rinsed with 20 column volumes buffer A, washed with 20 column volumes 4 % buffer B (10 mM bis-tris-propane pH 9.0, 1 mM MgCl2, 1 M NaCl). AAVs were eluted with 35 % buffer B. Eluted fractions were concentrated and buffer exchanged to PBS + 0.001% (v/v) Poloxamer 188 using 100 kDa Amicon filters. Purity of viral preparations were assessed with SDS-PAGE/Colloidal Commassie staining and AAV titers determined using Gel green® (Biotium)^136^.

### Quantitative RT-PCR analysis

Mice were sacrificed at the age of 3 months via cervical dislocation. Brains were isolated, individual brain regions were separated, shock frozen in liquid nitrogen and stored at -80°C until further use. RNA isolation of cerebellar tissue was performed using Trizol Fisher Scientific). 2 µg of total RNA was used for reverse transcription using the high Capacity cDNA Reverse Transcription Kit (Applied Biosystems) following the manufacturer’s instructions. qPCR was performed with PowerUp^TM^ SYBR^TM^ Green Master Mix (Applied Biosystems) in a StepOnePlus^TM^ Real-Time PCR System (Applied Biosystems). Relative expression of gene transcripts was assessed using the 2-Δ ΔCt method. The following forward (F) and reverse (R) primers were used for qRT-PCR: F5′-CCAGTACATTGCGGACTTCCTT-3′ and R5′-CTTTCCTTTGGTTTCTGGAACTTT-3′ for *Glut2* and F5′-CCTCCAAGGAGTAAGAAACCC-3′ and R5′-GGGTGCAGCGAACTTTATTG-3′ for *Gapdh* used as a reference gene.

### Stereotactic injections

For injections, mice were weighed and anesthetized with a mixture of Ketamine (100 mg/kg)/ Xylazine (20 mg/kg)/ Acepromazine (3 mg/kg) and placed in a stereotactic frame when fully asleep. An eye creme was applied on the eyes in order to prevent drying of the corneas. A local painkiller was injected subcutaneously at the operation field. The animal was fixed in a stereotactic frame provided with a heating pad (David Kopf Instruments). The skin was opened, and the skull was cleaned using NaCl. Point injection (AP from Bregma: -5.61 mm; ML: 00 mm; DV: -1 mm) was identified using Bregma and Lambda for navigation. Subsequently, a circular craniotomy was performed with a micro drill. A microinjection syringe (WPI) equipped with a 34g bevelled needle (WPI) was filled with corresponding AAV (Table S2) and lowered to the depth of -1 mm to inject 300 nl of AAV at a speed of 100 nL/min. After the injection was complete, the needle was kept in place for 3 min and slowly retracted over 1 min. The incision was closed with sutures (Ethicon). During and after the surgery the mice body temperature was kept at 37°C via a heating pad. Mice were injected with 5% glucose-saline solution (100 μL/10g) after the surgery and with carprofen (5 mg/kg) 24 h and 48 h after the surgery. Physical conditions of the animals were monitored daily to improve their welfare. Three weeks after virus injection, animals were perfused with Ringer and 4% PFA (see section above for IHC).

### Behavioral analysis

To assess motor coordination and balance alterations, WT, ATG5 cKO and ATG5:GLUT2 cKO mice were tasked to cross beams of 1,3 meter in length and different widths (5 mm, 12 mm, and 25 mm). Mice crossed a different beam each day, for three consecutive days, starting from the widest one (25 mm). For each mouse, three to five trials per beam were collected. Mice were recorded using eight high-speed cameras (mV Blue Cougar XD; 200 frames/second) strategically positioned around the beam (3D SIMI Motion). Multiple camera views were analyzed to count the number of slips, which were then averaged across trials per individual mouse. For the 2D kinematic reconstructions of the hindlimb, we used the camera perpendicular to the beam. The iliac crest, hip, knee, ankle and hindpaw coordinates were tracked using DeepLabCut^137^ in markerless animals. We also used DeepLabCut to label the beam position, and establish the baseline for the vertical axis. The initiation of swing, the end of swing and the end of stance were manually annotated for each step. Steps resulting in footslips were excluded from the kinematic analysis. Please note that ATG5 cKOs displayed a higher number of slips on the narrowest beam (5 mm), therefore only three mice could be analyzed in this task. We used AutoGaitA to integrate the manual annotations of individual steps with the DeepLabCut tracked coordinates. AutoGaitA is a python-based algorithm that a) normalizes the co Mice at were euthanizedordinate across steps, b) calculates angles, velocities and accelerations, c) averages trials per mouse, d) averages mice per genotype, e) compares data across genotypes. The group values were analyzed using ANOVA or PCA analysis as indicated in the Figure Legend, performed in either AutoGaitA or GraphPad Prism.

### Statistical analysis

Sample sizes were not chosen based on pre-specified effect size. Instead, multiple independent experiments were carried out using several samples replicates, as detailed in the figure legends. For all experiments, there was enough statistical power to detect the corresponding effect size. Statistical analyses were done *ex vivo* on cell values (indicated by data points) and/or mice for *in vivo* from at least three independent experiments (indicated by “N”, biological replicates). Excel (Microsoft, USA) and GraphPad Prism version 9 (GraphPad Software, Inc., USA) were used for statistical analysis and result illustration (unless otherwise stated). Statistical analysis of normalized data between the two groups was performed using a one-tailed unpaired Student’s t-test. Statistical significance between two groups for normally distributed non-normalized data was evaluated with a two-tailed unpaired Student’s t-test. For comparisons between two groups in a set of multiple data, multiple T-Test with posthoc correction using two-stage linear step-up procedure of Benjamini, Krieger and Yekutieli was performed. For comparison between more than two groups, one-way ANOVA followed by a Holm-Sidak post hoc test was applied. Two-way ANOVA (or mixed model ANOVA, if the number of N was not equal between conditions) was used for the comparison of two groups and two independent variables, followed by Holm-Sidak post hoc test for multiple comparisons. P values of less than 0.05 were considered statistically significant. Data are reported as mean values ± standard error of the mean (SEM).

## Data availability

All data needed to evaluate the conclusions in the paper are present in the paper and/or the source data. Proteome data of all experiments are deposited in the database PRIDE and accessible for public after publishing. Source data are provided with this paper. Additional data related to this paper may be requested from the corresponding author.

## Acknowledgements

We thank S. Müller and Dr. M. Schröter for their expert assistance. We are indebted to Dr. C. Jüngst (CECAD Imaging Facility), Dr. S. Müller, Dr. J.-W. Lackmann (CECAD Proteomic Facility) and Dr. S. Brodesser (CECAD Metabolomics/Lipidomics) for their help and expert assistance. We are very gratefully to Prof. J. Brüning (Max Planck for Metabolism Research) for making us aware of *Slc2a2* flox mice in B. Thorens lab. We thank Prof. J. Vogt for providing LPA and Prof. S von Karstedt for providing RSL3. Funding: The work of NLK is funded by the Deutsche Forschungsgemeinschaft (DFG, German Research Foundation): EXC 2030–390661388, KO 5091/4-1, DFG-431549029–SFB 1451, DFG-233886668-GRK1960 and DFG-411422114 - GRK 2550.

## Author contributions

Conceptualization: NLK

Methodology: JT, GG, FL, PG, HE, NLK

Investigation: JT, MT, EY, LI, IK, HE, PG

Visualization: JT, GG, NLK

Supervision: GG, GS, FC, AD, BN, NLK

Writing—original draft: JT & NLK

Writing—review & editing: JT & NLK

