## Supplementary Material for "Autophagy regulator ATG5 preserves cerebellar function by safeguarding its glycolytic activity"

**This PDF file includes:**

Supplementary figure legends 1-9

Appendix Table S1

Appendix Table S2

Appendix Table S3

Appendix Table S4

Legends for Table EV1, Table EV2, Table EV3, Table EV4

Videos EV1-3

**Supplementary figure legends.**

**Figure S1. Deregulated autophagy without absence of p62 and NBR1 foci in ATG5 cKO PCs.**

**a,** Protein levels of ATG5~ATG12 conjugate are significantly decreased in cortical brain lysates from 1-month and 3-month-old ATG5 cKO mice compared to the WT set to 1. N=4 for each genotype and condition. Two-way ANOVA followed by Holm-Šidák multiple comparisons (1 month ATG5 cKO: 0.35 ± 0.04; p<0.0001, 3 months ATG5 cKO: 0.16 ± 0.23; p<0.0001).

**b**, LC3 lipidation at autophagosome membranes (LC3II/LC3I ratio) is significantly decreased in cortical brain lysates from 1-month and 3-month-old ATG5 cKO mice compared to the WT set to1. N=4 for each genotype and condition. Two-way ANOVA followed by Holm-Šidák multiple comparisons (1 month ATG5 cKO: 0.47 ± 0.1; p<0.0001, 3 months ATG5 cKO: 0.39 ± 0.04; p<0.0001).

**c**, Protein levels of p62 are significantly increased in cortical brain lysates from 1-month and 3-month-old ATG5 cKO mice compared to the WT set to1. N=3 for each genotype at 1 month of age and N=4 for each genotype at 3 months of age. Two-way ANOVA followed by Holm-Šidák multiple comparisons (1 month ATG5 cKO: 2.65 ± 0.51; p=0.017, 3 months ATG5 cKO: 3.16 ± 0.54; p<0.0032).

**d,** Protein levels of ATG5~ATG12 conjugate are significantly decreased in cerebellar brain lysates from 1-month and 3-month-old ATG5 cKO mice compared to the WT set to1. N=4 for each genotype and condition. Two-way ANOVA followed by Holm-Šidák multiple comparisons (1 month ATG5 cKO: 0.14 ± 0.05; p<0.0001, 3 months ATG5 cKO: 0.29 ± 0.05; p<0.0001).

**e**, LC3 lipidation at autophagosome membranes (LC3II/LC3I ratio) is significantly decreased in cerebellar brain lysates from 1-month and 3-month-old ATG5 cKO mice compared to the WT set to1. N=4 for each genotype and condition. Two-way ANOVA followed by Holm-Šidák multiple comparisons (1 month ATG5 cKO: 0.8 ± 0.09; p=0.038, 3 months ATG5 cKO: 0.67 ± 0.06; p=0.0038).

**f**, Protein levels of p62 are significantly increased in cerebellar brain lysates from 1-month and 3-month-old ATG5 cKO mice compared to WT set to1. N=4 for each genotype and condition. Two-way ANOVA followed by Holm-Šidák multiple comparisons (1 month ATG5 cKO: 2.22 ± 0.18; p<0.0001, 3 months ATG5 cKO: 1.78 ± 0.14; p=0.0003).

**g**, Protein levels of GFAP are unaltered in cortical brain lysates from 3-month-old ATG5 cKO mice compared to the WT set to1. N=4 for each genotype and condition. Two-way ANOVA followed by Holm-Šidák multiple comparisons (1 month ATG5 cKO: 0.9 ± 0.1, 3 months ATG5 cKO: 1.09 ± 0.11).

**h**, Protein levels of GFAP are increased in cerebellar brain lysates from 3-month-old ATG5 cKO mice compared to WT set to1. N=4 for each genotype and condition. Two-way ANOVA followed by Holm-Šidák multiple comparisons (1 month ATG5 cKO: 1.02 ± 0.07, 3 months ATG5 cKO: 1.32 ± 0.08; p=0.0028).

**i,** Protein levels of calbindin are significantly decreased in cerebellar brain lysates from 3-month-old ATG5 cKO mice compared to the WT set to 1. N=4 for each genotype and condition. Two-way ANOVA followed by Holm-Šidák multiple comparisons (1 month ATG5 cKO: 0.84 ± 0.09, 3 months ATG5 cKO: 0.306 ± 0.106; p<0.0001).

**j,** Representative confocal images of the cerebellum from WT and ATG5 cKO mice at the age of 3 and 12 months expressing tdTomato as a reporter of Cre recombination, immunostained for parvalbumin. Scale bar: 200 µm.

**k,l**, Representative confocal images of the cerebellum and cortex from WT mice at the age of 1 month (k) and WT/ATG5 cKO mice at 3 months (l) expressing tdTomato as a reporter of Cre recombination, immunostained for p62. Scale bar: 25 µm.

**m,n**, Representative confocal images of the cerebellum and cortex from ATG5 cKO mice at the age of 1 month (m) and WT/ATG5 cKO mice at 3 months (n) expressing tdTomato as a reporter of Cre recombination, immunostained for NBR1. Scale bar: 25 µm.

Data information: 1M and 3M indicate 1 and 3 months, respectively. S All graphs show mean ± SEM. n.s.—non-significant; * indicates P ≤ 0.05; ** indicates P ≤ 0.01; *** indicates P ≤ 0.001; **** indicates P ≤ 0.0001.

**Figure S2. Unperturbed mitochondria function in the ATG5 cKO cerebellum.**

**a**, EM-based analysis of mitochondria cristae perimeter normalized to mitochondria area in WT and ATG5 cKO PC somata. N=4 for each genotype. Two-tailed unpaired T-test (WT: 18.4 ± 1.14, ATG5 cKO: 21.41 ± 0.46).

**b,** Analysis of oxygen consumption rate (OCR) during the Seahorse XF Cell Mito Stress test in WT and ATG5 cKO primary cerebellar cells (DIV17) (N=5 cultures per genotype). Descriptive statistics in source data for Fig. S2.

**c**. Representative images of mitochondria-localized MitoKeima (mt-mKeima)-expressing PCs in WT and ATG5 cKO cerebellar organotypic slice culture at DIV21. Scale bar: 20µm (1 µm insert).

**d**, mt-mKeima-based fluorescence image analysis of mitophagy (red fluorescence divided by green fluorescence) in PC soma in WT and ATG5 cKO cerebellar organotypic slice cultures. Each dot represents one animal with N=4 (WT)/3(cKO) per genotype. Two-tailed unpaired T-test (WT: 0.59 ± 0.02, ATG5 cKO: 0.58 ± 0.059).

**e**, Protein levels of BNIP3 are unaltered in cerebellar brain lysates from 3-month-old ATG5 cKO mice compared to the WT set to 1. N=3 for each genotype. One-tailed unpaired t-test (ATG5 cKO: 1.03 ± 0.07).

**f**, Representative confocal images of the cerebellum from ATG5 cKO mice at the age of 3 month immunostained for TIM23 as mitochondria marker and co-immunostained for BNIP3. Scale bar: 25 µm.

Data information: All graphs show mean ± SEM. n.s.—non-significant; * indicates P ≤ 0.05; ** indicates P ≤ 0.01; *** indicates P ≤ 0.001; **** indicates P ≤ 0.0001.

**Figure S3. Proteomic analysis of deregulated pathways in 1- and 3-month-old ATG5 cKO cerebellum**

**a**,**b**, ShinyGO v0.741-based GO analysis of Biological Processes (BP)-enriched terms in the cerebellar proteome (p < 0.05 and log2 fold change of < > 0.25) at 1 month (a) and 3 months of age (b).

**c,d**, ShinyGO v0.741-based GO analysis of Cellular Component (CC)-enriched terms in the cerebellar proteome (p < 0.05 and log2 fold change of < > 0.25) at 1 month (c) and 3 months of age (d).

**e,f**, Venn diagram of commonly upregulated (e) and downregulated (f) proteins (p < 0.05 and log2 fold change of < > 0.25) in the 1- and 3 months-old ATG5 cKO cerebellum.

**g,h,** ShinyGO v0.741-based GO analysis of Cellular Component (CC)- (g) and Biological Processes (BP) (h) enriched terms in the commonly upregulated and downregulated cerebellar proteome (p < 0.05 and log2 fold change of < > 0.25) at 1 month and 3 months of age.

**Figure S4. The decrease in TCA cycle intermediates is secondary to observed changes in glycolic metabolism in the ATG5 cKO cerebellum.**

**a,b,** Relative profile of TCA cycle intermediates in 1-month old (a) and 3-month-old (b) WT and ATG5 cKO cerebellum identified using targeted metabolomics approach. Each dot represents one animal (N=5 per genotype). Multiple T-tests with linear Benjamini, Krieger and Yekutieli correction. Descriptive statistics is available in source data material for Fig. S4.

**c,** Extracellular acidification rates (ECAR), produced by the flux of H^+^ upon conversion of glucose to lactate in primary cerebellar cells (DIV17) from WT (N=5) and ATG5 cKO (N=6) mice cultured in 30 mM glucose during Seahorse XF glycolysis stress test with injections of 10 mM glucose, 1.5 µM oligomycin and 50 nM 2-DG.

**d**, Analysis of glycolytic capacity in the Seahorse XF glycolysis stress test performed with WT and ATG5cKO cultured cerebellar cells cultured in either 10 mM or 30 mM glucose. Each dot represents one well with ± 20 000 cells (from N=5 WT and N=6 ATG5 cKO mice). Two-way ANOVA followed by Holm-Šidák multiple comparisons (WT 10 mM: 0.68 ± 0.04; p<0.0001 in comparison to WT 30 mM, ATG5 cKO 10 mM: 1.05 ± 0.07; p=0.0007 in comparison to WT 10 mM, WT 30 mM: 1.292 ± 0.05; p=0.0019 in comparison to ATG5 cKO 30 mM, ATG5 cKO 30 mM: 1.53 ± 0.12; p<0.0001 in comparison to ATG5 cKO 10 mM).

**e,** Enrichment of ^13^C-glucose in Glucose-6-P M+6 fraction in acute cerebellar 3 month-old WT and ATG5 cKO slices (N=10 for WT, N=8 for ATG5 cKO mice).

**f,** Analysis of metabolic flux of glucose in TCA cycle intermediates in acute cerebellar 3 months-old WT and ATG5 cKO slices using ^13^C-glucose as a tracer (N=10 for WT, N=8 for ATG5 cKO). WT set to 1. One-tailed unpaired T-test (descriptive statistics can be found in source data for Fig. S4).

Data information: All graphs show mean ± SEM. n.s.—non-significant; * indicates P ≤ 0.05; ** indicates P ≤ 0.01; *** indicates P ≤ 0.001; **** indicates P ≤ 0.0001.

**Figure S5. No changes in pAMPK and/or GLUT1, GLUT3 and GLUT4 transporters in ATG5 cKO cerebellum.**

**a,** Analysis of 2-NBDG uptake efficiency in WT cerebellar organotypic slice culture at DIV 21. Each dot represents a single PC for 30 min 0.5 mM 18 cells (44.93 ± 4.04) for 45 min 0.5 mM 28 cells (71.96 ± 2.71), 60 min 0.5 mM 37 cells (58.53 ± 1.8) and 30 min 1 mM 23 cells (76.39 ± 5.64).

**b**, Area under the curve (AUC) analysis of Laconic signal (mVenus/eCFP ratio) in WT and ATG5 cKO cerebellar interneurons. Each dot represents a single PC (for WT 156 cells N=6 and for ATG5 cKO 128 cells from N=5). Two-tailed unpaired T-test (WT: 69.38 ± 0.65, ATG5 cKO: 67.78 ± 0.64).

**c,d**, Protein levels of AMPK and its phosphorylated form (pAMPKα-Thr172) are unaltered in cerebellar brain lysates from 3-month-old ATG5 cKO mice compared to the WT set to 1. N=4 for each genotype and condition. One-tailed unpaired t-test (ATG5 cKO: 1.13 ± 0.24).

**e,f,** Representative confocal images of WT and ATG5 cKO cerebellum at the age of 1 and 3 months expressing tdTomato as a reporter of Cre recombination, immunostained for GLUT1 (e) and analysis of its fluorescent levels compared to WT set to 1 (f). Scale bar: 50 µm. Each dot represents one animal (N=3 for 1-month old animals and N=5 for 3-months old animals per genotype). Two-way ANOVA followed by Holm-Šidák multiple comparisons (ATG5 cKO 1 month: 0.98 ± 0.21, ATG5 cKO 3 months: 0.76 ± 0.11).

**g,h,** Representative confocal images of WT and ATG5 cKO cerebellum at the age of 1 and 3 months expressing tdTomato as a reporter of Cre recombination, immunostained for GLUT3 (g) and analysis of its fluorescent levels compared to WT set to 1 (h). Scale bar: 50 µm. Each dot represents one animal (N=4 per genotype and condition). Two-way ANOVA followed by Holm-Šidák multiple comparisons (ATG5 cKO 1 month: 1.04 ± 0.08, ATG5 cKO 3 months: 1.16 ± 0.23).

**i,j,** Representative confocal images of WT and ATG5 cKO cerebellum at the age of 1 and 3 months expressing tdTomato as a reporter of Cre recombination, immunostained for GLUT4 (g) and analysis of its fluorescent levels compared to WT set to 1 (h). Scale bar: 50 µm. Each dot represents one animal (N=3 for 1-month old animals and N=4 for 3-months old animals per genotype). Two-way ANOVA followed by Holm-Šidák multiple comparisons (ATG5 cKO 1 month: 0.96 ± 0.1, ATG5 cKO 3 months: 1.07 ± 0.14).

**k,l,** Representative confocal images of WT and ATG5 cKO cerebellum at the age of 1 month expressing tdTomato as a reporter of Cre recombination, immunostained for hexokinase II (HK2) (k) and analysis of its fluorescent levels compared to WT set to 1 (l). Scale bar: 50 µm, inserts 10 µm. Each dot represents one animal (N=4 per genotype). One-tailed unpaired T-test (ATG5 cKO: 1.35 ± 0.15; p=0.03).

**m,** qPCR analysis of *Slc2a2* mRNA levels of WT and ATG5 cKO cerebellum at the age of 3 months. mRNA levels were normalized to the levels of housekeeping gene *Gapdh* and to WT levels and set to 1. Each dot represents one animal with n=4 per genotype. One-tailed unpaired T-test (WT: 1 ± 0.35, ATG5 cKO: 1.06 ± 0.29).

Data information: 1M and 3M indicate 1 and 3 months, respectively. m.g.v.-mean gray value. All graphs show mean ± SEM. n.s.—non-significant; * indicates P ≤ 0.05; ** indicates P ≤ 0.01; *** indicates P ≤ 0.001; **** indicates P ≤ 0.0001.

**Figure S6. Unaltered LC3 puncta density in cortical interneurons during maturation of the cortex**

**a,** Fluorescence-based analysis of GLUT2 protein levels in PC dendrites in 1- and 3-month old WT cerebellum. Each dot represents one animal (N=5 for 1-month old animals and N=3 for 3-month old animals). Two-tailed unpaired T-test (1 month: 34.47 ± 3.96, 3 months: 12.54 ± 2.47; p=0.007).

**b,c**, Immunoblot analysis of LC3 protein levels and its lipidation at autophagosomal membranes (LC3II/LC3I ratio) in control and chloroquine (CQ)- treated (6h) cerebellar and cortical acute slices of 3-month old WT animals. LC3 levels were normalized to control treated acute slices and set to 1. Each dot represents one animal (N=5 for cerebellum and N=3 for cortex). Two-way ANOVA followed by Holm-Šidák multiple comparisons (cerebellum CQ: 1.5 ± 0.1; p=0.0006, cortex CQ: 1.32 ± 0.14; p=0.029).

**d,e,** Representative confocal images (d) and fluorescence-based analysis of LC3 puncta density in cortical interneurons of 1- and 3-month old WT mice. WT mice were expressing tdTomato as a reporter of Cre recombination, immunostained with LC3. Scale bar, 50 µm. Each dot represents one animal (N=4 for 1 month, and N=3 for 3 months). Two-tailed unpaired T-test (1 month: 0.022 ± 0.005, 3 months: 0.024 ± 0.003).

Data information: Squares in Fig. S6d indicate regions magnified. All graphs show mean ± SEM. n.s.—non-significant; * indicates P ≤ 0.05; ** indicates P ≤ 0.01; *** indicates P ≤ 0.001; **** indicates P ≤ 0.0001.

**Figure S7**. **Levels of methylglyoxal-modified proteins are not altered in cerebellar interneurons.**

**a,** Representative confocal images of immunostained methylglyoxal(MG)-modified proteins in WT and ATG5 cKO PCs at the age of 1 month. WT and ATG5 cKO mice were expressing tdTomato as a reporter of Cre recombination. Scale bar, 50µm, inserts 10µm.

**b,** Immunofluorescence analysis of levels of methylglyoxal(MG)-modified proteins in WT and ATG5 cKO PC dendrites at the age of 1 and 3 months. Each dot represents one animal (N=4 for 1-month old animals and N=5 for 3-month old animals). MG-modified protein levels were normalized to WT and set to 1. Two-way ANOVA followed by Holm-Šidák multiple comparisons (ATG5 cKO 1 month: 1.09 ± 0.075, ATG5 cKO 3 months: 1.33 ± 0.085; p=0.0018).

**c,** Immunofluorescence analysis of levels of methylglyoxal(MG)-modified proteins in WT and ATG5 cKO cerebellar interneurons at the age of 1 and 3 months. Each dot represents one animal (N=4 for 1-month old animals and N=5 for 3-month old animals). MG-modified protein levels were normalized to WT and set to 1. Two-way ANOVA followed by Holm-Šidák multiple comparisons (ATG5 cKO 1 month: 1.14 ± 0.037, ATG5 cKO 3 months: 1.14 ± 0.092).

**d,e**, Representative confocal images (d) and immunofluorescence analysis (e) of cleaved caspase 3 levels in 3-month-old WT and ATG5 cKO mice. WT and ATG5 cKO mice were expressing tdTomato as a reporter of Cre recombination. Scale bar 200µm. Each dot represents one animal (N=3 per genotype). Cleaved caspase 3 protein levels were normalized to WT and set to 1. Two-way ANOVA followed by Holm-Šidák multiple comparisons (PC: 0.96 ± 0.14, cerebellar interneurons: 0.97 ± 0.164, cortical interneurons: 0.93 ± 0.04).

**f,g**, GCAMP7f baseline fluorescence (f) and its area under the curve analysis (AUC, g) in untreated PCs, LPA-treated WT PCs and ATG5 cKO PCs. Each dot in (g) represents a single PC (N=3 per genotype with 159 cells for WT, 79 for WT+LPA and 117 for ATG5 cKO). One-way ANOVA followed by Holm-Šidák multiple comparisons (WT: 2334 ± 178.8, WT+LPA: 2055 ± 194, ATG5 cKO: 2378 ± 193.8).

Data information: 1M and 3M indicate 1 and 3 months, respectively. Squares in Fig. S7a indicate regions magnified. All graphs show mean ± SEM. n.s.—non-significant; * indicates P ≤ 0.05; ** indicates P ≤ 0.01; *** indicates P ≤ 0.001; **** indicates P ≤ 0.0001.

**Figure S8. Genotype and phenotype of ATG5 cKO:GLUT2 cKO mice.**

**a,** Mice were screened by PCR analysis of tail DNA, using the primer pairs indicated in Table S1, which generates 350 and 700 bp products from the floxed *Atg5* and wild-type alleles, and/or 235 and 326 bp products from the floxed *Slc2a2* and wild-type alleles, respectively. Double-foxed mice for *Atg5* and *Slc2a2* with additional hemizygosity for the Cre gene (detected by the unique 326 bp product amplified in the Cre -PCR reaction, in addition to the 235 bp internal positive control) were defined as ATG5 cKO:GLUT2 cKO mice.

**b,** Bodyweight chart of WT, ATG5 cKO, GLUT2 cKO and ATG5 cKO:GLUT2 cKO mice.

**c**, Bar histogram showing bodyweight of 8-week-old WT, ATG5 cKO, GLUT2 cKO and ATG5:GLUT2 cKO mice. Each dot represents a single mouse (N=6 for WT, N=7 for ATG5 cKO, N=5 for GLUT2 cKO and N=7 for ATG5:GLUT2 cKO mice). One-way ANOVA followed by Holm-Šidák multiple comparisons (WT: 19.87 ± 0.82, ATG5 cKO: 14.84 ± 0.66; p=0.014 in comparison to WT, GLUT2 cKO: 21.18 ± 1.88; p=0.003 in comparison to ATG5 cKO, ATG5:GLUT2 cKO: 17.23 ± 1).

**d**, Representative confocal images of GLUT2 levels in WT and GLUT2 cKO PCs at the age of 3 months. Scale bar, 100µm.

**e,** Representative confocal images of GLUT2 levels in WT and ATG5:GLUT2 cKO PCs at the age of 3 months. Scale bar, 100µm.

**f,** Immunofluorescence analysis (m.g.v. after background subtraction) of GLUT2 proteins levels in s in WT, GLUT2 cKO and ATG5:GLUT2 cKO PCs (WT 28.03±0.633, GLUT2 cKO 11.25±0.405, p<0.0001, GLUT2 cKO 8.309 ± 0.235, p<0.0001, one-way ANOVA followed by Holm-Šidák multiple comparisons). Each dot represents single cells from N=3 WT, N=1 GLUT2 cKO and N=3 ATG5:GLUT2 cKO.

**g,** Immunoblotting analysis of protein levels of GLUT2 in cerebellar brain lysates from 3-month-old WT, ATG5 cKO and ATG5:GLUT2cKO mice.

**h**, Analysis of 2-NBDG uptake in WT and GLUT2cKO PCs in cerebellar organotypic slice culture at DIV 21. Each dot represents a single PC (for WT 62 cells from N=4 and for GLUT2 cKO 49 cells from N=4). Two-tailed unpaired T-test (WT: 74.62 ± 3.8, GLUT2 cKO: 86.9 ± 5.36).

**i,** Bar diagram of the time spent by WT and ATG5cKO mice on the rotarod at the age of 3 months. Each dot represents a single animal with N=11 for WT and N=12 for ATG5 cKO. Two-tailed unpaired T-test (WT: 190.7 ± 23.64, ATG5 cKO: 167.5 ± 22.93).

**j**, Bar diagram showing the average number of slips as 12-month-old WT and ATG5 cKO mice when crossing beams of 25mm and 12 mm widths. Each dot represents a single animal with n=5 for WT and N=7 for ATG5 cKO. Two-way ANOVA followed by Holm-Šidák multiple comparisons (WT 25 mm: 0 ± 0, ATG5 cKO 25 mm: 0.93 ± 0.24, WT 12mm: 0.067 ± 0.067, ATG5 cKO 12 mm (3.59 ± 0.83; p=0.0002).

Data information: m.g.v.-mean gray value. Squares in Fig. S8d,e indicate regions magnified. All graphs show mean ± SEM. n.s.—non-significant; * indicates P ≤ 0.05; ** indicates P ≤ 0.01; *** indicates P ≤ 0.001; **** indicates P ≤ 0.0001.

**Figure S9. Kinematic analysis of motor coordination in 3-month-old WT, ATG5 cKO and ATG5;GLUT2 cKO mice**

**a**, Individual frames from videos showing WT, ATG5 cKO and ATG5;GLUT2 cKO mice walking on the narrow beam (25mm).

**b**, Line graphs showing the variation in hip, knee and ankle angles during a normalized step cycle in WT, ATG5cKO and ATG5:GLUT2 cKO mice walking on a 25mm-wide beam. Red arrowheads indicate the points in the step cycle at which the relative positions of the joint differ in ATG5 cKOs compared to controls and ATG5:GLUT2 cKOs. N=9 mice WT and ATG5 cKO, N=6 for ATG5:GLUT2 cKOs.

**c**, Line graphs showing the variation in knee and hip angles during a normalized step cycle as mice walked on a 25mm-wide beam. N=9 mice for WT and ATG5 cKOs, N=6 for ATG5:GLUT2 cKOs. Statistical significance was calculated with two-way ANOVA followed by Bonferroni’s post-hoc test.

**d**,**e,** Bar graphs showing the duration of stance (d) and swing (e) as mice cross beams of different widths. Individual mice are depicted as grey filled circle. Statistical significance was calculated with one-way ANOVA followed by Tukey’s post-hoc test.

**f,i,l,** Line graphs showing the coordinates of the iliac crest, hip, knee, ankle and hindpaw on the y-axis during a normalized step cycle as mice walked on a 12mm-wide beam. Red arrowheads in (i) indicate the points in the step cycle at which the relative positions of the join differ in ATG5 cKOs compared to controls and ATG5:GLUT2 cKOs. N=8 mice for controls, N=11 mice for ATG5 cKOs, N=6 for ATG5:GLUT2 cKOs.

**g,j,m**, Line graphs showing the variation in hip, knee and ankle angles during a normalized step cycle as mice walked on a 12mm-wide beam. Red arrowheads in (j) indicate the points in the step cycle at which the relative values of the angle differ in ATG5 cKOs compared to controls and ATG5:GLUT2 cKOs. N=8 mice for controls, N=11 mice for ATG5 cKOs, N=6 for ATG5:GLUT2 cKOs.

**h,k,n** Line graphs showing the variation in hip, knee, ankle and hindpaw velocities during a normalized step cycle as mice walked on a 12mm-wide beam. Red arrowheads in (k) indicate the points in the step cycle at which the relative positions of the join differ in ATG5 cKOs compared to controls and ATG5:GLUT2 cKOs. N=8 mice for controls, N=11 mice for ATG5 cKOs, N=6 for ATG5:GLUT2 cKOs.

**o**, PCA analysis based on the y-coordinates, angles, and velocities per each normalized point (n=25) of the step cycle as mice walked on a 12mm-wide beam. Individual mice are represented as colored circles, and dashed lines group mice of similar genotype.

**p,s,v,** Line graphs showing the coordinates of the iliac crest, hip, knee, ankle and hindpaw on the y-axis during a normalized step cycle as mice walked on a 5mm-wide beam. N=10 mice for controls, N=3 mice for ATG5 cKOs, N=6 for ATG5:GLUT2 cKOs.

**q,t,w**, Line graphs showing the variation in hip, knee and ankle angles during a normalized step cycle as mice walked on a 5mm-wide beam. Red arrowheads in (t) indicate the points in the step cycle at which the relative values of the angle differ in ATG5 cKOs compared to controls and ATG5:GLUT2 cKOs. N=10 mice for controls, N=3 mice for ATG5 cKOs, N=6 for ATG5;GLUT2 cKOs.

**r,u,x,** Line graphs showing the variation in hip, knee, ankle and hindpaw velocities during a normalized step cycle as mice walked on a 5mm-wide beam. Red arrowheads in (u) indicate the points in the step cycle at which the relative positions of the join differ in ATG5 cKOs compared to controls and ATG5:GLUT2 cKOs. N=10 mice for controls, N=3 mice for ATG5 cKOs, N=6 for ATG5:GLUT2 cKOs.

**y**, PCA analysis based on the y-coordinates, angles, and velocities per each normalized point (n=25) of the step cycle as mice walked on a 5mm-wide beam. Individual mice are represented as colored circles, and dashed lines group mice of similar genotype.

Data information: All graphs show mean ± SEM. Data presented in Fig. S9b-x show mean as a filled dark line, and SEM as the shaded area around it. n.s.—non-significant; * indicates P ≤ 0.05; ** indicates P ≤ 0.01; *** indicates P ≤ 0.001; **** indicates P ≤ 0.0001.

**Appendix Table S1:** Primers used for genotyping of genetically modified mice used in the current study.

| **Gene** | **Sequence (5’ - 3’)** |
| --- | --- |
| *Glut2*  7S  7AS  7AS Δ | CCA ATC CCT TGG TTC ATG GTT GC  CGT AAG GCC CAA GGA AGT CCT GC  CTG CTA AAG CGC ATG CTC CAG AC |
| *Atg5*  forward1  forward2  reverse | GAA TAT GAA GCC ACA CCC CTG AAA TG  ACA ACG TCG AGC ACG CTG GCG AAG G  GTA CTG CAT AAT GGT TTA ACT CTT GC |
| *Cre*  Cre_1  Cre_2  Cre_3  Cre_4 | GAA CCT GAT GGA CAT GTT CAG G  AGT GCG TTC GAA CGC TAG AGC CTG T  TTA CGT CCA TCG TGG ACA  TGG GCT GGG TGT TAG CC |
| *tdTomato*  tdTomato_1  tdTomato_2  tdTomato_4  tdTomato_4 | AAG GGA GCT GCA GTG GAG TA  CCG AAA ATC TGT GGG AAG TC  GGC ATT AAA GCA GCG TAT CC  CTG TTC CTG TAC GGC ATG G |

**Appendix Table S2:** AAVs used for transduction of neurons e-vivo and in-vivo in the current study.

| **Identifier** | **Name** | **Titer [vg/ml]** | **Origin** |
| --- | --- | --- | --- |
| ATeam | ssAAV-9/2-hSyn1-Ateam1.03YEMK-WPRE-hGHp(A) | 8.1x10E12 | Viral Vector Facility (VVF) |
| Laconic | ssAAV-9/2-hSyn1-Laconic-WPRE-hGHp(A) | 6.9x10E12 | Viral Vector Facility (VVF) |
| GCamP7f | ssAAV-9/2-mCaMKIIα-jGCaMP7f-WPRE-bGHp(A) | 1.1x10E13 | Viral Vector Facility (VVF) |
| MitoTimer | ssAAV-5/2-mTH-dlox-MitoTimer(rev)-dlox-WPRE-bGHp(A) | 6.2 x 10E12 | Viral Vector Facility VVF |
| EGFP-ATG5 | pAAV2/RH10-mDLX-EGFP-ATG5 | 4.78 x 10E12 | Custom-made |
| EGFP | pAAV2/RH10-mDLX-EGFP | 1.31 x 10E12 | Custom-made |
| mt-Keima | AAV9S-Syn1-mt-mKeima |  | Vector Builder |

Appendix Table S3: List of primary and secondary antibodies used in immunohistochemical studies.

| **Antibody target** | **Concentration** | | **Manufacturer** | | **Catalog number** |
| --- | --- | --- | --- | --- | --- |
| AKT | 1:500 | | Cell Signalling | | 46855 |
| Calbindin | 1.500 | | Novus Biologicals | | NBP2-50028 |
| Cleaved Caspase-3 | 1:5000 | | Cell Signalling | | 9661S |
| Extracellular GLUT2 | 1:300 | | Merck | | 07-1402-I |
| GFAP | 1:500 | | Sigma | | G3893 |
| GFP | 1:1000 | | Abcam | | ab13970 |
| GLUT1 | 1:300 | | Novus Biologicals | | NB110-39113SS |
| GLUT2 | 1:300 | | Novus Biologicals | | NBP2-22218SS |
| GLUT3 | 1:300 | | Invitrogen | | OSG00012w |
| GLUT4 | 1:300 | | Novus Biologicals | | NBP1-4953355 |
| Hexokinase II | 1:500 | | Abcam | | ab227198 |
| LC3 | 1:500 | | Biozol | | M152-3 |
| mCathepsin D | 1:300 | | R&D systems | | AF1029 |
| Methylglyoxal | 1.300 | | Novus Biologicals | | NBP2-59368 |
| NBR1 | 1:300 | | Santa Cruz Biotechnology | | sc-130380 |
| Parvalbumin | 1:500 | | SySy | | 195002 |
| p62 | 1:1000 | | Progen | | GP62-C |
| Ubiquitin | 1:300 | | Sigma | | U5379 |
| Alexa Fluor 488 Goat Anti-Chicken IgG | 1:500 | Life Technologies GmbH | | A11039 | |
| Alexa Fluor 488 Goat Anti-Rabbit IgG | 1:500 | Life Technologies GmbH | | A11034 | |
| Alexa Fluor 488 Goat Anti-Guinea Pig IgG | 1:500 | Life Technologies GmbH | | A11073 | |
| Alexa Fluor 488 Goat Anti-Mouse IgG | 1:500 | Life Technologies GmbH | | A11029 | |
| Alexa Fluor™ 488 Donkey Anti-Mouse | 1:500 | Life Technologies GmbH | | A32766 | |
| Alexa Fluor 568 donkey anti-goat IgG Life | 1:500 | Life Technologies GmbH | | A10057 | |
| Alexa Fluor 568 goat anti-rabbit IgG | 1:500 | Life Technologies GmbH | | A11011 | |
| Alexa Fluor 647 Donkey Anti-Rabbit IgG | 1:500 | Life Technologies GmbH | | A31573 | |
| Alexa Fluor 647 Goat Anti-Guinea Pig IgG | 1:500 | Life Technologies GmbH | | A21450 | |
| Alexa Fluor 647 Goat Anti-Rabbit IgG | 1:500 | Life Technologies GmbH | | A21245 | |
| Alexa Fluor 647 Goat Anti-Mouse IgG | 1:500 | Life Technologies GmbH | | A21236 | |

Appendix Table S4: List of primary and secondary antibodies used in immunoblotting studies.

| Antibody target | Dilution | Manufacturer | | Catalog number |
| --- | --- | --- | --- | --- |
| ATG5 | 1:1000 | Abcam | | ab108327 |
| α-Tubulin | 1:5000 | Synaptic Systems | | 302 211 |
| β-Actin | 1:3000 | Sigma | | A-5441 |
| Calbindin | 1:1000 | Novus Biologicals | | NBP2-50028 |
| Cleaved-Caspase 3 | 1:500 | Cell Signalling | | 9661S |
| GFAP | 1:1000 | Sigma | | G3893 |
| GLUT1 | 1:1000 | Novus Biologicals | | NB110-39113SS |
| GLUT2 | 1:500 | Novus Biologicals | | NBP2-22218SS |
| GLUT2 | 1:500 | Proteintech | | 20436-1-AP |
| GLUT3 | 1:500 | Invitrogen | | OSG00012w |
| GLUT4 | 1:500 | Novus Biologicals | | NBP1-4953355 |
| LC3B | 1:1000 | Novus Biologicals | | NB600-1384 |
| Methylglyoxal | 1:1000 | Novus Biologicals | | NBP2-59368 |
| NBR1 | 1:500 | Santa Cruz Biotechnology | | sc-130380 |
| p62 | 1:1000 | Progen | | GP62-C |
| Vinculin | 1:5000 | Abcam | | ab129002 |
| Rabbit anti-Mouse IgG (H+L) peroxidase-conjugated | 1:10 000 | | Sigma | A9044 |
| Goat anti-Rabbit IgG (H+L) peroxidase-conjugated | 1:10 000 | | Sigma | A0545 |
| Goat anti-Guinea Pig IgG (H+L) peroxidase-conjugated | 1:5 000 | | Jackson Immuno Research | 106-035-003 |
| Rabbit anti-Chicken IgG (H+L) peroxidase-conjugated | 1:5 000 | | Millipore | AP162P |

**Videos**

**Video EV1**

Kinematic analysis of WT, ATG5 cKO and ATG5:GLUT2 cKO mice while crossing 5 mm beam.

**Video EV2**

Kinematic analysis of WT, ATG5 cKO and ATG5:GLUT2 cKO mice while crossing 12 mm beam.

**Video EV3**

Kinematic analysis of WT, ATG5 cKO and ATG5:GLUT2 cKO mice while crossing 25 mm beam.
