## Supplementary figures and images for "Autophagy regulator ATG5 preserves cerebellar function by safeguarding its glycolytic activity"

### Supplementary Material 1-9

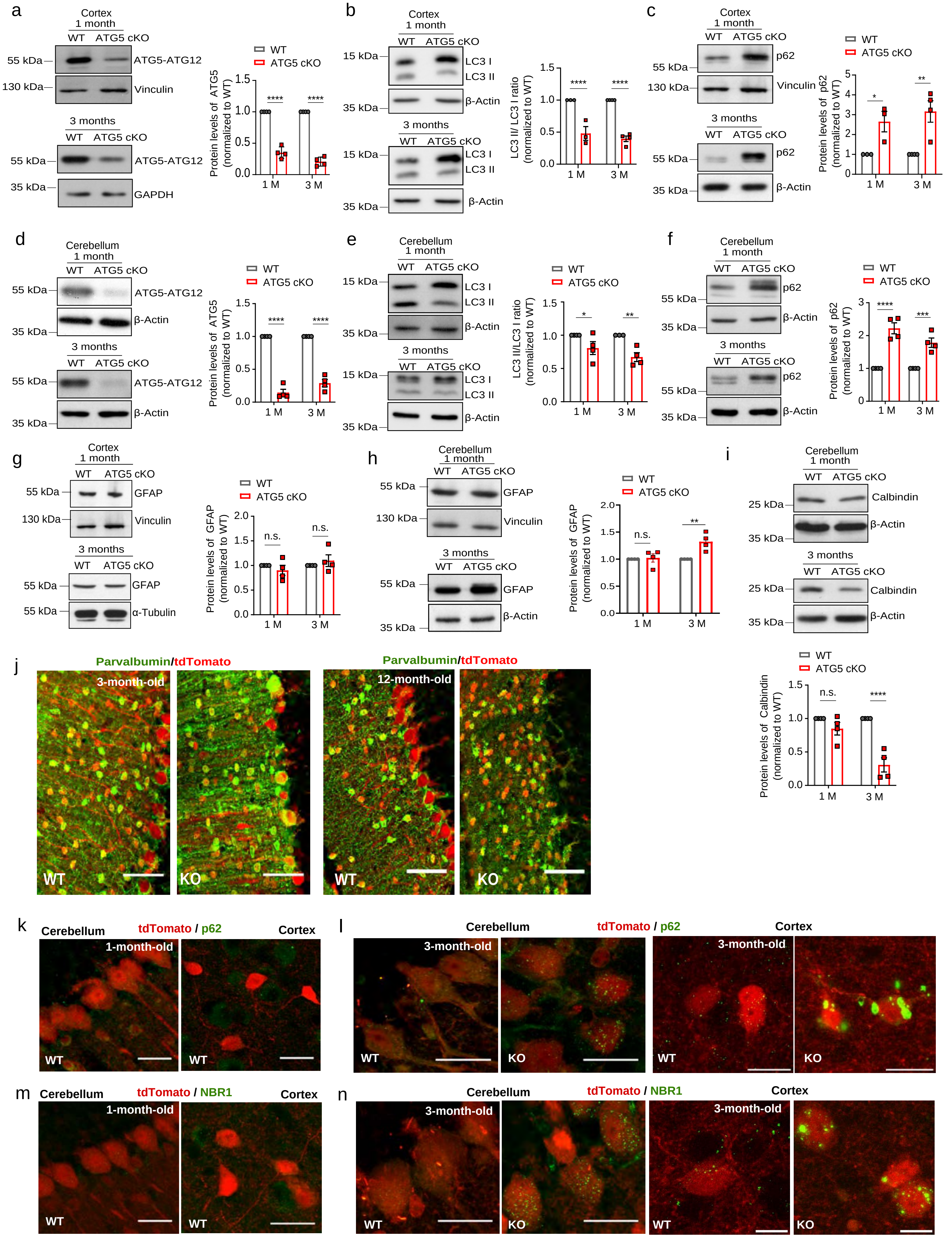

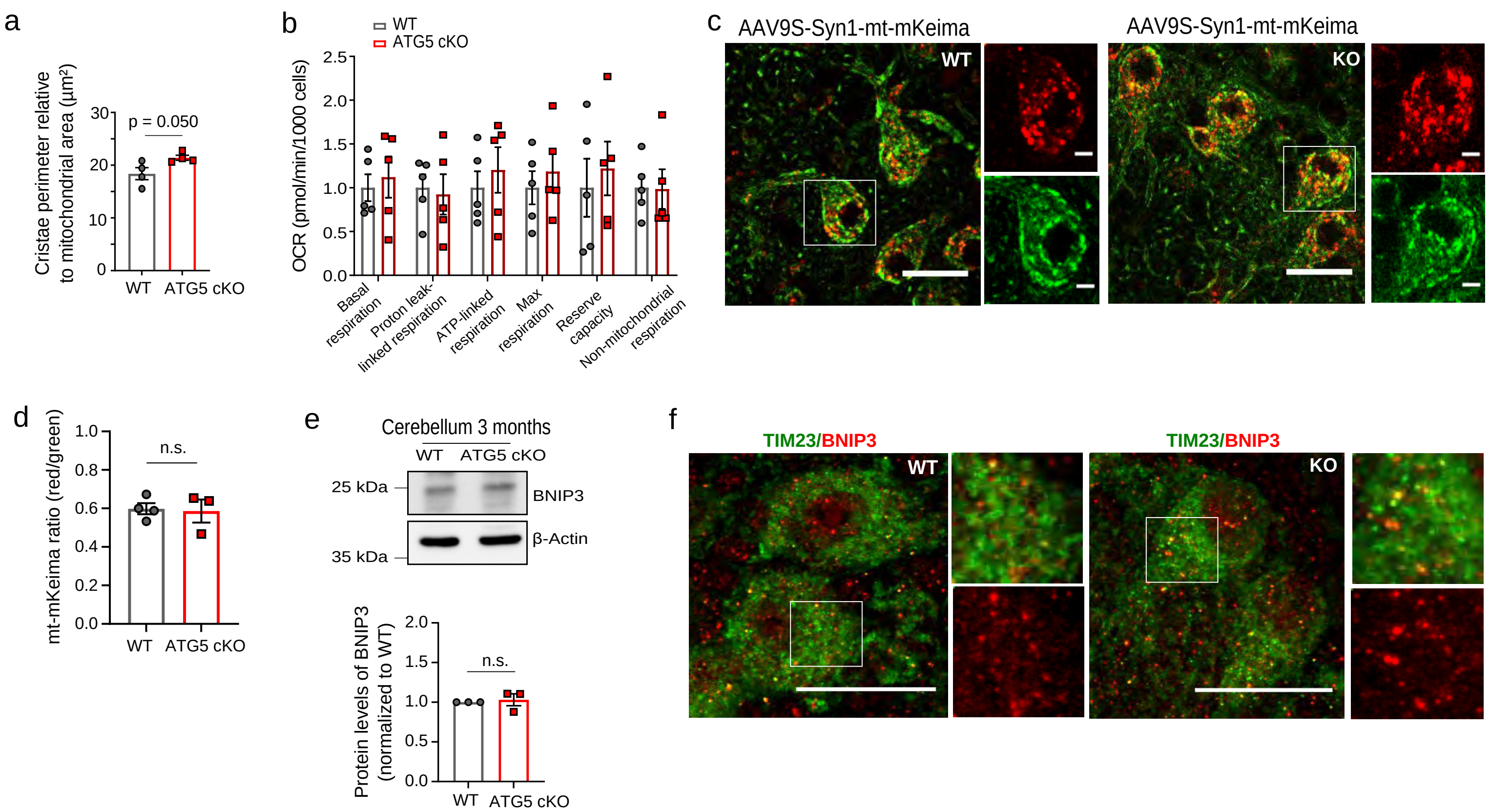

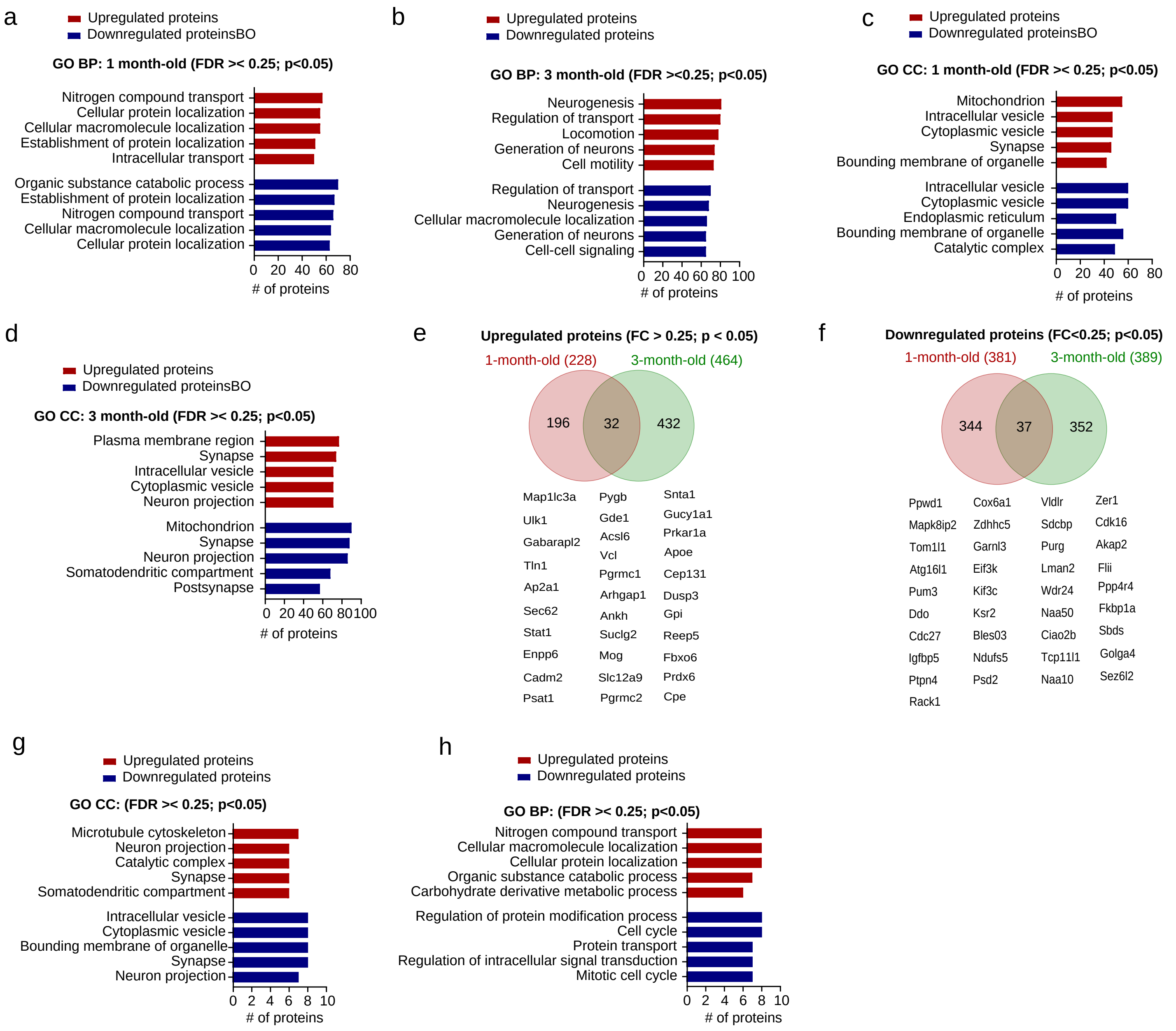

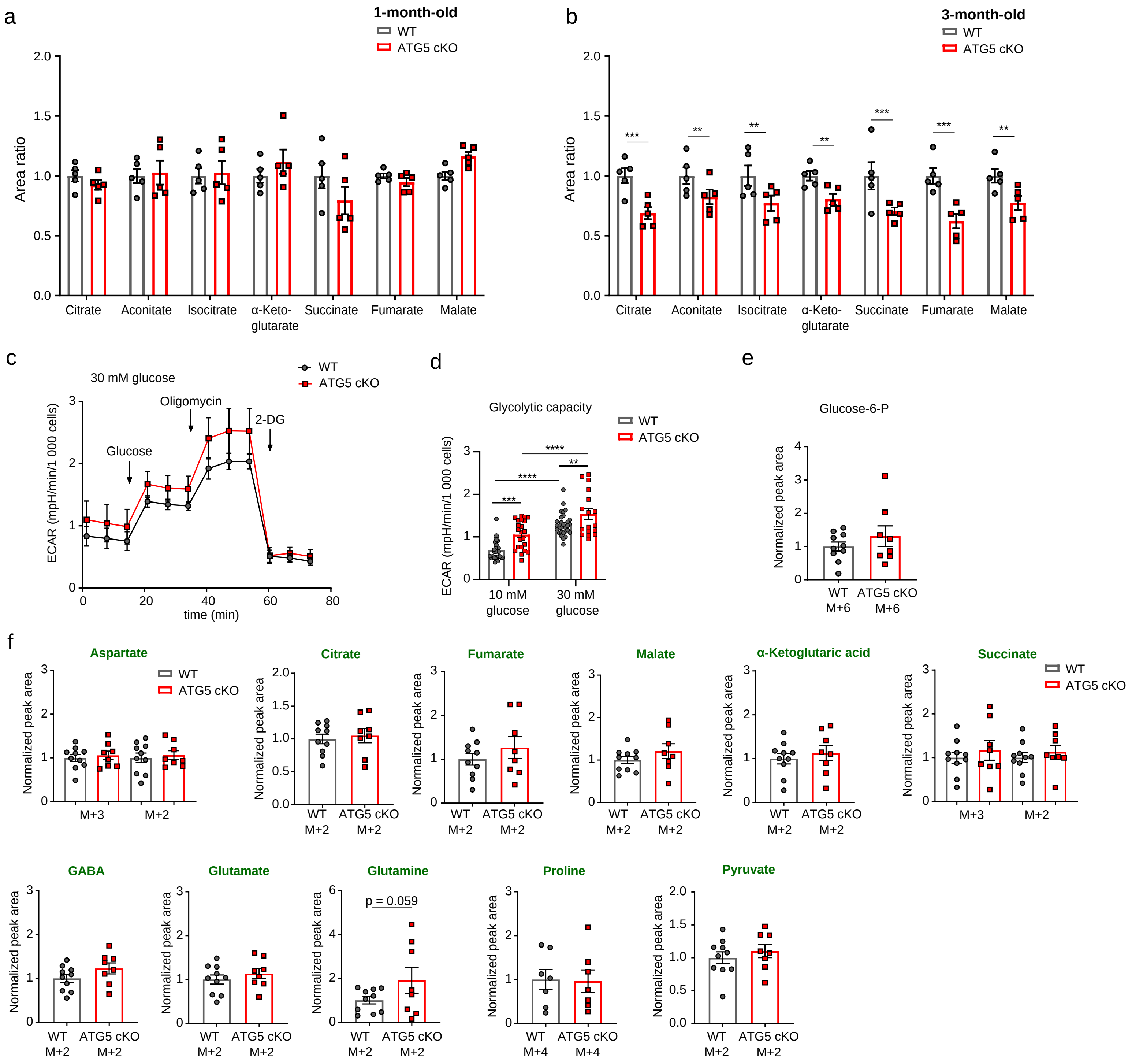

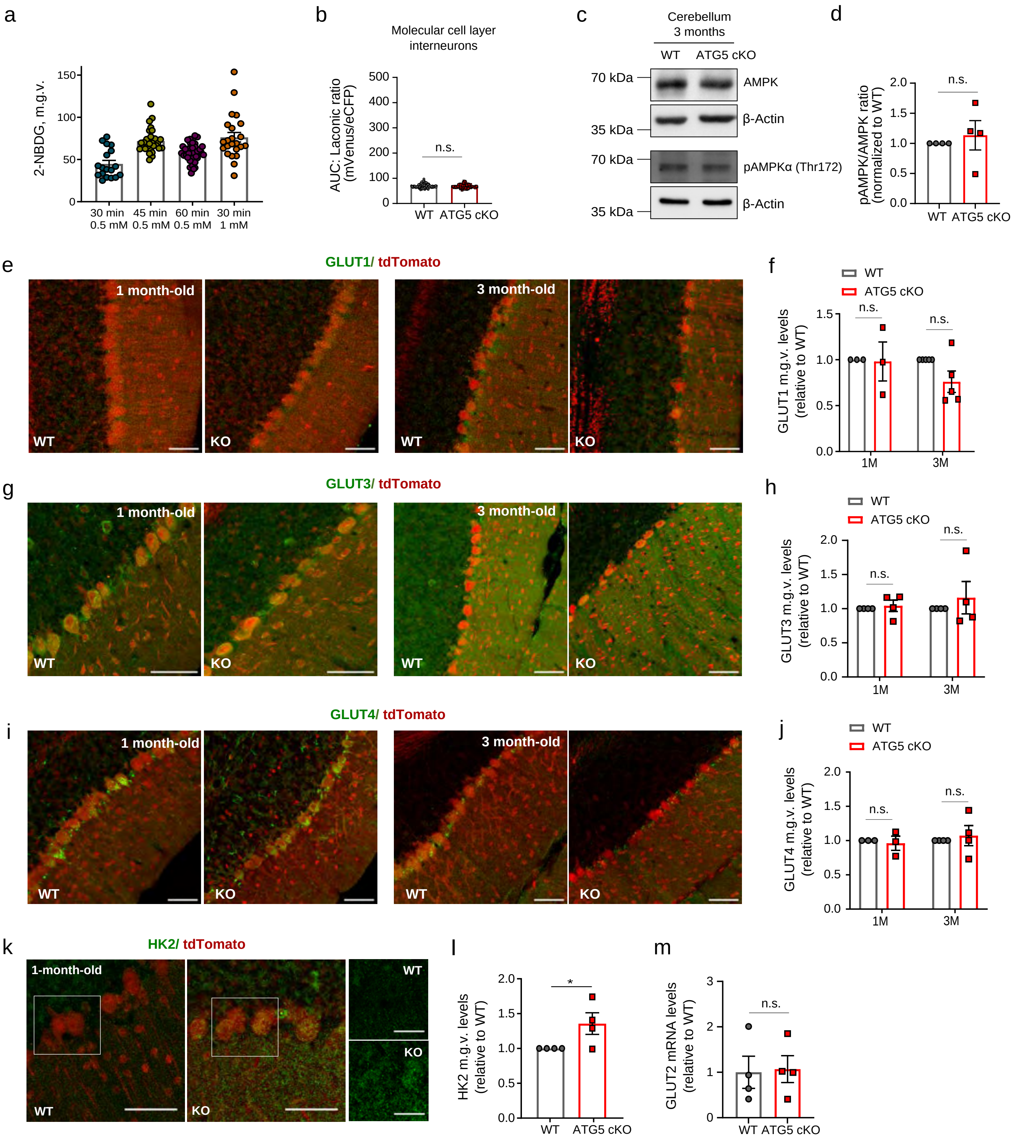

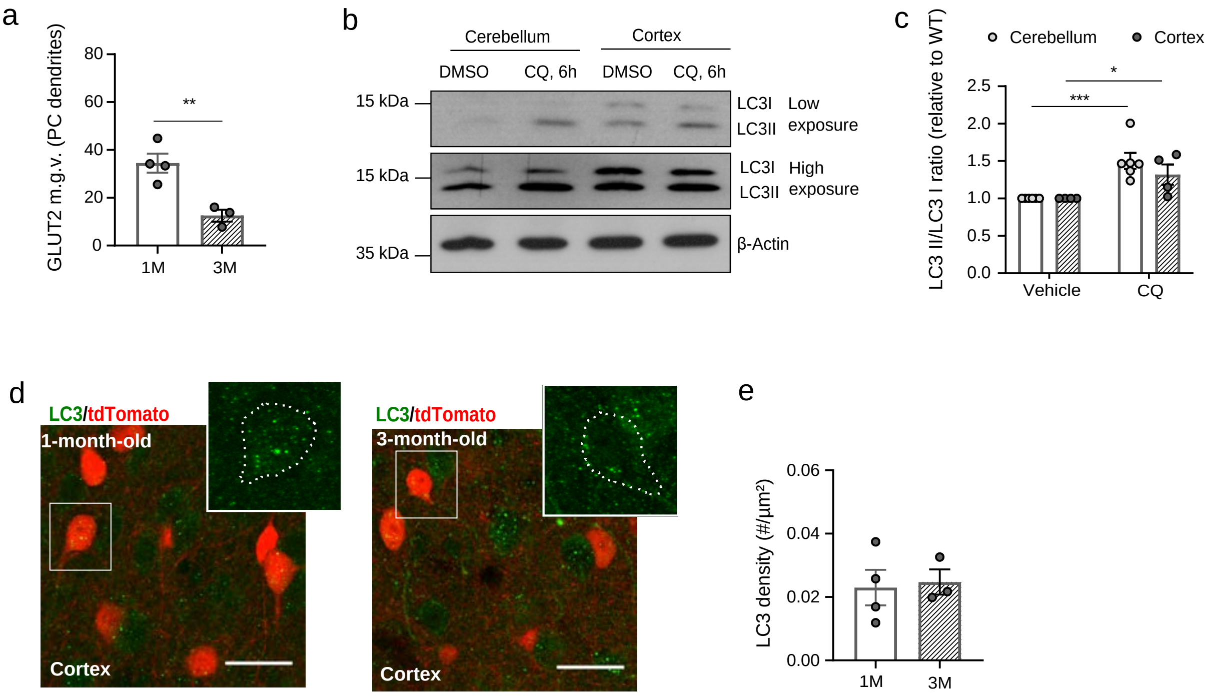

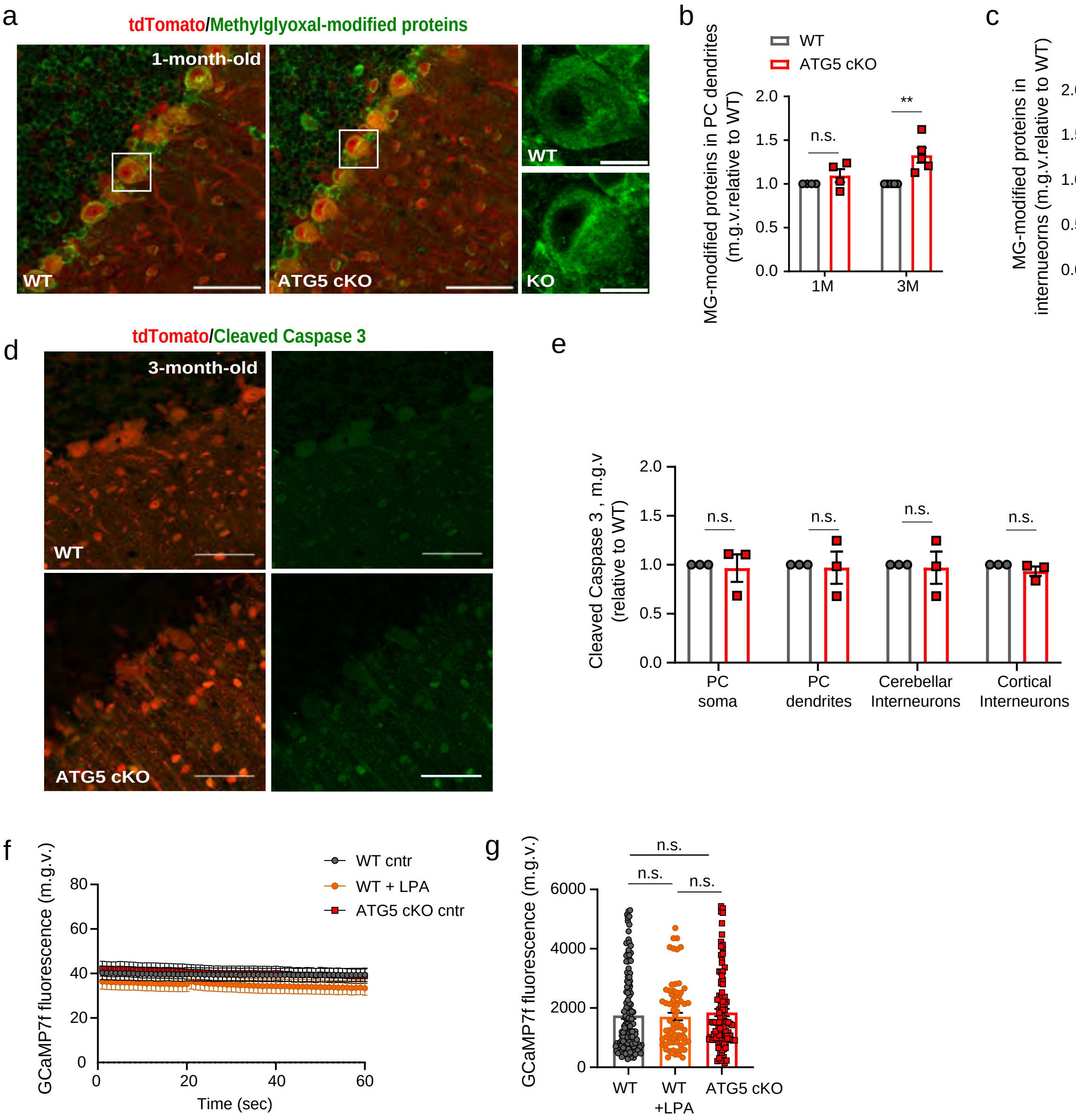

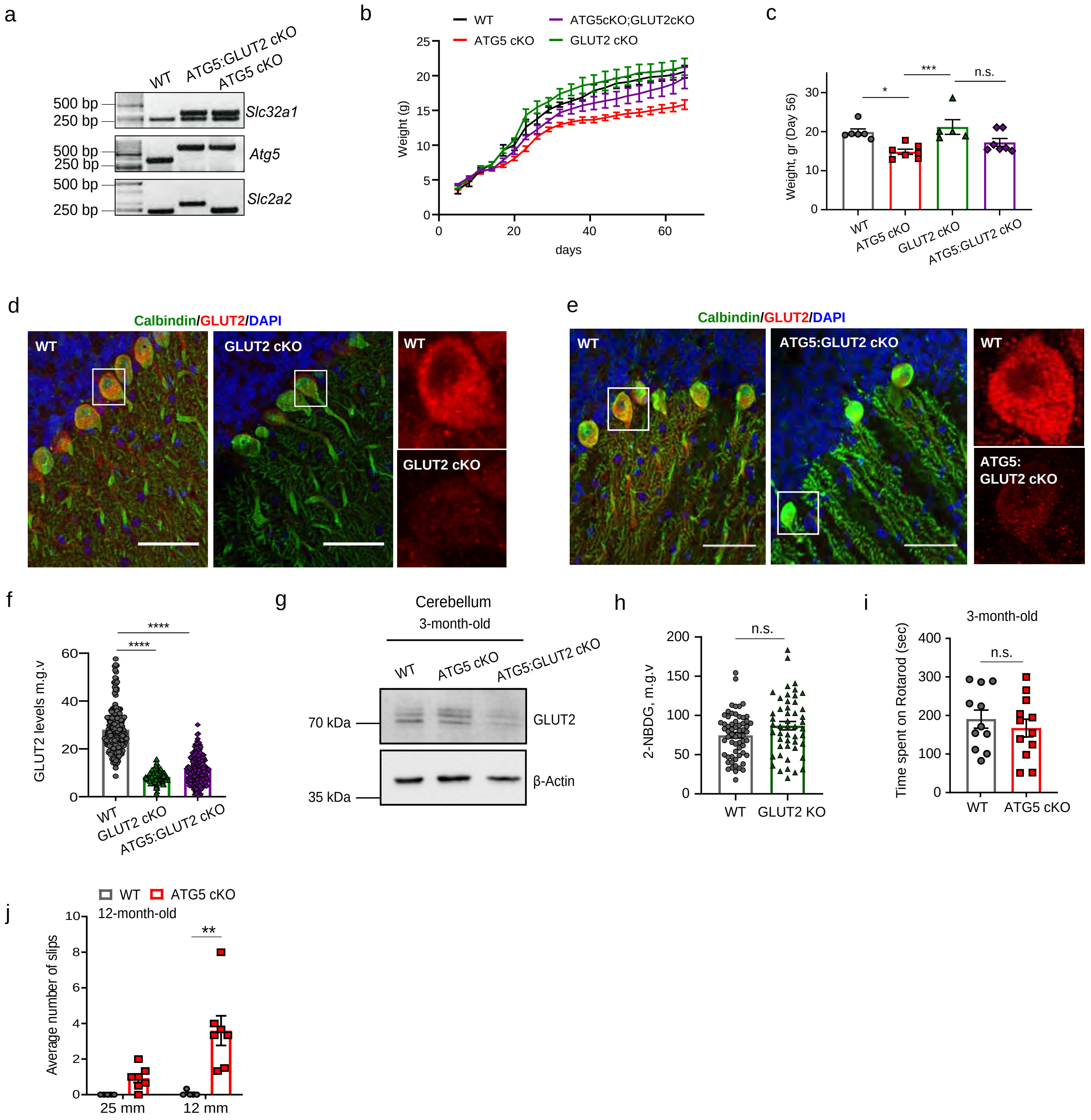

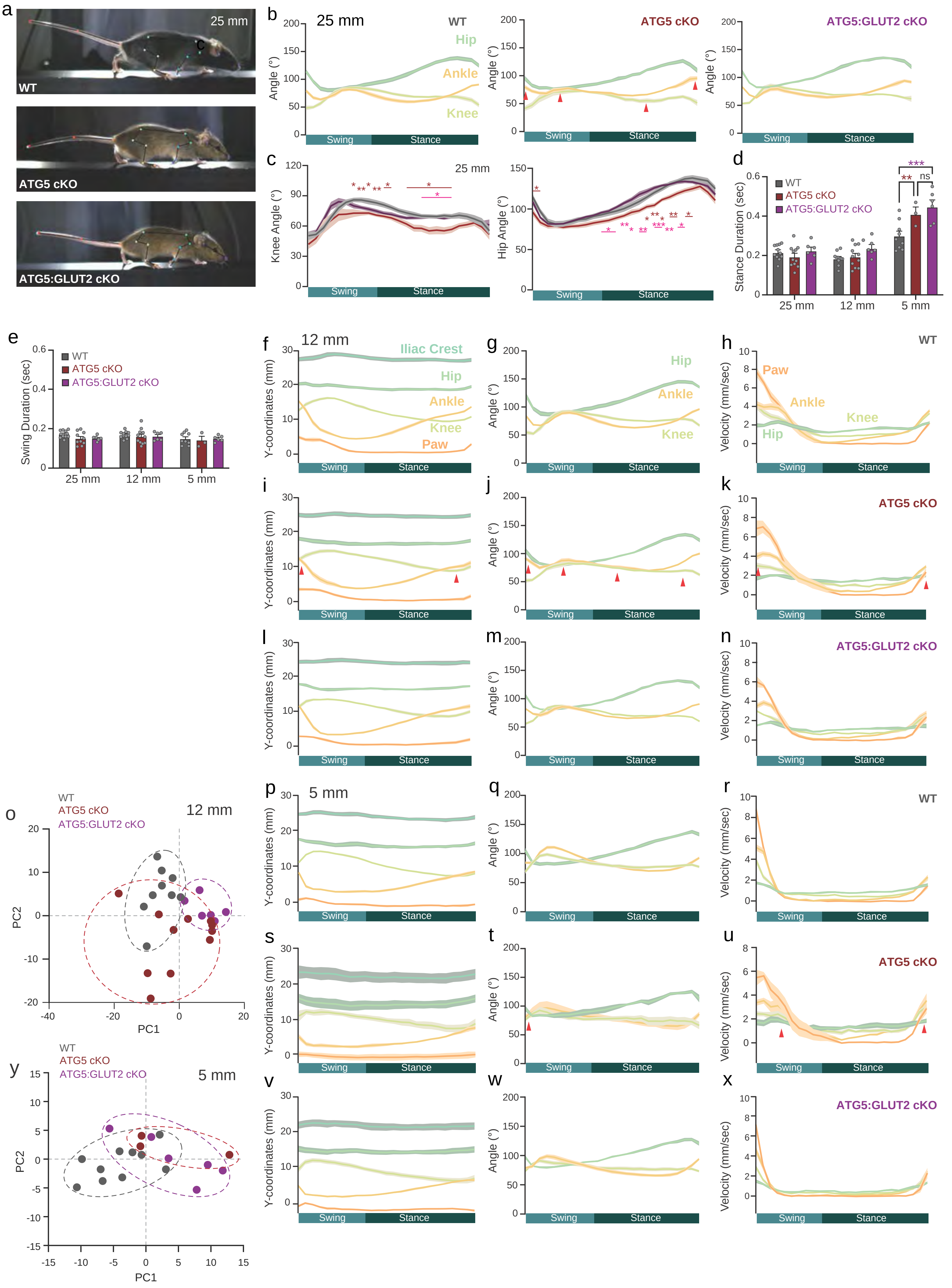
